# De novo Design of Polymorph-Specific Binders Targeting α-Synuclein Fibrils

**DOI:** 10.64898/2026.06.13.732066

**Authors:** Ahmed Sadek, Nolwen L. Rey, Antonin Kunka, Zander Harteveld, Sandrine Georgeon, Joseph Schmidt, Alexander K. Buell, Ronald Melki, Hilal A. Lashuel, Bruno E. Correia

## Abstract

Alpha-synuclein (aSyn) aggregation into amyloid fibrils is a hallmark of synucleinopathies, a group of neurodegenerative disorders that includes Parkinson’s disease. Beyond the central role of fibrils in disease pathology and propagation, distinct aSyn fibril polymorphs have been associated with different synucleinopathies, presenting opportunities for differential diagnosis and structure-based disease-specific phenotyping. Here, we present AmyBind, a computational pipeline for the *de novo* design of mini-protein binders that target polymorph-specific structural features of amyloid fibrils. Using this approach, we developed two aSyn fibril-specific binders, one of which exhibited polymorph-specificity. Neither binder interfered with fibril elongation *in vitro*, consistent with their designed binding mode on lateral fibril surfaces. In cell-based models, both binders showed polymorph-specific colocalization but did not affect fibril uptake or seeding, indicating their utility as fibril-selective labeling agents in cellular contexts. These findings demonstrate the feasibility of structure-guided, polymorph-specific amyloid targeting and provide a foundation for developing early-stage diagnostics and therapeutics for synucleinopathies and other amyloid-related diseases.

## Introduction

Neurodegenerative diseases (NDs) are a diverse group of neurological disorders characterized by progressive neuronal loss, leading to impairments in memory, cognition, and motor function^1^. A key hallmark of NDs is the misfolding and aggregation of soluble proteins, resulting in the formation of amyloid fibrils that accumulate in the form of intracellular inclusions and/or extracellular deposits^2^. Synucleinopathies, a subgroup of NDs that includes Parkinson’s disease (PD), dementia with Lewy bodies (DLB), and multiple system atrophy (MSA)^3^, are characterized by the aggregation of the protein α-synuclein (aSyn) and its accumulation in a predominantly fibrillar form within neuronal intracellular inclusions, such as Lewy bodies and Lewy neurites in PD, or glial cytoplasmic inclusions (GCIs) in MSA^4–6^.

Although the etiology of NDs remains unclear, increasing evidence point to fibril-mediated seeding, inclusion formation, and cell-to-cell propagation of protein aggregates, in a prion-like manner, as key drivers of pathology spreading in the brain, neurodegeneration, and disease progression^7–15^. Interfering with fibril-mediated seeding mechanisms further represents a promising therapeutic avenue to halt disease progression^16,17^. Given their early involvement and central role in pathogenesis^14,18^, aSyn fibrils represent promising targets for the development of early-stage, disease-modifying therapeutics. Recent structural studies of aSyn fibrils have revealed distinct polymorphs across synucleinopathies, thus presenting an opportunity for polymorph-specific targeting strategies that could enable disease differentiation and the development of fibril- and disease-specific diagnostics and therapeutics^19^.

Despite this potential of targeting aSyn fibrils, progress has been hampered by several challenges. To date, only six aSyn fibril structures isolated from patients’ brains have been solved, one shared by PD and DLB^20^, four from MSA^21,22^, and one from juvenile-onset synucleinopathy^23^, compared to sixteen tau fibril structures spanning eight tauopathies^19^. This limited structural repertoire constrains our understanding of aSyn polymorphism in synucleinopathies. Comparative analyses further show that aSyn fibrils assembled *in vitro* differ not only across preparation protocols but also from their pathological counterparts^19,24^. Coupled with the scarcity of patient-derived material and the difficulty of reproducing disease-relevant aSyn fibril folds *in vitro*^25–29^, these limitations have restricted the development of polymorph-selective, translationally viable molecules through conventional screening and optimization strategies^30–32^.

In response to these challenges, small-molecules, peptides and protein-based strategies have been explored to modulate aSyn aggregation, propagation, and polymorph differentiation, with varying degrees of conformational specificity and translational success. Several small molecules have been developed and reported as amyloid tracers for positron emission tomography (PET)^33^, enhancers of fibril disassembly^34–36^, or inhibitors of fibril seeding^37,38^ and prion-like transmission^39,40^. Despite promising *in vitro* findings, clinical translation remains challenging for most candidates due to factors, such as incomplete understanding of their mechanisms of action, poor blood-brain barrier permeability, toxicity, and limited target specificity^33,38,41–43^.

In parallel, protein engineering has emerged as a powerful strategy for developing site-specific binders against a wide range of molecular targets^44–48^, thus inspiring its application to identify novel fibril-binding proteins for diagnostic and therapeutic purposes. Towards this goal, several approaches have been explored, including grafting known fibril-binding motifs^49,50^ onto protein scaffolds to enhance stability and solubility^51^. These chimeric constructs exhibited selectivity for fibrillar over monomeric forms of wild-type (WT) aSyn and specificity for aSyn relative to Aβ fibrils implicated in Alzheimer’s disease^51,52^. However, the structural basis of their binding, the identity of their fibril-interacting sites, and their ability to distinguish WT from mutant aSyn fibrils, or to discriminate among WT aSyn fibril polymorphs, remain unresolved^51^.

The intrinsically disordered nature of monomeric amyloidogenic proteins has hindered the rational design of binders^53^. To address this challenge, a recent study used models of aggregation-prone sequences that adopt β-sheet conformations as structural templates to engineer peptide traps capable of sequestering the monomer and preventing its aggregation^54^. Despite effectively inhibiting aggregation, these traps lacked conformational specificity and also bound fibrillar forms^54^.

Leveraging structural insights to target aSyn fibrils^24^, several protein design strategies have been developed to engage solvent-accessible sections of the sequences inside the fibrils. These include fibril end-capping approaches that inhibit fibril seeding by blocking monomer addition^17,55^ and polymorph-differentiation strategies that exploit differences in the solvent accessibility of residues Q62–V66 among distinct aSyn polymorphs^56^. However, all have been shown to inhibit *de novo* aSyn aggregation *in vitro*, suggesting that they also bind to aSyn monomers or other intermediates on the pathway to fibril formation. This highlights a key limitation of targeting sequence motifs that are recognized in both monomeric and fibrillar conformations or that do not differentiate among different fibril polymorphs and underscores the potential of exploiting structural features unique to the fibrillar state to design conformation-specific binders^17,55,56^.

Here, we introduce AmyBind, a computational pipeline for the development of *de novo* amyloid-specific binders, utilizing protein design tools such as Rosetta^57^ and MaSIF (molecular surface interaction fingerprinting) framework^58^. AmyBind accounts for the unique structural features of the fibrils, including their helical symmetry and rise. Unlike existing amyloid-targeting strategies, our approach exploits structural differences among amyloid fibril polymorphs to enable the design of conformation- and polymorph-specific binders. This is achieved by targeting the grooves (i.e., sites) that run along the fibril main axis and arise from the distinct folding and packing of the protofilaments.

As a proof-of-concept, we targeted sites within two distinct polymorphs of recombinant WT aSyn fibrils formed *in vitro*, Polymorph 1a^59^ and Polymorph 2b^60^, representing the two major structural classes observed among aSyn fibrils^61^. Lead candidate binders were identified through yeast surface display screening against three well-characterized *in vitro* WT aSyn fibril preparations: Ribbons (Ri), Fibrils (Fm), and Fibrils-91 (F91)^62,63^. These preparations differ not only in morphology and Proteinase K digestion profiles but also in their membrane-binding properties and seeding capacities in disease-relevant neuronal models, underscoring their structural and biochemical divergence^15,63,64^.

These selected candidates were then expressed as GFP-fusion proteins, and their binding to different fibrils was assessed using flow-induced dispersion analysis (FIDA). To further assess their functional effects, we investigated their impact on aSyn aggregation *in vitro* using Thioflavin T (ThT) fluorescence assays and examined their influence on fibril uptake and seeding in HeLa and Neuro2A cells.

Using our newly developed pipeline, we identified two binders that selectively recognized aSyn fibrils, with no measurable binding to monomeric or oligomeric aSyn, or to fibrils formed by other amyloidogenic proteins. One binder further exhibited polymorph selectivity for the Ri fibril polymorph. Consistent with their designed engagement of lateral fibril surface sites, neither binder perturbed fibril elongation *in vitro* nor fibril-induced seeding in cellular models, while both retained polymorph-specific colocalization in cells. To our knowledge, these are the first experimentally validated, computationally designed *de novo* amyloid binders that exploit polymorph-specific structural sites on fibrils and exhibit target, conformational and polymorph selectivity without interfering with fibril growth and seeding activity. Together, these results position AmyBind as a framework for generating polymorph-selective binders and support its broader application for targeting other amyloid fibrils in the development of early-stage diagnostics and disease-modifying agents.

## Results

### AmyBind: a computational workflow to design binders targeting amyloid fibrils

Under diverse *in vitro* and pathological conditions, amyloid fibrils adopt distinct folds^19,24^, giving rise to polymorph-specific structural features that represent potential targets for conformation-specific binder design. Among these are conformation-specific grooves along fibril lateral surfaces, where stacks of monomers form sites with distinct structural features that are challenging to target both with small molecules and proteins. Unlike the deep pockets or extensive interfaces commonly exploited in molecular recognition, these sites form shallow, periodically repeating surfaces that offer few canonical interaction features, thereby limiting opportunities for high-affinity and selective binding^65–68^. To overcome these issues, we developed AmyBind, a computational pipeline for *de novo* design of protein binders to amyloid fibril sites, engaging the fibril surface in a previously unexplored mode, parallel to fibril axis (the direction of fibril growth), reminiscent of the chaperone-like BRICHOS domain binding to Aβ_42_ fibrils^69^ (**Fig. 1A**).

**Figure 1.**
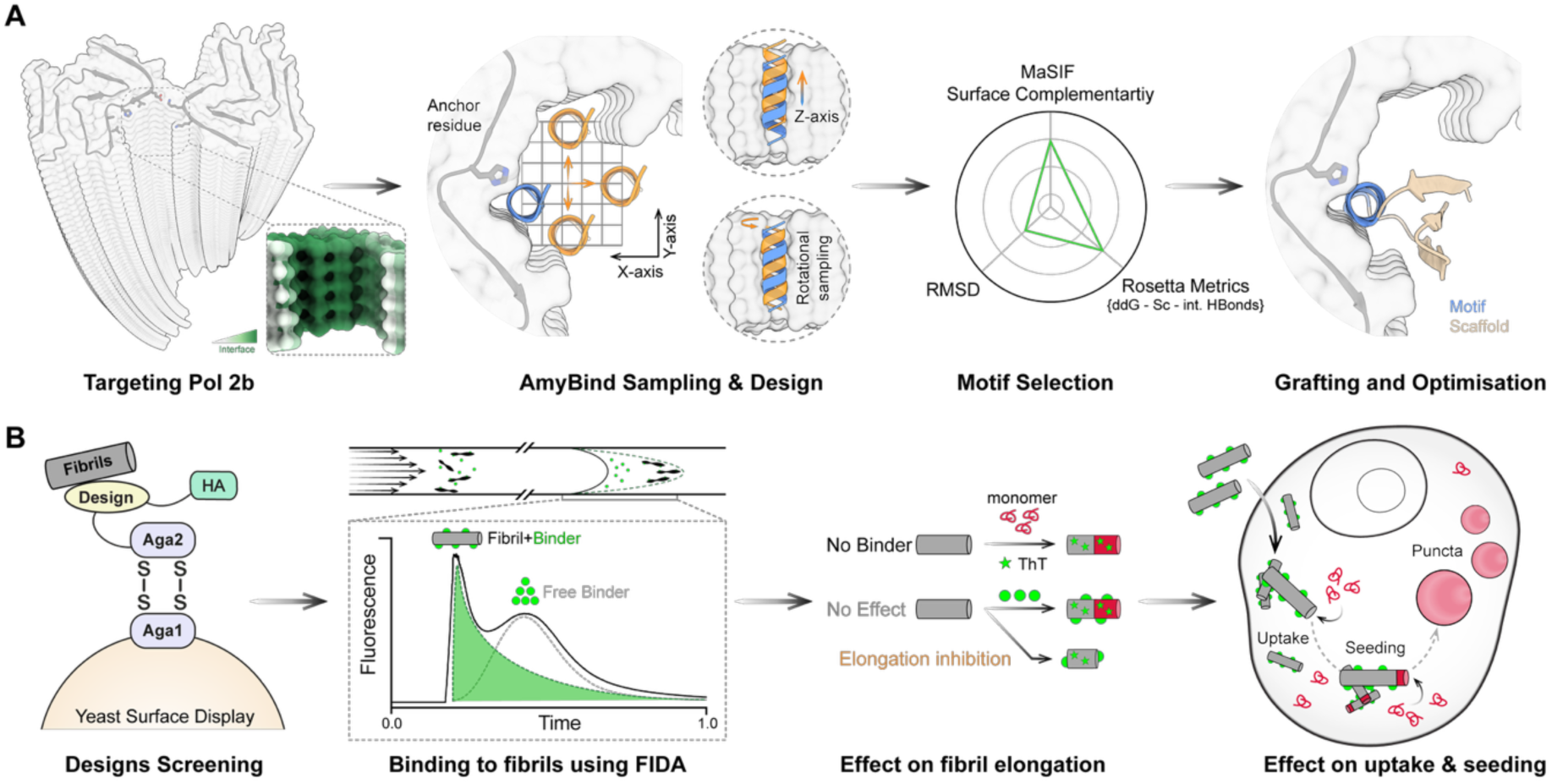
Computational design and experimental characterization of conformation-specific aSyn fibril binders. **(A)** Overview of the AmyBind computational workflow. The process starts with a structural model of the WT aSyn Polymorph 2b fibrils (PDB 6SST), with the targeted site residues shown as sticks and a close-up illustrating the repetitive structural features within the targeted site, including sub-site pockets on the lateral fibril surface. The site surface is colored according to MaSIF-site predictions of PPI potential (PPI sites in green and non-PPI sites in white). At the targeted site, a naïve helix (blue) is aligned along the fibril main axis relative to an anchor residue at the site surface. The docking site is then sampled by grid-based translations in X, Y, and Z planes, combined with rotational sampling around the Z-axis (orange). At each sampled position, helices (motifs) are designed in Rosetta and filtered based on Rosetta metrics, MaSIF fingerprint complementarity, and C_α_ RMSD alignment (≤2 Å) between MaSIF-docked and Rosetta models. Selected helices are subsequently grafted onto a mini-protein scaffold library and further optimized in Rosetta. **(B)** Experimental validation workflow starts with screening the designs using yeast surface display against different aSyn polymorphs. Lead candidates are then expressed as GFP-fusion proteins, and their binding profiles are assessed using FIDA. Effects on fibril elongation are evaluated *in vitro* using ThT binding in aggregation kinetics, while effects on fibril internalization (uptake) and seeding activity are assessed in PD-like cellular models.

Using AmyBind, we set out to design binders targeting two sites from distinct polymorphs of human WT aSyn fibrils formed *in vitro*: Polymorph 1a^59^ (Pol 1a) and Polymorph 2b^60^ (Pol 2b) (**Supplementary Fig. S1A, B**). MaSIF-site predictions indicated that both sites were prone to mediate protein-protein interactions (PPI) (**Supplementary Fig. S1C**). To capture fibril periodicity, AmyBind first sampled the site space with a naïve helix positioned at the targeted interface and aligned to the fibril axis. Systematic grid and rotational sampling along Cartesian axes were then applied, and helices were designed in Rosetta^57^ under symmetry constraints, optimizing positions where fibril helical rise (∼4.8 Å) and helical turn rise (5.4 Å) were compatible (**Fig. 1A**).

Initial candidates were filtered by Rosetta metrics and further screened and docked with MaSIF-search^58^ for surface fingerprint complementarity, retaining only those with C_α_ root-mean-square deviation (RMSD) ≤ 2 Å to Rosetta models (**Fig. 1A**). This process yielded 298 and 182 seeds for Pol 1a and Pol 2b, respectively (**Supplementary Fig. S2**). These seeds were grafted onto a mini-protein library using Rosetta MotifGraft^70^ and further optimized through multiple Rosetta-based design strategies (**Supplementary Table S1**). A final set of ∼6,000 designs was selected based on Rosetta metrics **(Supplementary Fig. S3A**) and chosen for experimental testing as outlined in (**Fig. 1B**).

For Pol 1a, selected seeds were grafted onto helical hairpin scaffolds and optimized through full redesign or interface-focused refinement, with additional stabilization via disulfide bond engineering in Rosetta. In Pol 2b, the larger site accommodated multiple scaffold folds. As with Pol 1a, grafted designs underwent full redesign or two rounds of interface-focused refinement. In the first round of interface refinement (rd1), no constraints were applied to interface residue composition. In the second round (rd2), constraints were introduced to avoid methionines and glycines at the interface while limiting phenylalanine, tryptophan, and tyrosine to a maximum of 10% for each to reduce hydrophobicity and potential aggregation.

While some rd2 designs showed higher changes in binding energy (ddG) and lower shape complementarity scores than rd1 designs, they exhibited improved interface hydrogen bonding and fewer buried unsatisfied polar atoms (**Supplementary Fig. S3A**). This may have resulted from an increased presence of polar and charged amino acids, particularly serine, in rd2 compared to rd1 designs when analyzing interface compositions (**Supplementary Fig. S3B**).

### Screening and identification of mini-protein binders targeting aSyn fibrils

Approximately 6,000 *de novo* designed mini-protein binders were screened using yeast surface display against two aSyn fibril preparations (Ri and Fm)^62,63^, followed by two rounds of sorting. Yeast cells exhibiting binding signals were isolated for deep sequencing, and candidates showing ≥10-fold enrichment in the binding relative to the non-binding population were selected for further characterization (**Supplementary Table S2**). The binding of the 25 selected designs, adopting diverse folds (**Supplementary Fig. S4**), was then evaluated by yeast display against three aSyn fibril preparations (Ri, Fm, and F91)^62,63^ revealing distinct binding profiles among the designs (**Fig 2A**). Binding site specificity was further confirmed for a subset of these designs using interface knock-out mutants and scaffold-only controls, both of which abolished binding (**Supplementary Table S2)**.

**Figure 2.**
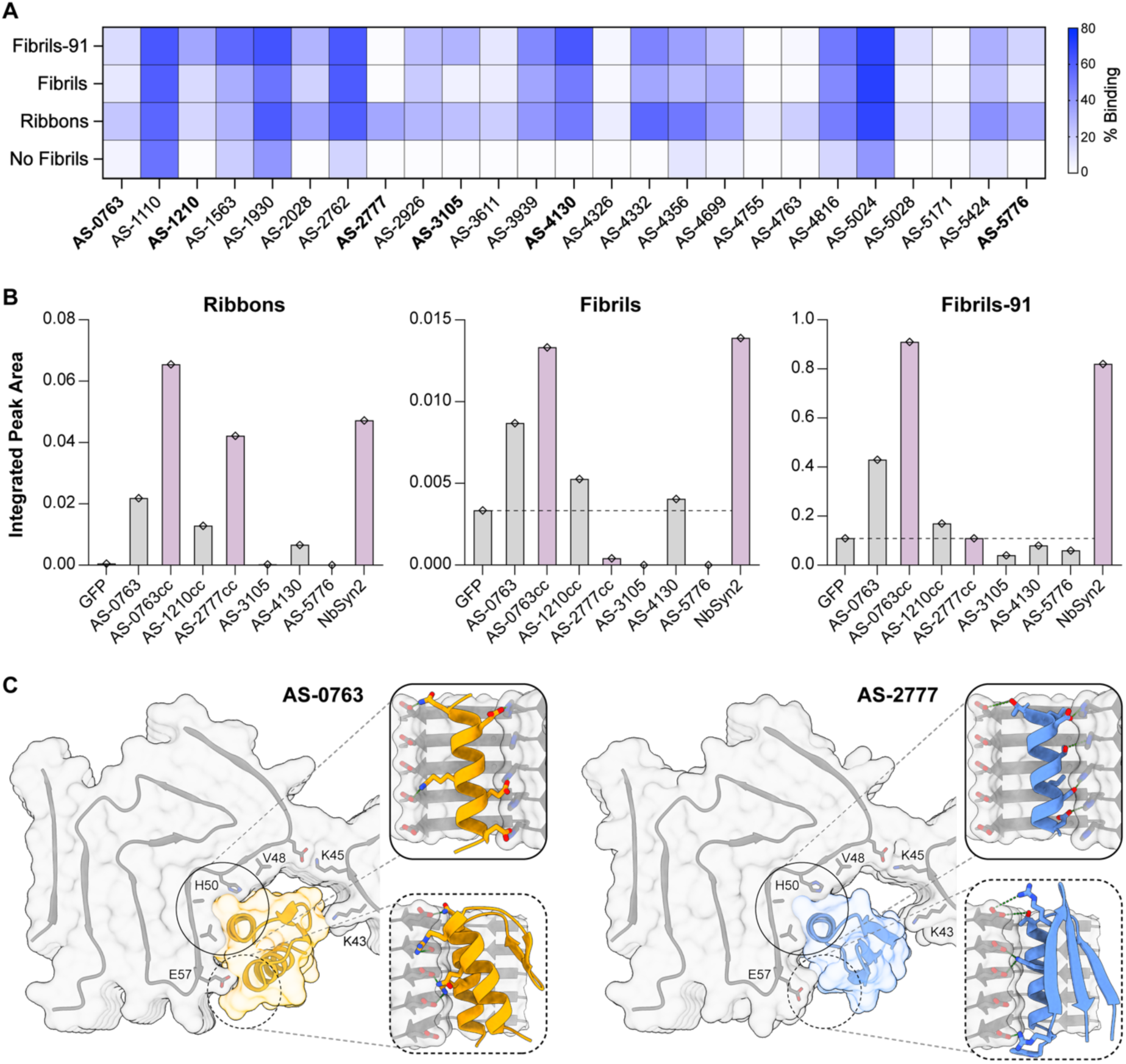
Screening and characterization of designed mini-protein binders. **(A)** Yeast surface display screening results for 25 identified candidates against three WT aSyn fibril preparations (Ri, Fm, and F91; 250 nM), with coloring indicating the percentage of yeast cells in the binding population. Experimentally validated designs are highlighted in bold. **(B)** GFP-fused designs were screened against the same WT aSyn fibril preparations using FIDA, with comparisons to controls: NbSyn2 (positive control, violet) and GFP (negative control, dashed line). Selected designs include AS-0763cc and AS-2777cc (violet). The integrated peak area (y-axis), represent signal from non-diffusive particles in FIDA, corresponding to GFP-fused proteins bound to fibrils, was calculated as described in [^75^] for Ribbons and Fibrils or by applying Simpson’s Rule Cumulative Integration^77^ for Fibrils-91, with individual measurements shown at 5 µM fibril concentration. **(C)** Rosetta structural models of lead binders AS-0763 (yellow) and AS-2777 (blue). Insets highlight polar interactions between the designed binding motif and the key fibril interface residue H50 (solid-line boxes/circles) and secondary interactions between the optimized scaffold and fibril residue E57 (dashed-line boxes/circles).

As a positive control, we employed NbSyn2, a nanobody known to bind the C-terminus of aSyn (epitope: Y136-A140)^71,72^, was expressed on the surface of yeast cells and tested during the screening. This epitope resides within the disordered C-terminal region of all available cryogenic electron microscopy (cryo-EM) structures of full-length aSyn fibrils and is therefore likely solvent-exposed and accessible^19,24^. Furthermore, its conservation across all aSyn species used in this study supports the use of NbSyn2 as a robust and reliable experimental control.

Selected designs were expressed as GFP-fusion proteins. Expression efficiency varied across constructs, with some exhibiting low or undetectable levels. To improve stability and folding, disulfide-stabilized variants were generated using the Disulfide by Design tool^73^, and evaluated *in silico* for foldability using AlphaFold2^74^, based on predicted local distance difference test (pLDDT) scores and C_α_ RMSD relative to the Rosetta models (**Supplementary Fig. S5**). The engineered disulfide bond was positioned away from interface residues, except in AS-2777cc, where one interfacial residue was substituted to accommodate the stabilizing disulfide bond, and enhance structural stability. Yeast display screening confirmed that the introduced disulfide bond did not affect binding, as AS-2777cc retained the same binding profile as AS-2777 (**Supplementary Fig. S6A**).

Four designs (AS-0763, AS-3105, AS-4130, and AS-5776) and three disulfide-stabilized variants (AS-0763cc, AS-1210cc, and AS-2777cc) were successfully expressed, and purified. Their binding to WT aSyn fibril preparations (Ri, Fm, and F91) was evaluated using FIDA. This technique distinguishes diffusive free binder from non-diffusive fibril–binder complexes undergoing transient incomplete separation in laminar flow^75^, enabling semi-quantitative assessment of binding. Because each aSyn fibril polymorph exhibits a distinct FIDA profile, comparisons between binders and controls were evaluated separately for each fibril type.

We then performed an all-against-all screen at a single fibril concentration (5 µM fibrils, 100 nM binder or control) to compare the binding of the various designs against NbSyn2 (positive control) and GFP (negative control). GFP has been shown not to bind fibrils possessing a “fuzzy coat”, such as those formed from full-length aSyn and tau^76^; therefore, detected binding signals primarily reflect interactions mediated by the designed proteins. Of the candidates tested, AS-0763cc and AS-2777cc produced binding signals beyond GFP background and comparable in magnitude to NbSyn2 (**Fig. 2B**). The polymorph specificity of AS-2777cc for aSyn Ri, previously observed in yeast surface display, was recapitulated in FIDA, whereas AS-0763cc displayed broad binding across all three aSyn polymorphs, consistent with the behavior of its parent design AS-0763 in both yeast and FIDA binding assays (**Fig. 2A, B, Supplementary Fig. S6A**). Notably, AS-0763cc exhibited higher binding signals than AS-0763 across all aSyn polymorphs, likely due to enhanced structural stability. Furthermore, yeast surface display confirmed the binding specificity of the parent designs, AS-0763 and AS-2777, as interface knock-out mutants and scaffold-only controls abrogated binding (**Supplementary Fig. 6B, C**). Based on these findings, AS-0763cc and AS-2777cc were selected for further characterization.

Examination of the Rosetta structural models of AS-0763 and AS-2777 revealed similar positioning of their helices within the fibril binding pocket and fibril-interacting residues on one protofilament (**Fig. 2C**). Both designs engaged fibril residues H50 and E57 through polar interactions, including salt bridges and hydrogen bonds. However, their distinct folds resulted in differences in how each binder interact with residue V48 from the same protofilament and residue K43 from the second protofilament constituting this fibrillar polymorph, as well as in the extent to which they fill the binding pocket. Despite these structural differences, their Rosetta metrics remained comparable (**Supplementary Table S3**). Notably, each design exhibited a distinct binding profile when tested against different aSyn fibril polymorphs (**Fig. 2A, B, Supplementary Fig. 6A**), underscoring both the challenges of computationally designing polymorph-specific binders and the importance of screening across multiple fibril polymorphs.

### Selective targeting of aSyn fibril polymorphs by designed mini-proteins

We performed a full titration with AS-2777cc and AS-0763cc against aSyn fibril preparations (Ri, Fm, and F91)^62,63^ using FIDA (**Fig. 3A, Supplementary Fig. S7A-C**). The titrations confirmed that AS-2777cc selectively binds aSyn Ri, whereas AS-0763cc binds to all fibril preparations. Binding specificity was further confirmed using scaffold-only controls, which showed no measurable signal in comparison with GFP (**Supplementary Fig. S7D-F**). The signal from the bound species in FIDA showed saturation only for titration of AS-2777cc with aSyn Ri. For this series, we assessed the binding affinity by fitting the FIDA-derived data using nonlinear four-parameter curve analysis (**Supplementary Fig. S8**). The binding affinity (*K_D_*) values from the two independent runs were consistent, indicating low-micromolar affinity of AS-2777cc for aSyn Ri (apparent *K_D_* = 2.7 ± 0.3 µM).

**Figure 3.**
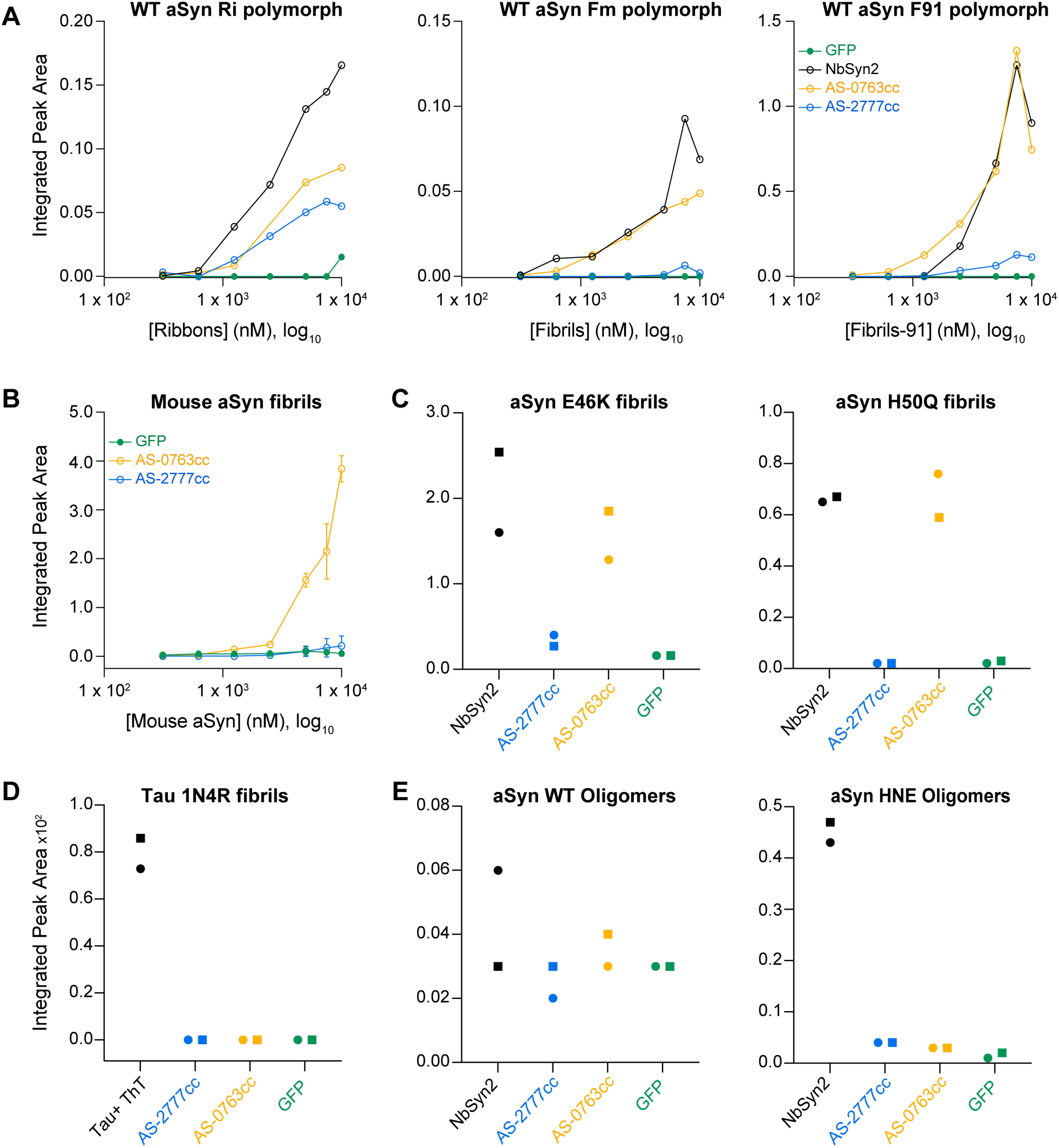
Assessment of designed binders’ specificity for aSyn fibril polymorphs using FIDA. **(A)** Titration binding curves of GFP-fused binders against aSyn fibril preparations, measuring binding to Ribbons (Ri), Fibrils (Fm), and Fibrils-91 (F91). Points represent areas of the non-diffusive peaks, obtained as described in [^75^], corresponding to the fibril-bound AS-0763cc (yellow), AS-2777cc (blue), NbSyn2 (positive control; black), or GFP (negative control; green). **(B–E)** FIDA screening of GFP-fused designs against **(B)** aSyn fibrils from mouse aSyn (5 µM), **(C)** human aSyn with familial mutations E46K and H50Q (5 µM each), **(D)** WT human Tau fibrils (1N4R isoform; 4 µM), and **(E)** WT aSyn oligomers (0.5 mg/ml) and HNE-modified aSyn oligomers (1 µM). Mouse aSyn fibrils were assessed by titration binding (n=2, mean±SD), while all others were screened at a single concentration with each point representing a technical replicate (markers: Run I, circles; Run II, squares). Integrated peak areas for (B, C, E) were calculated using Simpson’s Rule Cumulative Integration^77^, while for panel (D) areas were calculated as described in [^75^]. Integrated peak areas (y-axis) are compared to NbSyn2 or Tau+ThT (positive control) and GFP (negative control).

To assess sequence and conformational specificity of the binders, we first tested them against fibrils formed *in vitro* from mouse aSyn, whose sequence differs from that of human aSyn at seven residues, including the A53T substitution, located at the targeted interface. Cryo-EM structures of this preparation revealed an overall fold and inter-protofilament interface similar to the targeted polymorph^78^, making mouse aSyn fibrils an ideal control for evaluating conformational specificity and species differentiation. Across a full titration series measured by FIDA, AS-2777cc showed no binding to mouse aSyn fibrils, whereas AS-0763cc exhibited clear binding, confirming the specificity of AS-2777cc for the WT aSyn Ri polymorph (**Fig. 3B**).

Next, we tested the binders against fibrils formed *in vitro* from human aSyn carrying diverse mutations, including two familial mutations (E46K, H50Q) and seven variants containing one to three K-to-Q substitutions (**Supplementary Fig. S9A**)^79^, using FIDA at a single fibril concentration. The familial mutations either disrupt the formation of the targeted polymorph, as in E46K which eliminates the inter-protofilament K45-E46 stabilizing salt bridge within WT aSyn Pol 2b^80^, or alter a key interface residue essential for the predicted binding mode, as in the H50Q substitution. Fibrils formed by either familial mutant have been reported to adopt architectures that differ from the targeted polymorph^80–82^. The K-to-Q variant panel, which yields fibrils with distinct biophysical properties^79^, was used alongside the familial mutants to further probe AS-2777cc conformational specificity and to help delineate residues that contribute to the AS-0763cc binding site.

Across all nine mutant fibrils, AS-2777cc exhibited exceptional conformational specificity, showing binding comparable to GFP (negative control) except for a weak signal toward variant KQ3 (K32Q + K34Q), which was approximately four-fold lower than that of AS-0763cc. This weak interaction may reflect a subpopulation within the KQ3 preparation that adopts a fold resembling the WT Ribbons polymorph, although additional characterization of KQ3 fibrils is required to confirm this possibility. In contrast, AS-0763cc bound all but one variant (KQ5 = K58Q + K60Q) (**Fig. 3C, Supplementary Fig. S9B)**. This broad binding profile across three distinct WT polymorphs, eight human aSyn mutant fibrils, and mouse aSyn fibrils indicates that AS-0763cc engages a conserved site among these fibrils. Its lack of binding to the KQ5 variant further suggests that residues K58 and K60 or their spatial organization within the fibrillar scaffold play a key role in mediating AS-0763cc interactions with the aSyn fibrillar polymorphs.

To further evaluate target specificity, we assessed the binding propensity of AS-2777cc and AS-0763cc to two fibrillar isoforms of WT Tau (1N3R and 1N4R)^83^, which share no sequence identity with aSyn, using FIDA. Tau fibril non-diffusive behavior in FIDA was confirmed separately by mixing them with ThT for detection (**Supplementary Fig. S10A**). No measurable binding above GFP (negative control), was observed for either binder, indicating their selectivity for aSyn fibrils (**Fig. 3D, Supplementary Fig. S10B**).

Next, we examined whether AS-2777cc and AS-0763cc bind aSyn oligomers using FIDA. Two forms of human WT aSyn oligomers^84^ were tested: oligomers generated directly from monomers (WT; 0.5 mg/mL) and oligomers produced via cross-linking with 4-hydroxy-2-nonenal (HNE; 1 µM). Oligomer preparations were first analyzed in the absence of binders to assess their non-diffusive behavior in the assay (**Supplementary Fig. S11A**). Neither design showed detectable binding, confirming their selectivity for aSyn fibrils (**Fig. 3E, Supplementary Fig. S11B**). Interestingly, NbSyn2 bound HNE-modified, but not WT, aSyn oligomers, potentially reflecting differential accessibility of the aSyn C-terminus in these species.

Lastly, we used FIDA to assess binder interaction with aSyn monomer, but capillary adhesion of GFP-labeled species hindered quantification of changes in diffusive species hydrodynamic radius. Instead, isothermal titration calorimetry (ITC) confirmed that neither AS-2777cc nor AS-0763cc bound monomeric aSyn, in contrast to NbSyn2 as shown before^85^ (**Supplementary Fig. S12**).

Altogether, these results establish that AmyBind can generate binders with exclusive specificity for aSyn fibrils, without interacting with monomeric or oligomeric aSyn species nor fibrils formed by other amyloidogenic proteins. AS-2777cc demonstrated selective recognition of the WT aSyn Ri polymorph, whereas AS-0763cc appeared to bind a conserved motif shared across twelve of the thirteen screened aSyn fibrils formed from WT, mutant human aSyn and mouse aSyn (**Fig. 3A-C, Supplementary Fig. S9B**). These findings highlight the potential of AmyBind to yield fibril-selective binders, while also underscoring the necessity of thorough experimental screening to identify candidates with true polymorph specificity. This specificity landscape provides an essential foundation for the subsequent characterization and functionalization of the identified binders.

### Binders do not interfere with aSyn fibril elongation *in vitro*

Following the validation of AS-2777cc and AS-0763cc binding specificity, we evaluated their effect on aSyn aggregation using a ThT assay. Experiments were conducted in the presence of fragmented, preformed Ri or Fm seeds (2.5% of aSyn monomer) under quiescent conditions, ensuring that fibril elongation was the dominant process.

Seeds were incubated with binders or scaffold-only controls at a 1:2 ratio, corresponding to a 10-fold molar excess of binders relative to the estimated number of fibril binding sites, assuming one site spans five monomer planes. ThT binding kinetics showed no measurable differences in aggregation profiles for either fibril preparation compared with experimental controls, including elongation kinetics measured in the absence of binders or in the presence of scaffold-only constructs, GFP, or NbSyn2, which has previously been reported not to affect aSyn aggregation^71^ (**Fig. 4A, Supplementary Fig. S13A**). SDS-PAGE analysis of residual aSyn monomer at the assay endpoint further confirmed the absence of binder-induced inhibition of fibril elongation (**Fig. 4B, Supplementary Fig. S13B, S14A, and S15A**).

**Figure 4.**
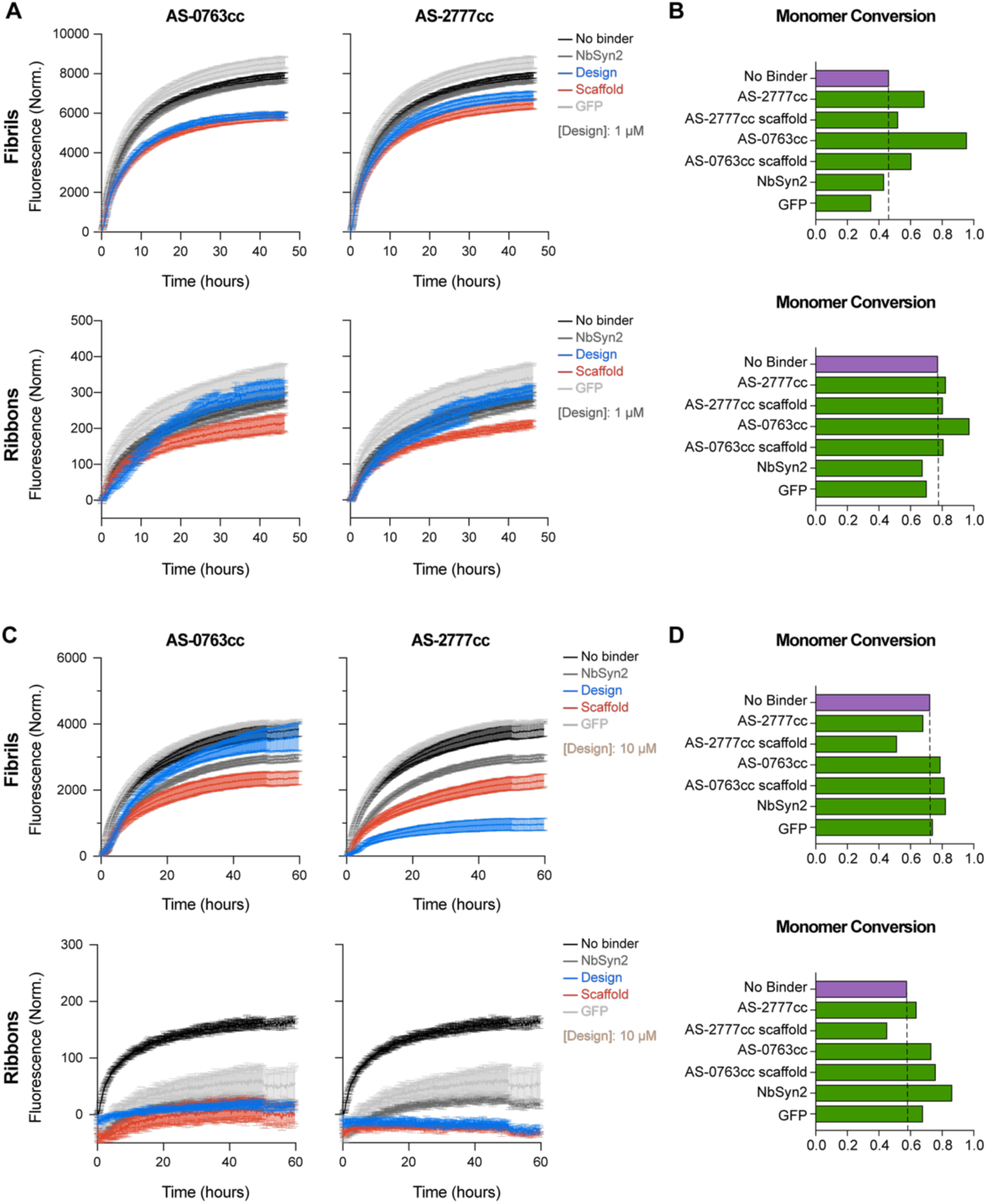
Effect of designed proteins on the elongation kinetics of aSyn seeds monitored by ThT binding and SDS-PAGE. **(A)** aSyn aggregation kinetics followed using ThT showing the effect of 1 µM designs on the elongation of 0.5 µM aSyn seeds at a 2:1 binder:seeds stoichiometry (in monomer equivalents) in the presence of 20 µM aSyn monomer (**Fibrils, top; Ribbons, bottom**) **(B)** Quantification of aSyn monomer conversion into fibrils at the endpoint of the elongation experiments in (A) using SDS-PAGE. The dashed line represents aSyn monomer conversion in the absence of binders. **(C)** ThT aggregation kinetics curves showing the effect of 10 µM designs on the elongation of 0.5 µM aSyn seeds at a 1:2 binder:aSyn monomer stoichiometry in the presence of 20 µM aSyn monomer (**Fibrils, top; Ribbons, bottom**) **(D)** Quantification of aSyn monomer conversion into fibrils at the endpoint of elongation experiments in (C) after fibrils sedimentation by centrifugation using SDS-PAGE. The dashed line represents aSyn monomer conversion in the absence of binders. SDS-PAGE gels used for monomer quantification are shown in **Supplementary Fig. S14**. ThT kinetics experiments were performed in triplicate (outlier measurements were excluded, AS-0763cc experiments with Fibrils shown in C were performed in duplicate); mean values are plotted, and error bars represent standard deviation (SD).

Increasing the binder concentration to 10 µM relative to aSyn monomers (20 µM) led to a reduction in ThT fluorescence in most cases (**Fig. 4C, Supplementary Fig. S13C**). However, SDS-PAGE quantification of residual aSyn monomer indicated no inhibition of aSyn aggregation, as monomer depletion remained comparable to that observed in the no-binder control (**Fig. 4D, Supplementary Fig. S13D, S14B, and S15B**). This suggests that the binders compete with ThT binding to fibrils.

To examine concentration-dependent effects, binders were tested at two concentration ranges: low (0.125–1 µM, Fibrils: **Supplementary Fig. S16A**, Ribbons: **Supplementary Fig. S17A**), and high (1.25–10 µM, Fibrils: **Supplementary Fig. S16B**, Ribbons: **Supplementary Fig. S17B**). ThT fluorescence progressively decreased with increasing binder concentration only in the higher range. However, since 10 µM binders did not affect aSyn monomer aggregation (**Fig. 4D, Supplementary Fig. S13D**), the observed signal reduction was likely due to competition between binders and ThT binding rather than inhibition of fibril elongation. This was confirmed by repeating elongation reactions in tubes without ThT under the same conditions (0.5 µM aSyn Ri or Fm seeds, 20 µM aSyn monomer, and 10 µM binder), run in parallel with **Fig. 4C** and **Supplementary Fig. S13C** experiments. Quantification of residual aSyn monomer at the experiment’s endpoint after centrifugation using SDS-PAGE confirmed the absence of inhibitory effect on aSyn aggregation, except for AS-2777cc scaffold (**Supplementary Fig. S18**).

The lack of inhibition of seeded elongation strongly indicates that the binders interact with fibril surfaces rather than their ends. This is in accordance with our design strategy and has relevant implications for future applications, including fibril detection and targeted fibril degradation^86^, providing a foundation for the development of fibril-based diagnostics and therapeutics.

### Binders exhibit polymorph-specific colocalization without influencing fibril uptake or seeding in cellular models

To assess whether the binders are co-internalized with aSyn fibril polymorphs or influence their uptake in cellular disease-like models, we exposed Neuro2A and HeLa cells to ATTO647-labeled aSyn polymorphs (Fm or Ri) together with GFP-fused binders or controls (1µM each). Live-cell imaging was performed immediately after addition of trypan blue to quench external GFP fluorescence from the non-internalized binders, thereby enabling selective detection of internalized GFP-fused binders (FITC channel) and ATTO647-labeled polymorphs (Cy5 channel) (**Fig. 5A**).

**Figure 5.**
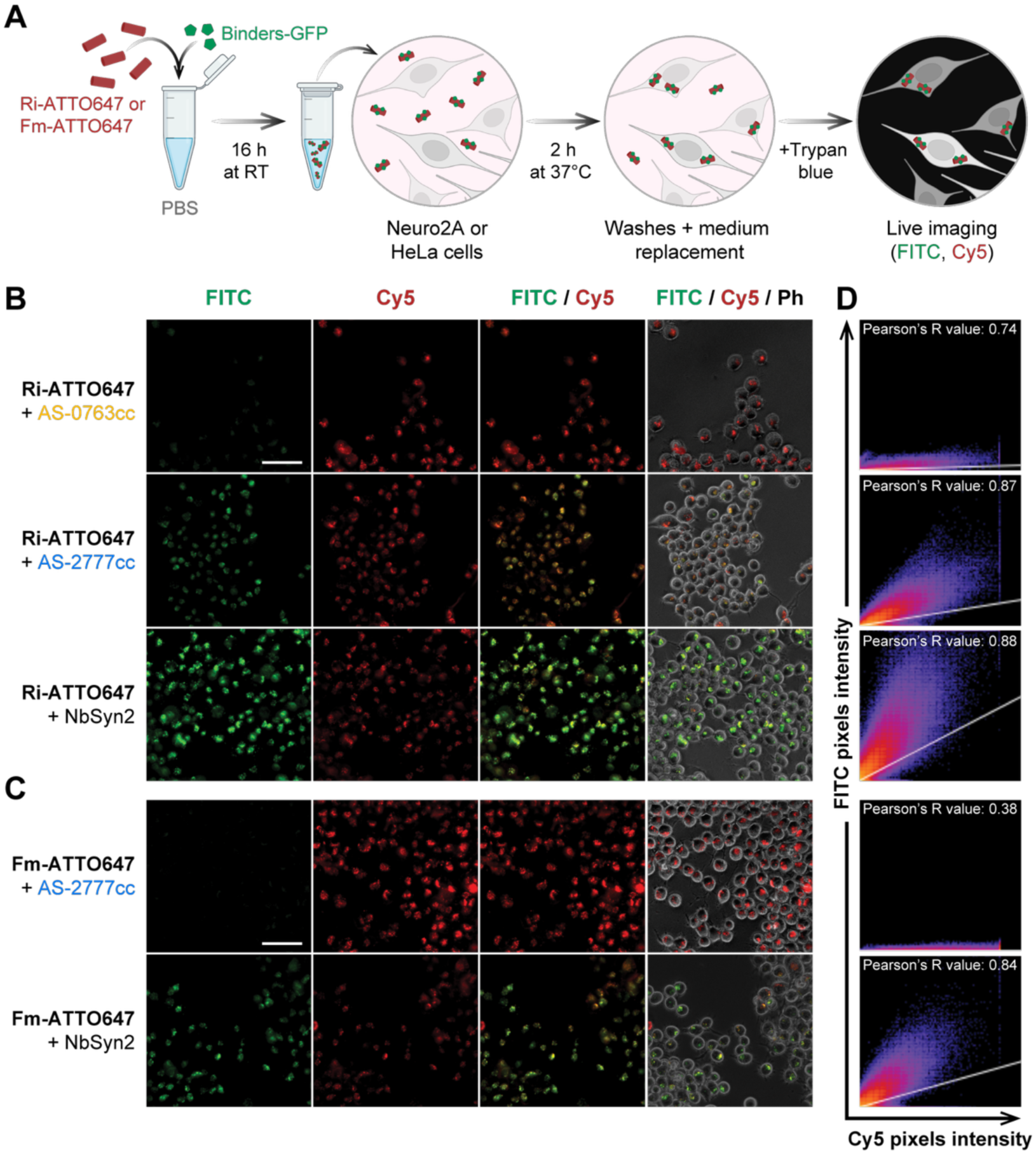
Internalization of aSyn polymorphs labeled with binders in Neuro2A cells. **(A)** Experimental outline for assessing the effect of GFP-fused binders on the internalization of fragmented ATTO647-labeled aSyn polymorphs, Ribbons (Ri-ATTO647) or Fibrils (Fm-ATTO647) (average length: 45 nm). **(B, C)** Representative live-cell fluorescence micrographs acquired after Neuro2A cells were exposed to Ri-ATTO647 **(B)** or Fm-ATTO647 **(C)** together with GFP-fused binders (FITC: green; Cy5: red; Ph: Phase contrast). Cy5 signals correspond to aSyn polymorphs bound to or internalized by cells, whereas FITC signals reflect exclusively internalized GFP-fused binders. Additional control conditions are shown in **Supplementary Fig. S20. (D)** Scatter plots showing pixel-based colocalization between the Cy5 (x-axis) and FITC (y-axis) channels for the micrographs shown in **(B)** and **(C)**. Object-based colocalization analysis is provided in **Supplementary Fig. S21A.** The experiment was performed in two cell lines. Data obtained in HeLa cells are shown in **Supplementary Fig. S19–S21.** Scale bar = 50 µm.

In the presence of the Ri polymorph, AS-0763cc, AS-2777cc, and the positive-control nanobody NbSyn2 were internalized and colocalized with Ri in both Neuro2A (**Fig. 5B)** and HeLa cells (**Supplementary Fig. S19A**). In contrast, in the presence of the Fm polymorph, NbSyn2 was internalized and colocalized with Fm, whereas AS-2777cc was not, further confirming its specificity for the Ri polymorph (Neuro2A: **Fig. 5C**; HeLa: **Supplementary Fig. S19B**). Colocalization of GFP-fused-binders with ATTO647-labeled aSyn polymorphs was further assessed using pixel-based (**Fig. 5D**, **Supplementary Fig. S19C**, **S20A**) and two-dimensional object-based analyses (**Supplementary Fig. S21**). In both cell lines, AS-2777cc exhibited stronger colocalization with Ri than AS-0763cc, whereas NbSyn2 showed robust colocalization with both polymorphs (Neuro2A, **Fig. 5B, C, Supplementary Fig. S21A**; HeLa, **Supplementary Fig S19**, **S21B**).

No internalization was detected for the negative controls, including AS-0763cc scaffold and GFP alone, in the presence of the Ri polymorph, supporting the conclusion that binding arises from the designed interfaces rather than the GFP tag (Neuro2A: **Supplementary Fig. S20A**; HeLa: **Supplementary Fig. S19A**). In the absence of fibrils, neither binders nor controls were internalized by either cell line, confirming that binding to fibrils is required for internalization within the time period studied (**Supplementary Fig. S20B, C**). By contrast, ATTO647-labeled aSyn polymorphs were detected in both cell lines, regardless of the presence or absence of binders and controls, indicating that the binders did not interfere with fibrils uptake (**Fig. 5B, C**, **Supplementary Fig. S19, S20A**).

To determine whether binder engagement influences downstream fibril-induced aSyn seeding, we used HeLa cells expressing human WT aSyn fused to mCherry. Cells were exposed to unlabeled aSyn polymorphs (Fm or Ri) with either GFP-fused binders, their respective negative control, GFP alone (1 µM each) or no binders (PBS). Following exposure to fibrils, accumulation of aSyn-mCherry into bright punctae, indicative of relocalization and seeding^62^ was assessed by confocal microscopy (**Supplementary Fig. S22A, B, S23A, B**). In the absence of fibrils, cells displayed a homogeneous cytoplasmic mCherry signal (**Supplementary Fig. S23C**). Quantification of punctae per cell in the presence of fibrils revealed no differences across experimental conditions (i.e., with or w/o binders), indicating that the binders did not alter fibril-induced seeding of expressed Syn-mCherry monomers (**Supplementary Fig. S22C, S23D**). This finding is consistent with their designed lateral binding mode and with the lack of inhibition observed in the *in vitro* elongation assays (**Fig. 4**).

In line with the internalization data, AS-2777cc and NbSyn2 partially colocalized with aSyn-mCherry punctae following exposure to the Ri polymorph. In contrast, after exposure to the Fm polymorph, only NbSyn2 exhibited partial colocalization, whereas AS-2777cc localized predominantly at the cell membrane similar to the negative control GFP. This membrane-associated signal may reflect adherence of extracellular binders trapped in the culture medium (**Supplementary Fig. S22, S23**).

Together, these results confirm the polymorph specificity of the Ri-selective binder AS-2777cc, consistent with its Ri-specific internalization and colocalization. Although both binders (AS-2777cc and AS-0763cc) and the positive-control nanobody NbSyn2 were internalized and colocalized with fibrils, they neither altered fibril uptake nor seeding in our cellular models and remained stable and detectable within cells following exposure to fibril–binder mixtures.

## Discussion

Current treatments of neurodegenerative diseases such as PD and related synucleiniopathies provide only symptomatic relief^87^ and diagnosis is typically based on clinical symptoms^88^. However, efforts to identify early-stage biomarkers are gaining momentum, as early intervention may be critical for effective disease management^89^. In PD, aSyn fibrils represent a promising biomarker candidate due to their key roles in pathology formation and propagation^12^ and disease progression^11,90^. Furthermore, the detection of aSyn fibrils in biological fluids of patients^91^, using aSyn seed amplification assays, supports the diagnosis of PD with greater than 90% accuracy^92–95^. Recent advances in cryo-EM have enabled structural characterization of aSyn fibrils formed both *in vitro* and in disease contexts, revealing a diverse structural landscape^19,24^. The observation that distinct fibril polymorphs are associated with different synucleinopathies^19^, suggests a possible structure-based classification of synucleinopathies and underscores the potential for developing polymorph-and disease-specific binders and diagnostics. However, the structural divergence between *in vitro* and pathological fibril folds poses a challenge for the development of polymorph-selective diagnostic or therapeutic molecules^29,96^.

Current efforts leveraging *in vitro*-assembled aSyn fibrils in both screening-based drug discovery and structure-based protein design have led to the development of small-molecule and protein-based binders that achieve target selectivity but rarely exhibit polymorph specificity^17,30,31,55,97–99^. Structure-informed design of peptide binders exploiting differential solvent accessibility of a protein segment across fibril conformers has enabled fibril polymorph differentiation; however, the extent to which such binders distinguish a polymorph from another among the diverse conformational states of aSyn remains an open question^56^. Collectively, these studies face key challenges including, the limited ability to reproduce pathological fibril folds *in vitro* and a reliance on targeting linear, continuous, solvent-accessible protein segments in the fibrillar state, which inherently complicates the discrimination of binders between distinct oligomeric states of amyloidogenic proteins.

To address these limitations and explore the potential for developing polymorph-specific binders against aSyn fibrils, we developed AmyBind, a computational framework for designing *de novo* amyloid-selective mini-protein binders with conformational specificity for distinct fibrillar polymorphs. This pipeline integrates Rosetta^57^ for binder design and optimization with MaSIF^58^ for interface scoring and filtering. To account for the unique structural features of amyloid fibrils, including their helical symmetry and rise, AmyBind employs a discrete grid-sampling algorithm to optimize binder positioning. Unlike existing binders that target solvent-exposed β-sheet regions perpendicularly to the fibril axis, for fibril end capping^17,55^ or selective surface binding^56^, AmyBind introduces the first computationally designed binders that engage polymorph-specific sites in a parallel orientation along the fibril axis. This configuration enables conformational specificity by requiring a composite interface formed by multiple stacked monomeric units, thereby allowing both polymorph differentiation and species selectivity.

As a proof of concept, we targeted sites from two distinct *in vitro* polymorphs of recombinant human WT aSyn fibrils, Polymorph 1a^59^ and Polymorph 2b^60^. Using yeast surface display coupled with fluorescence-activated cell sorting and next generation sequencing, we identified lead candidates from an initial pool of ∼6,000 designs. Subsequent biochemical characterization revealed two aSyn fibril-specific binders, AS-0763cc and AS-2777cc, with no detectable binding to fibrils formed by other amyloidogenic proteins, such as Tau, or to other aSyn species, including monomers and oligomers. Although both binders exhibited target selectivity, our experimental characterization showed that AS-2777cc engaged the intended target interface and displayed polymorph specificity, binding exclusively to the aSyn Ri polymorph with minimal interaction with other WT or mutant aSyn fibrils tested. In contrast, AS-0763cc appeared to recognize a distinct but conserved motif shared across nearly all tested aSyn fibril types, with the exception of the KQ5 mutant variant. Consistent with their designed engagement of lateral fibril surfaces, neither binder affected fibril elongation *in vitro*, a process driven by monomer addition at the fibril ends^100^. In cell-based models, both binders co-internalized with aSyn fibrils via endocytosis and retained polymorph-specific colocalization without altering fibril uptake or seeding.

Our findings highlight the potential of AmyBind to generate aSyn fibril-specific binders that discriminate among distinct conformational states of aSyn and support its broader application to other amyloid fibrils. Because these binders do not affect fibril elongation or seeding, yet retain polymorph-specific colocalization in cellular models, they could serve as non-perturbative probes for accurate quantification of fibril formation and studying fibril-mediated cellular, propagation and pathogenic properties in a target- and polymorph-selective manner.

Furthermore, the distinct polymorph selectivity profiles of the binders could be leveraged for different applications. Broadly reactive binders that capture the diversity of aSyn fibrils, such as AS-0763cc, could support the development of diagnostics or therapeutic strategies for disease groups defined by a shared fibrillar protein, such as synucleinopathies. By contrast, polymorph-selective binders, such as AS-2777cc, could enable the development of fibril-targeting disease-specific therapeutics^86,101–103^, for example. Through chimeric derivatives of binders developed as protein-based PET tracers for disease diagnosis^104,105^, or as modules for targeted fibril clearance through intracellular^86,101–103^ or extracellular^106–110^ degradation pathways. Targeting fibrils to inhibit their growth, seeding activity and propagation represents a viable disease-modifying strategy to slow the progression of PD and other NDs^11,55,86,102,111^.

For this proof-of-concept study, we used readily available *in vitro* WT aSyn fibrils, given the difficulty of reproducing pathological fibril folds *in vitro*^19,25^. Notably, protofilament folds resembling the *in vitro* Polymorph 2b^60^ structure used here have been observed in fibrils formed *in vitro* from mouse aSyn^78^ as well as from human aSyn seeded with MSA patient-derived material, termed Fold A and Fold B^25^. Structural alignment revealed close similarity between Polymorph 2b and the MSA-amplified Fold B fibril, including conservation of the AS-2777cc targeted site (**Supplementary Fig. S24A, B**). Consistent with this structural similarity, docking of AS-2777cc to MSA-amplified Fold B fibrils using MaSIF-search yielded interface surface complementarity scores that met our predefined thresholds (**Supplementary Fig. S24C**). These analyses suggest that AS-2777cc may engage disease-relevant fibrillar folds and support its potential translational relevance, although experimental validation with amplified fibrils will be required to establish this directly.

For some disease-associated amyloids, such as Tau fibrils in Alzheimer’s disease^112^, pathology-related folds can be reproduced *in vitro*^113^, facilitating their use in drug discovery and binder design. In contrast, for other amyloid fibrils for which reproducing pathological folds *in vitro* remains challenging^29^, future design campaigns using AmyBind could benefit from incorporating comparative structural analysis tools, such as FibrilSite^96^, to identify sites conserved between pathological and *in vitro*-assembled fibrils. This strategy could help select more representative *in vitro* models and enhance the translational relevance of designed binders.

In conclusion, we demonstrated the feasibility of computationally designing *de novo* polymorph-specific binders against amyloid fibrils using our developed method AmyBind. Although the current implementation yielded two aSyn fibril-specific binders, integration of recent advances in protein design and structure prediction, including RFdiffusion^47,48^, ProteinMPNN^114^, Boltz^115^, or AlphaFold^74,116^ could improve binder affinity, stability, and experimental success rates^46,117–119^. We believe that the combination of these approaches and the expansion of the amyloid structure landscape offer unique opportunities to turn fibrils into druggable targets and transform diagnostics and drug discovery in NDs.

This study has two main limitations: first, we employed FIDA to evaluate fibril–binder interactions; however, quantitative determination of binding stoichiometry and true affinity by FIDA was constrained by technical factors associated with such in-solution measurements, including partial binder adhesion to the capillary and signal variability across capillaries. Second, structural characterization of AS-2777cc in complex with the Ribbons preparation by cryo-EM was impeded by the lack of fibril helicity and by heterogeneity in fibril length following fragmentation. Nevertheless, we obtained compelling biochemical evidence for specific engagement of the targeted fibril interface, supported by appropriate experimental controls for both the binder and the fibril. Although Polymorph 2b was identified within the aSyn “Fibrils” polymorph preparation^63^, it constituted only approximately 20% of the sample population, with the majority corresponding to Polymorph 2a, in which the targeted site is inaccessible^60^. This population imbalance likely accounts for the absence of detectable binding to this preparation. In contrast, within the Ribbons polymorph the targeted region resides in a comparatively less structurally constrained context and remains solvent accessible along the fibril surface, thereby enabling productive engagement by AS-2777cc^120,121^.

Conformation-specific binders enable differential disease diagnosis by distinguishing fibril polymorphs, a capability that has rarely been achieved by other targeting approaches, including small molecules^43^ and antibodies^84^. Their fibrillar form selectivity can also be leveraged to promote fibril clearance, thereby potentially limiting disease progression and propagation^86,101–103^. This approach, combined with advances in peptide delivery systems for blood–brain barrier penetration^122,123^, paves the way for a new class of diagnostics and therapeutics for currently incurable diseases.

## Data availability

All data are accessible on Zenodo at https://doi.org/10.5281/zenodo.18007441

## Code availability

The code and scripts for running AmyBind are available online at https://github.com/LPDI-EPFL/amybind

## Acknowledgment

We thank SCITAS at EPFL for support in running our pipeline and analysis. We thank Davide Demurtas and Stéphanie Clerc-Rosset for their assistance with fibril TEM imaging and characterization (BioEM Facility at EPFL). We are grateful to Pablo Gainza and Andreas Scheck for their help with MaSIF implementation in our computational pipeline, Martin Pacesa for assistance with Cryo-EM structural analysis, and Lukas Jarosch for testing AmyBind grid generation on the high-performance computing cluster (LPDI at EPFL). We thank Senthil T. Kumar, Pedro Magalhães, and Rajasekhar Kolla for their support with the recombinant WT aSyn protein production and purification used in this study (LMNN at EPFL). We also thank Tobias Mühlethaler and Timothy Sharpe for carrying out the ITC measurements and analyses (Biophysics Facility at Biozentrum, University of Basel). The ITC measurements were supported by the SNSF R’Equip Grant 213436 “Efficient Characterization of Biomolecular Interactions by Automated Isothermal Titration Calorimetry”. This research received financial support from EraPerMed DEEPEN-iRBD project and Agence Nationale de la Recherche (ANR-22-PERM-0006) to RM and SYNSELECT project (ANR-21-CE16-0039-01) to N.L.R. and benefited from access to the electron microscopy facility Imagerie-Gif. This research was supported by the European Research Council, through an ERC CoG grant to AKB (EMMA, 01088163) and a MSCA grant to AK (101106115). BioRender was used to create Fig 5A, Supplementary Fig. S22A.

## Contributions

A.S., Z.H., H.A.L., and B.E.C. conceived the work. A.S. and Z.H. developed AmyBind and performed the computational design and analysis. A.S. performed the high throughput screening for lead identification. A.S., S.G. and J.S. expressed, purified, and performed biophysical characterization of the binders. A.S., N.L.R., A.K., and R.M. generated aSyn fibrillar polymorphs and tau fibrils. A.S. and A.K. conducted the *in vitro* characterization of binder-fibril interactions. N.L.R. and R.M. performed experiments in cellular models. A.S. and N.L.R. prepared the illustrations. A.S., A.K., N.L.R., and Z.H. wrote the manuscript with input from all authors.

## Ethics declarations

### Competing interests

EPFL has filed a provisional patent application that incorporates the findings presented in this Article. A.S, Z.H., H.A.L., and B.E.C. are named as coinventors of the patent (European Patent Office, EP 25224619.4). H.A.L is the founder and CEO of ND BioSciences, a spinoff from the Lashuel lab focusing on developing novel therapies and diagnostics for neurodegenerative diseases. The remaining authors declare no competing interests.

## Methods

### Prediction of Protein interaction sites on Fibrils

The MaSIF-site tool from the Molecular Surface Interaction Fingerprinting (MaSIF) framework^58^, trained to predict protein surface regions with a propensity to form protein–protein interactions (PPIs), was applied to aSyn fibril polymorphs structures (1a, PDB: 6CU7; 2b, PDB: 6SST). Protein surfaces were preprocessed and decomposed into patches through upstream computational steps before being input to MaSIF-site. MaSIF-site assigned a per-vertex score to each surface point, indicating its likelihood of becoming a buried surface area within a PPI. To identify potential interface patches on target protein, the MaSIF-site output was further divided into 12 Å overlapping patches, with an overall score for each patch calculated by averaging the per-vertex prediction for all points within the patch.

### Docking and Filtering De-Novo Designs Using MaSIF-Search

The MaSIF-search tool from the MaSIF framework^45,58^, designed to evaluate surface complementarity between interacting proteins, was used to identify potential interacting *de novo* designs for fibril sites. Briefly, the *de novo* designs were provided as the binding seed database to MaSIF-search. MaSIF-site predictions and MaSIF-search fingerprints were computed for both the target fibril and the binders.

The docking process in MaSIF-search involved two stages. In the first stage, patches from the target site and the designs were compared, and selections were made based on fingerprint distances (Euclidean distance between target and seed fingerprint). In the second stage, the selected designs were aligned to the target using the random sample consensus (RANSAC) algorithm implemented in Open3D^124^. This alignment utilized fingerprints of all the vertices in the target and design patches to identify the best vertex correspondences.

The uniqueness of the design-fibril complex poses limited the use of the interface post-alignment (IPA) neural network for scoring the aligned patches. Hence, we relied on the patch-to-patch surface fingerprint similarity described by the descriptor distance score. This score is computed as the sum of the squared inverse Euclidean distances between the target fingerprint and the binder’s inverted fingerprint, including only those within a 1.5 Å threshold, meaning the higher the score the better the surface complementarity. To identify the thresholds for the descriptor distance score, MaSIF-search was benchmarked using 5,188 patch pairs from 872 native complexes in the published MaSIF-search test set. The MaSIF-search IPA scores for docked patches showed that 70% of the complex patches (3,665 / 5,188; 826 native complexes) exceeded the recommended threshold (0.8, dashed line) for a potential real interaction (**Supplementary Fig. S25A**). The median values for the calculated descriptor distance scores were used as references for the quality of the *de novo* designed interfaces (Full set: 17.7; Patch matches with IPA score ≥ 0.8: 21.8; **Supplementary Fig. S25B**)

Finally, the structural similarity between the MaSIF-search docked and the initial Rosetta-designed model was evaluated by (1) aligning the target structures and (2) calculating the C_α_ RMSD of the designs via Biopython package^125^.

### AmyBind: a Surface motif regularity-guided binder design pipeline

Amyloid fibril structures possess unique attributes, such as helical symmetry, helical twist, and helical rise, resulting from the stacking of identical amyloid-forming monomeric protein units, resulting in unique repeating surface patterns. To accommodate these patterns in the binder design process, we developed a framework comprising discrete space sampling, motif design in Rosetta^57^, filtering using MaSIF-search^45^, and motif grafting and design refinement using Rosetta^57^. This pipeline is designed to address the structural and functional intricacies of amyloid fibrils, enabling the development of binders with high target specificity.

As a proof of concept, we targeted two sites from distinct *in vitro* polymorphs of human WT aSyn fibrils (**Fig. 1A**): Polymorph 1a^59^ (Pol 1a, PDB: 6CU7) and Polymorph 2b^60^ (Pol 2b, PDB: 6SST). Because only the fibril core is structurally resolvable, whereas the N- and C-termini remain disordered and can form a “fuzzy coat” surrounding the core^59,60^, site select required careful inspection of the cryo-EM maps in addition to the deposited PDB structures (Pol 1a: EMD-7618; Pol 2b: EMD-10305). This strategy minimized the risk of selecting regions near the disordered termini that could sterically hinder binder engagement.

The Pol 1a site is located in the C-terminus of aSyn and is formed by residues **S87**, I88, A89, **A90**, A91, **T92**, G93, F94, **V95**, K96, **K97**. In contrast, the Pol 2b site is positioned in the N-terminus, spanning two interacting aSyn units and formed by residues **K43**, T44, **K45** from one unit and **E46**, G47, **V48**, V49, **H50**, G51, V52, **A53**, T54, **V55**, A56, **E57** from the adjacent unit. Residues with solvent-accessible side chains are highlighted in bold.

### Discrete site space sampling

A naïve helix was positioned parallel to the fibril growth direction within the desired fibril site. The site space was systematically sampled by incrementally moving the naïve helix in 0.75 Å steps along each of the site’s Cartesian coordinates, one at a time, using the fibril’s eigenvectors (principal directions) as references to guide the movement direction. After each step, the helix-fibril complex was exported to serve as a starting point for motif design in Rosetta^57^.

### Motif *de novo* design in Rosetta

The naïve helix in the input helix-fibril complex was designed in Rosetta^57^ using fixed backbone design, incorporating a penalty for buried unsatisfied polar atoms in the scoring function^126^ and a sequence symmetry constraint between the helix residues *i* and *i*+7 to account for the repetitive nature of the fibril. To identify the optimal helix placement at a given position, rotational space sampling was performed by iteratively repeating the design process 71 times, rotating the helix 5° around its axis in each iteration. The resulting helices were filtered first based on Rosetta metrics including binding energy (ddG, rigid body docking: ≤ -20) and shape complementarity (≥ 0.6) yielding 451 and 4,144 unique seeds for Polymorph 1a and Polymorph 2b, respectively. The filtered designs were further evaluated using MaSIF-search as described above resulting in 298 and 182 seeds for Polymorph 1a and Polymorph 2b, respectively. The final selected helices were used for grafting into mini-proteins using Rosetta^57^.

### Motif grafting and design refinement

The selected designed helices were matched to a database of 4,347 small globular proteins (<100 amino acids) sourced from the PDB^127^, two computationally designed mini-protein libraries^128,129^ and an AlphaFold2 proteome prediction database^130,131^ using Rosetta MotifGraft, as previously described in [^45^]. For Polymorph 1a, the site size restricted grafting to scaffolds with fewer than 38 residues, whereas for Polymorph 2b, the larger site size accommodated multiple folds, including EHEE, HEEH, and HHH (**Supplementary Fig. S4**). Grafts were filtered based on Rosetta metrics, including binding energy (ddG with side chain repacking; Pol 1a: ≤ -40; Pol 2b: ≤ -25), graftClash score (0), and graftRMSD (≤ 2), yielding 5,597 and 4,963 grafts for Polymorph 1a and Polymorph 2b, respectively, to be further optimized using multiple design protocols in Rosetta (**Supplementary table S1**).

For Polymorph 1a, seeds were grafted onto hairpins and optimized using four strategies (FO, SO, SO_pm_, and SO_alt_; **Supplementary Table S1**). Except for SO_alt_, hairpin backbone minimization was permitted during design. These designs were filtered based on Rosetta metrics including binding energy (ddG with side chain repack ≤ -50) and shape complementarity (≥ 0.65). The filtered designs were pooled and stabilized via disulfide bond engineering in Rosetta (Disulf; **Supplementary Table S1**). The selected disulfide-stabilized variants then underwent a final round of refinement to diversify the designed interfaces (Disulf_opt_; **Supplementary Table S1**).

For Polymorph 2b designs, two rounds of interface-focused refinement were performed. In round 1 (rd1), no constraints were imposed on interface residue composition. In round 2 (rd2), constraints were introduced to prohibit methionine and glycine at the interface while limiting phenylalanine, tryptophan, and tyrosine to a maximum of 10% for each to reduce hydrophobicity and potential aggregation. A selection of designs from rd1 strategies (S6-S9; **Supplementary Table S1**) was used as input for rd2 optimization.

Comparing the interface composition between the two rounds, we observed an increase in polar and charged amino acids, with the largest increase in serine. That shift may explain the increase in hydrogen bonds at the interface in rd2 designs compared to rd1 designs. Finally, designs were selected based on Rosetta metrics and forwarded for experimental testing (**Supplementary Fig. S3**).

### Disulfide bond engineering using Disulfide by Design tool

Disulfide bonds were introduced into the designed binders to enhance structural stability using the Disulfide by Design tool^73^. Sequences incorporating cysteine residues at the predicted positions were evaluated using AlphaFold2^74^ (AF2) in single sequence mode with six recycles to assess *in silico* foldability. AF2 predictions of disulfide-stabilized variants yielded acceptable pLDDT scores ≥70 (AS 0763cc: 70, AS 1210cc: 87, and AS 2777cc: 95; **Supplementary Fig. S5**), whereas non-stabilized designs either failed to be predicted or did not match the Rosetta model. Additionally, C_α_ RMSD relative to Rosetta models was evaluated.

### Preparation of designs library for yeast surface display

A library of approximately 6,000 designs targeting the WT aSyn polymorphs (1a, PDB: 6CU7; 2b, PDB: 6SST) was reverse-translated into DNA sequences and synthesized as an oligo pool (Twist Bioscience). The oligo pool underwent two rounds of PCR: the first amplified the library, and the second incorporated 45 base pair homology regions specific to the pCTcon2 vector backbone (Addgene, 41843). The resulting DNA fragments were mixed with linearized pCTcon2 vector at a 5:1 insert-to-vector ratio and transformed into EBY-100 yeast via electroporation, following established protocols^132^. Transformed yeast cells were cultured in minimal glucose medium (SDCAA) at 30°C, passaged at least once, and induced in minimal galactose medium (SGCAA) overnight at 30°C.

### Flow cytometry analysis and sorting of designs library

Induced yeast cells were processed at 4°C in PBS containing 0.1% BSA as the reaction buffer. Cells were washed, incubated with sonicated WT aSyn fibrils for 2 h, and washed once again prior to antibody labeling. A primary antibody against aSyn C-terminus (in-house prepared; LASH-EGT405^133^; epitope: 113-127; 1:100 dilution) was applied for 45 minutes. Next, cells were pelleted and labeled with a FITC-conjugated goat anti-HA tag antibody (Bethyl; A190-138F; display tag; 1:100 dilution) and Alexa Fluor™ 647-conjugated donkey anti-mouse IgG (Invitrogen; A-31571; binds 1^ry^ antibody; 1:200 dilution) for 30 minutes. Labeled cells were pelleted, resuspended in buffer, and analyzed on a CytoFLEX S flow cytometer (Beckman Coulter) or sorted using a FACSAria II cell sorter (BD Biosciences) into three populations: non-binders (negative), potential binders (dim), and strong binders (high) based on Alexa Fluor™ 647 fluorescence intensity (**Supplementary Fig. S26**). Sorted cells were grown in SDCAA medium, and the strong binder population was prepared similarly for an additional sorting round. The target concentration was reduced between rounds (sort I: 1 μM; sort II: 0.5 μM) to select for higher-affinity binders.

### Post-sort processing and next-generation sequencing (NGS)

After sort II, yeast cells were cultured in SDCAA medium, pelleted, and plasmid DNA was extracted using the Zymoprep Yeast Plasmid Miniprep II kit (Zymo Research) following the manufacturer’s instructions. Library preparation for NGS involved two sequential PCRs: the first amplified coding sequences for the designed proteins using vector-specific primers flanking the coding sequences for the designed proteins, and the second attached Illumina sequencing adapters and Nextera barcodes. PCR products were purified and desalted using either the QIAquick PCR purification kit (Qiagen) or AMPure XP beads (Beckman Coulter). Sequencing was performed on an Illumina MiSeq platform, generating 0.6–1.5 million reads per sample. Bioinformatics analysis included translating sequences in the correct reading frame, matching them to expected input sequences from the library, and calculating enrichment of each design by dividing counts in the binding population by those in the non-binding population. Hits were identified as designs with enrichment values in the binding population exceeding 10-fold compared to the non-binding population (**Supplementary Table S2**).

### Yeast surface display of individual designs

Genes encoding individual designs were obtained from Twist Bioscience with homology overhangs compatible with pCTcon2 (Addgene, 41843) for Gibson assembly cloning. Constructs were transformed into HB101 competent bacteria for plasmid amplification, followed by sequence verification using Sanger sequencing. Verified plasmids were transformed into competent EBY-100 yeast using the Frozen-EZ Yeast Transformation II Kit (Zymo Research) following the manufacturer’s instructions. Transformed yeast cells were cultured in minimal glucose medium (SDCAA) at 30°C, passaged at least once, and induced overnight in minimal galactose medium (SGCAA) at 30°C. Induced yeast cells were prepared for flow cytometry analysis as detailed in the “*Flow cytometry analysis and sorting of designs library*” section. For experiments conducted at room temperature (RT), incubation times were adjusted as follows: yeast cells were incubated with target fibrils for 30 minutes, followed by 15-minute incubation for each antibody labeling step. Labeled cells were analyzed using a CytoFLEX S flow cytometer (Beckman Coulter), and data were processed with FlowJo (BD Biosciences, v.10.8.1).

### De-novo designs expression and purification

Protein sequences used in this study are listed in **Supplementary Table S4**. DNA sequences encoding lead candidates were synthesized by Twist Bioscience and cloned into pET11 vector for bacterial expression or pHLSec vector for mammalian expression via Gibson assembly. Constructs were transformed into HB101 competent bacteria for plasmid amplification, extracted using GeneJET Plasmid Miniprep Kit (Thermo Fisher Scientific, K0503) or Plasmid Plus Midi Kit (Qiagen, 12945), and verified by Sanger sequencing.

Bacterial expression was performed in BL21 (DE3) or T7 Express Competent *E. coli* cells with IPTG induction (1 mM at OD600 0.6-0.8), followed by overnight growth at 18°C. Harvested cells were pelleted by centrifugation at 4,000g for 20 minutes at 4°C, resuspended in lysis buffer (50 mM Tris, pH 7.5, 500 mM NaCl, 5% glycerol, 1 mg/mL lysozyme, 1 mM PMSF, and 1 µg/mL DNase), and lysed via sonication. Lysates were clarified by centrifugation at 30,000g for 30 minutes and filtered through 0.45 µM filters.

Mammalian expression was performed using the Expi293 system (Thermo Fisher Scientific, A14635), with supernatant collected 6 days post-transfection and filtered through 0.45 µM filters. Bacterial lysates and mammalian supernatants containing 6xHis-tagged proteins were purified by Ni-NTA HisTrap affinity chromatography, followed by size-exclusion chromatography using a Superdex HiLoad 16/600 200 pg column on an ÄKTA pure system (GE Healthcare). Proteins were concentrated in PBS, aliquoted, and stored at −80°C until use.

Biophysical characterization of the expressed GFP-fusion proteins including our binders (AS-0763cc and AS-2777cc), their scaffold-only controls, NbSyn2 and GFP alone were monomeric and of the expected molecular weight (**Supplementary Fig. S27**).

### Size-exclusion chromatography–multi-angle light scattering (SEC-MALS)

The molecular weight and oligomeric state of purified proteins were determined using size-exclusion chromatography coupled with multi-angle light scattering (SEC-MALS). Chromatography was performed on a Superdex 75 300/10 GL column (GE Healthcare) at a flow rate of 0.5 mL/min, controlled by Chromeleon software (Thermo Fisher Scientific, v7.2.10). A 100 µl sample of purified protein (∼1 mg/mL) was injected, and signals for ultraviolet absorbance at 280 nm, refractive index (dRI), and light scattering were recorded using a miniDAWN TREOS detector (Wyatt). Molecular weight calculations were performed with ASTRA software (v6.1, Wyatt).

### Dynamic light scattering (DLS) measurements

Designs were diluted to 10 µM in PBS and loaded into the standard-grade capillaries for DLS measurements using the Prometheus Panta instrument (NanoTemper, Germany). The hydrodynamic radius was calculated from normalized mass-weighted distributions of sample populations obtained from the size distribution fit of the autocorrelation function (ACF). For each sample, ten ACFs were recorded from two independent capillaries.

### Expression and purification of human WT aSyn monomers

Human WT aSyn monomers were expressed and purified using two protocols^75,134^ to be used mainly for ThT-aggregation kinetic assays. Briefly, BL21(DE3) *E. coli* cells were transformed with pT7-7 plasmids encoding the respective gene (Addgene; 36046) and plated on ampicillin-containing agar plates. Transformed cells were cultured in Luria broth (LB) medium supplemented with ampicillin (50-100 μg/mL) and induced with IPTG (1 mM) at OD600 0.6–0.8, followed by incubation at 37°C and 180 rpm for 3-4 hours. Harvested cells were pelleted by centrifugation at 5,000g (protocol 1) or 4,000 rpm (protocol 2) for 30 min at 4°C. Pellets were resuspended in lysis buffer (protocol 1^75^: 10 mM Tris-HCl, pH 8.0; protocol 2^134^: 40 mM Tris-HCl, pH 7.4) containing 1 mM EDTA and 1 mM PMSF, then lysed using probe ultrasonication for 8 min. Insoluble fractions were removed by centrifugation at 20,000g (protocol 1) or 12,000 rpm (protocol 2) for 30 minutes at 4°C. The supernatant was boiled for 15–20 minutes, and heat-precipitated proteins were removed by centrifugation.

In protocol 1^75^, aSyn (WT, E46K, H50Q, or K43Q/K45Q) was precipitated by the addition of saturated ammonium sulfate solution ((NH_4_)_2_SO_4_; 4 mL per 1 mL of clarified supernatant), incubated for 15 min at 4 °C on a rocking platform, and centrifuged to collect the protein pellet. The pellet was dissolved in 25 mM Tris–HCl, pH 7.7 with 1 mM DTT and dialyzed against the same buffer for 16–18 h with a buffer exchange after 12 h at 4 °C. The dialyzed protein was purified using anion exchange chromatography (AEC) (HiTrap Q Hp, 5 mL) followed by size exclusion chromatography (SEC) (HiLoad 16/600 Superdex 200 pg column) on an ÄKTA System (GE Healthcare). Monomeric aSyn fractions eluted in 10 mM of sodium phosphate buffer (pH 7.4) were collected, snap frozen and stored at -80°C.

In protocol 2^134^, The clarified supernatant was filtered through 0.45-μm filters and purified by AEC (HiPrep Q Fast Flow 16/10) on an ÄKTA System (GE Healthcare). Purified fractions were further processed either via reverse-phase HPLC (high-performance liquid chromatography) using a C4 column (PROTO 300 C4, 10 μm, Higgins Analytical) with buffer A (0.1% trifluoroacetic acid (TFA) in water) and buffer B (0.1% TFA in acetonitrile) or by SEC using a Superdex HiLoad 16/600 200 pg column. For HPLC, proteins were eluted using a 35-45% gradient of buffer B over 40 minutes at 15 mL/min, and purity was confirmed by ESI-MS before pooling, snap-freezing, and lyophilization. For SEC, monomeric fractions eluted in PBS (pH 7.4) were collected, snap frozen, and stored at −80°C.

### ESI-MS analysis

Electrospray ionization mass spectrometry (ESI-MS) analyses were performed using LC-MS on an LTQ system (Thermo Scientific, San Jose, CA), as previously described in [^135^]. Proteins were desalted online via reversed-phase chromatography using a Poroshell 300SB C3 column (1.0 × 75 mm, 5 μm; Agilent Technologies, Santa Clara, CA) integrated with the LTQ system. Approximately 10 μl of each protein sample was injected at a flow rate of 300 μl/min. Proteins were eluted with a linear gradient of 5% to 95% solvent B against solvent A. Solvent A consisted of 0.1% formic acid in ultrapure water, while solvent B was 0.1% formic acid in acetonitrile. Charge state deconvolution and MS data analysis were performed using MagTran software (Amgen Inc., Thousand Oaks, CA).

### Preparation of monomeric aSyn

HPLC-purified lyophilized aSyn was reconstituted in PBS and filtered through 100-kDa spin filters to isolate monomeric aSyn in the filtrate, following a previously described protocol^134^. For concentration adjustment or buffer exchange, 3-kDa spin columns were used. The concentration of monomeric aSyn was measured using a NanoDrop spectrophotometer at 280 nm with an extinction coefficient of 5960 M^-1^ cm^-1^, calculated using the protein sequence (ProtParam, ExPASy). The quality of monomeric aSyn was verified via mass spectrometry (ESI-MS) and size-exclusion chromatography coupled with multi-angle light scattering (SEC-MALS) (**Supplementary Fig. S28**).

### Preparation of oligomers and fibrils from aSyn and Tau

#### Preparation of WT human aSyn fibrils for *in vitro* and cell culture studies

The aSyn fibrils polymorphs (Fibrils, Ribbons, and Fibrils-91) were assembled as previously described [^15,62,63^]. The aSyn monomers prepared as described in [^62,63^] were first dialyzed against the appropriate buffer corresponding to the polymorph to produce (Fibrils: 50 mM Tris-HCl pH 7.5, 150 mM KCl; Ribbons: 5 mM Tris-HCl pH 7.5; F91: 25 mM Na_2_PO_4_ pH 9.1) and incubated at 37°C under shaking (600 rpm) for 5 to 10 days depending on the polymorph.

aSyn fibrillization was measured by withdrawing 10 µl from three independent assembly reactions ran in parallel and diluting them at different time intervals into 390 µl of ThT (10 µM in water) as previously described in [^62^]. The increase in ThT fluorescence was assessed using a Cary Eclipse fluorescence spectrophotometer (Agilent, Les Ulis, France) set at 440nm and 480nm for excitation and emission, respectively.

Assembled polymorphs were ultracentrifuged to remove soluble aSyn from the solution (30 min, 35,000g) and then resuspended in sterile PBS to reach a final concentration of 350 µM (monomer equivalent). The morphology of the polymorphs was verified by transmission electron microscopy after negative staining with 1% uranyl acetate (Jeol 1400 TEM, Gatan Orius CCD camera) and the proteolytic profiles of the assemblies were verified by limited proteolysis with Proteinase K (3.8 µg/mL), as previously described in [^15,62,63^].

For *in vitro* experiments, the polymorphs were fragmented by sonication using a sonotrode (20 min, 0.5s pulses, VialTweeter probe powered by Sonicator UIS205V, Hielscher Ultrasonic, Teltow, Germany), then aliquoted (6 µL, 350 µM), flash-frozen in liquid nitrogen and stored at -80°C until use. Before use, the polymorphs were thawed at 30°C, diluted to the desired working concentration in PBS and stored at 30°C while in-use for a period no longer than five days.

For cell culture experiments, Fibrils and Ribbons were split into two batches. One batch was then labelled with ATTO647 by incubating the polymorphs with ATTO647 NHS-Ester (AD 647N, ATTO-TEK, Germany) for 45 min at RT at a molar ratio aSyn (equivalent monomers) to ATTO647 NHS-Ester of 1: 1. At the end of the incubation, unbound ATTO647 NHS-Ester was inactivated by addition of Tris-HCl pH 7.5 (final concentration of 40 mM) and incubation for 10 minutes. Labeled polymorphs were then ultracentrifuged to eliminate unbound ATTO647 dye and buffer (30 min, 35,000g) and were resuspended in sterile PBS at a final concentration of 350 µM. Labeled and unlabeled polymorphs were then fragmented by sonication as indicated above, aliquoted, flash frozen and stored at -80°C. Polymorphs were thawed at 37°C and used immediately.

#### Preparation of E46K, H50Q and K-to-Q aSyn mutant fibrils

Monomers of aSyn carrying E46K, H50Q, or K-to-Q mutations were expressed and purified according to protocol 1 described above. Each monomer (100 µM) was exchanged into 50 mM Tris-HCl buffer pH 7.5 containing 150 mM KCl and incubated at 37 °C for four weeks under constant agitation (600 rpm). The resulting fibrils were harvested by centrifugation (90 min, 16,000g, 25 °C), resuspended by the same buffer to 100 µM (monomer equivalent), sonicated two times (VialTweeter sonicator, 100% amplitude, 1s pulse / 1s pause, 4 min total time), flash frozen in liquid nitrogen, and stored at -80°C until further use.

#### Preparation of mouse aSyn fibrils

The mouse aSyn fibrils (5 mg/mL) used in this study were obtained from ND Biosciences, prepared as described in [^78^]. The product information of the utilized material is as follows: ND-05/23-037.001.

#### Preparation of Tau fibrils

The Tau fibrils used in this study were produced as described in [^83^]. The fibrils were prepared as 6 µl of 80 µM stock solution. The stocks were then diluted to the desired working concentration using PBS and stored at 30°C while in-use for a period no longer than five days.

#### Preparation of aSyn oligomers

The aSyn oligomers (WT and HNE-modified) used in this study were obtained from ND Biosciences and were prepared as described in [^84^].

The product information of the utilized material is as follows:

- WT aSyn oligomers (lyophilized Spontaneous aSyn oligomers; 100 µg; ND-11/21-002.001.1)
- HNE (4-hydroxy-2-nonenal) crosslinked aSyn oligomers (2 µM; ND_69_HUM_ASYN_HNE_OLIGO – POOLA3A8 – 10/20.005 002)

The concentration of HNE oligomers was measured using a NanoDrop spectrophotometer at 280 nm with an extinction coefficient of 5960 M^-1^ cm^-1^, corresponding to aSyn protein. Lyophilized spontaneous aSyn oligomers were resuspended in PBS to a final concentration of 1 mg/mL.

### ThT binding kinetics and residual monomer quantification

The effect of lead candidate designs and their knock-out mutants on the elongation of WT aSyn seeds was evaluated using ThT fluorescence assay (excitation/emission = 450/482 nm). Reaction mixtures contained 0.5 µM sonicated WT aSyn fibrils (seeds), 20 µM fresh aSyn monomer, 20 µM ThT, and varying concentrations of designs or their scaffold-only controls, all prepared in PBS. Assays were performed in triplicate (15 µl/well) using 384-well low-volume, non-binding black plates with a clear bottom (Corning, 3544) sealed using adhesive sealing sheets (ThermoScientific) under quiescent conditions at 37°C. ThT fluorescence was recorded from the bottom of the wells using a FLUOstar Omega reader (BMG LABTECH).

The residual monomer at the end of the aggregation reaction was quantified by pooling triplicate samples, centrifuging at 16,000g for 1 h at 25°C, and carefully decanting the supernatant for analysis by SDS-PAGE. Native aSyn monomer standards at 40, 20, and 10 µM were included on each gel to quantify residual monomer concentrations at the reaction endpoint. Gels were stained with InstantStain Coomassie Stain (INST-1L-181, Kem-En-Tec Nordic A/S, Denmark), imaged using a ChemiDoc Go system (BioRad, USA), and analyzed with Image Lab software (BioRad, USA).

### Isothermal titration calorimetry

Proteins were purified, resuspended or dialyzed against assay buffer (PBS [-Mg^2+^, -Ca^2+^], pH 7.4) overnight using Slide-A-Lyzer™ MINI Dialysis Devices, 3.5K MWCO (Thermo Fisher Scientific; 88400). ITC data were collected with an MicroCal PEAQ-ITC Automated instrument (Malvern Panalytical) at 25 °C, 10 µcal/s reference power, 750 rpm. stirring speed, and high gain with 18 2-µl injections at 180-sec intervals. Titrations were performed by titrating 150 µM Synuclein into 10 µM designed binders and binding-controls or into buffer as experimental control.

Baseline correction, peak integration, control subtraction, and data fitting were conducted using the MicroCal PEAQ-ITC Analysis Software (v 1.41) and the initial injection was not included for data analysis. Data were fit to a single-site binding model to derive *K*_D_, Δ*H* (change in binding enthalpy), and *N* (stoichiometry) values from the binding isotherm.

### Flow-Induced Dispersion Analysis (FIDA)

The non-diffusive profile of aSyn oligomers was evaluated using their intrinsic fluorescence via FIDA on a Fida1 instrument (FidaBio, Denmark) equipped with a 280 nm absorbance detector.

Binding profiling of GFP-fused designs against sonicated fibrils and oligomers was performed using FIDA on the same instrument but equipped with a 488 nm fluorescence detector. GFP-fused designs were prepared as a 200 nM stock solution in PBS (2x working concentration), and binding targets were serially diluted to 2x working concentration in PBS. Samples for analysis were prepared by equally mixing binders and targets.

FIDA samples were introduced into 1m-long capillary with inner diameter of 75 μm, applying the following injection method:

1. Wash 1 (1 M NaOH): 45 s, 3500 mbar.
2. Wash 2 (MQ water): 45 s, 3500 mbar
3. Equilibration (PBS): 30 s, 3500 mbar.
4. Sample application (GFP-fused designs/binding target): 20 s, 75 mbar.
5. Measurement and detection (PBS): 75 s, 1500 mbar.

In the assay, the pre-mixed binder/fibril sample (∼100 nL) undergoes transient incomplete separation^75^ in a laminar flow yielding a diffusion profile consisting of transient peak (or spikes) and gaussian distribution, corresponding to the non-diffusive (binders bound to fibrils) and diffusive (free binders) particles, respectively. Capillary adhesion of GFP-labeled species hindered quantification of free binders. Therefore, binding to fibrils was assessed by quantifying the signal of bound species from the non-diffusive peak area as described in [^75^] or using Simpson’s Rule Cumulative Integration^77^ for fibrils eluting as spikes within the elution time window of 0.18–0.25/0.3 min. Titration curves were generated by plotting integrated peak areas (Y-axis) against binding target concentrations (X-axis). The binding affinity (apparent *K_D_*) of AS-2777cc for aSyn Ribbons polymorph was determined by fitting saturating titration curves using nonlinear four-parameter curve analysis in GraphPad Prism (v 10.4.1).

### Cell culture experiments

Murine neuroblastoma cells (Neuro2A, ATCC, Manassas, VA) and HeLa cells (Sigma Aldrich, Saint-Quentin-Fallavier, France) were maintained in modified DMEM medium (Dulbecco’s Modified Eagle Medium) containing glucose and GlutaMAX (Gibco, 6196-026), supplemented with 10% fetal bovine serum (Gibco, 10270-106), 100 U/mL of penicillin and 100 µg/mL of streptomycin (Gibco, 10270-122). Cells were cultured at 37°C in humidified air with 5% CO_2_. For internalization and seeding assays, cells were plated in eight-well ibiTreat µ-Slides (Ibidi, 80826).

#### Internalization assay

Both HeLa and undifferentiated Neuro2A cells were used for internalization assays. ATTO647-labeled aSyn polymorphs (Fibrils or Ribbons) and GFP-fused binders were thawed and diluted in sterile PBS to a final concentration of 10 µM each. Binders and fibrils were mixed and incubated for 16 h at RT, protected from light. For conditions without fibrils, binders were diluted in sterile PBS to 10 µM and incubated under identical conditions. After incubation, mixtures were diluted to 2 µM in fresh pre-warmed modified DMEM medium. Half of the culture medium was replaced with medium containing either fibril-binder or PBS-binder mixtures to achieve final concentrations of 1 µM fibrils and 1 µM binders in the culture plate. Cells were then incubated for 2 h at 37°C.

Following exposure, the culture medium was removed, and cells were washed twice with warm phenol red-free medium and maintained in this medium at 37°C in a humidified incubator with 5% CO_2_ during live imaging on a fluorescence microscope (Axio Observer Z1, Zeiss). Imaging positions (10 per condition) were selected in a blinded manner using phase-contrast images and coordinates were recorded for subsequent acquisition. Trypan blue (0.4% stock, Gibco, 15250-061) was added before imaging at a final concentration of 0.14% to quench extracellular GFP fluorescence. Cells were then immediately imaged in Cy5, FITC, and phase contrast channels, at 20X magnification, using the pre-defined acquisition coordinates.

Blind coded image analysis was performed using ImageJ (v1.54p). Pixel-based colocalization analysis was conducted on 8-bit grayscale images using the Coloc2 plugin (v3.1.0, https://imagej.net/plugins/coloc-2), generating scatterplots and Pearson’s correlation coefficients. Artefacts, including overlapping cells (predominantly in Neuro2A cultures) and dead cells, were excluded by manual region-of-interest selection based on the Cy5 channel. Object-based colocalization analysis was performed using the 2D Object Colocalization macro for FIJI (https://github.com/dsrichardson/fiji_macros), which identifies objects based on local intensity maxima and assesses their spatial overlap between channels. Colocalization was expressed as the percentage of Cy5-positive objects colocalizing with FITC-positive objects per image, and mean values were calculated for each condition. Experiments were performed once in Neuro2A cells and once in HeLa cells. Graphs were generated using prism GraphPad (v 10.6.1).

#### Seeding assay

To assess the effect of binders on polymorph-induced seeding in cell culture, we used a HeLa cell line stably expressing human full-length WT aSyn fused to mCherry fluorescent protein (HeLa aSyn-mCherry). HeLa cells were transfected with an integrative plasmid (pCDNA 3.1) encoding the sequence for aSyn fused to mCherry fluorescent protein (the SNCA mCherry sequences are integrated between the Hind III and Bgl II restriction enzymes sites) using lipofectamine 2000 following the manufacturer instructions^62,136^. Transfected cells were selected in presence of 400 µg/mL of G418 for 3 weeks and sorted on the basis of their levels of aSyn-mCherry expression by FACS (FACSAria III, BD Biosciences).

For seeding assays, unlabeled aSyn polymorphs and GFP-fused binders were mixed and incubated 16 h prior to cell exposure, as described previously for internalization experiments. HeLa aSyn-mCherry cells were then exposed for 24 h to a modified DMEM medium containing the fibril-binder mixtures at a final concentration of 1 µM for each component. Following exposure, culture medium was removed and cells were washed twice with warm sterile PBS, fixed for 10 min at RT in warm 4% paraformaldehyde (PFA) in sterile PBS, and washed four times with cold sterile PBS.

Images of fixed cells (10 fields per condition per replicate) were acquired by sequential frame acquisition on an inverted confocal microscope (Leica SP8 X) using a 63X oil-immersion objective and a white-light laser. mCherry was detected using excitation at 568 nm (AF568 channel), and GFP was detected using excitation at 488 nm (FITC channel). Experiments were performed in two independent replicates. Quantification was performed manually on blind coded images using ImageJ by counting mCherry-positive cells and cells containing mCherry-positive punctae. The percentage of mCherry-positive cells containing punctae was quantified for each field as the ratio of mCherry-positive cells with punctae to the total number of mCherry-positive cells. Mean values were then calculated across all fields analyzed for each replicate and condition. Data are presented as bar graphs generated using Graphpad Prism (v 10.6.1), showing individual replicate means, with error bars representing the standard deviation.

