## supplementary data for "De novo Design of Polymorph-Specific Binders Targeting α-Synuclein Fibrils"

### Supplementary material

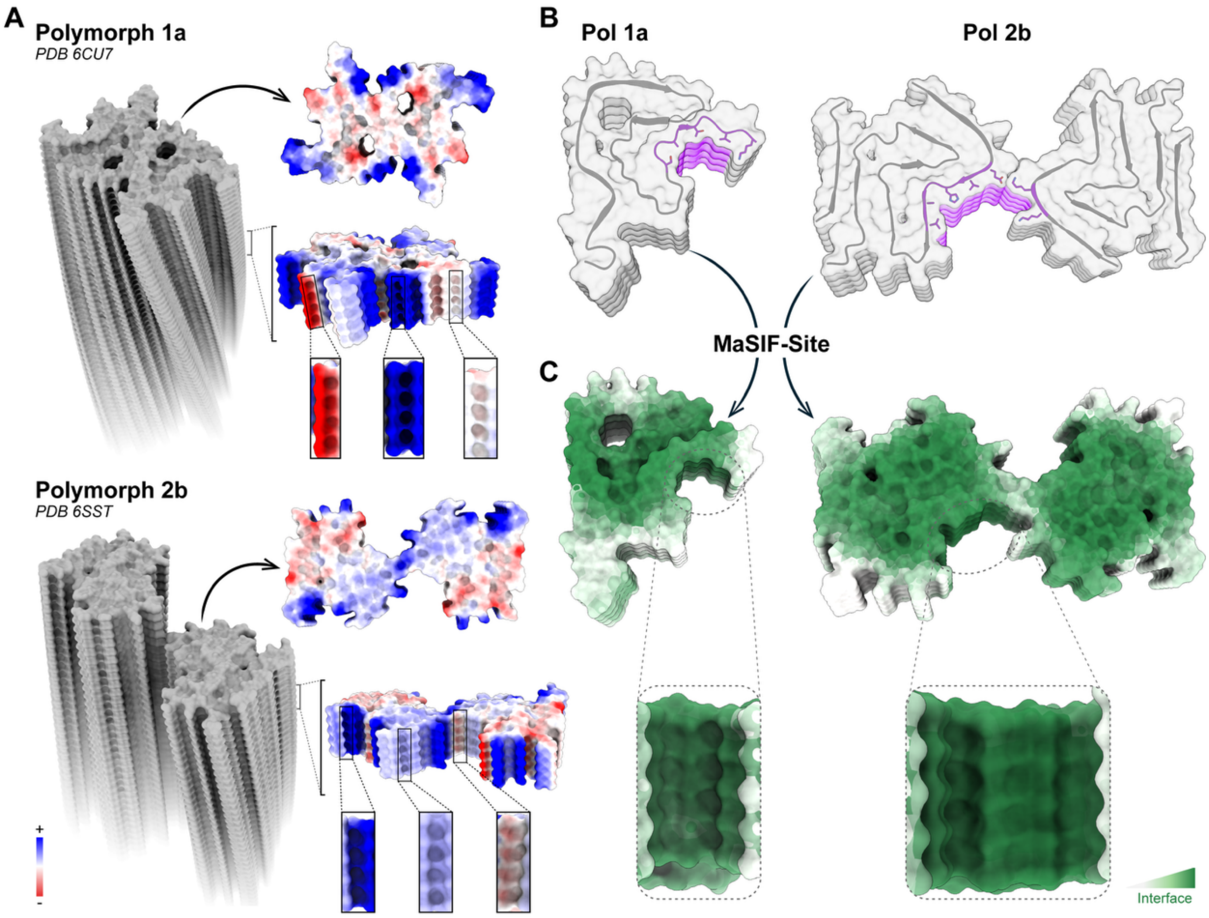


**Supplementary Figure S1 | Structural features of targeted aSyn polymorphs.**

**(A)** Structural models of aSyn Polymorphs 1a (PDB 6CU7; **top)** and 2b (PDB 6SST; **bottom)** fibrils showing surface electrostatic potential (red: negative; blue: positive). Insets highlight repetitive structural features, including sub-pockets along the lateral fibril surface. **(B)** Surface representations of both polymorphs with selected pockets highlighted in violet and pocket-forming residues shown as sticks. **(C)** MaSIF-site predictions of protein–protein interaction (PPI) potential for both polymorphs, with targeted pockets indicated by dashed circles, and zoomed-in side views illustrating their predicted capacity for PPI engagement (PPI sites: green; non-PPI sites: white).

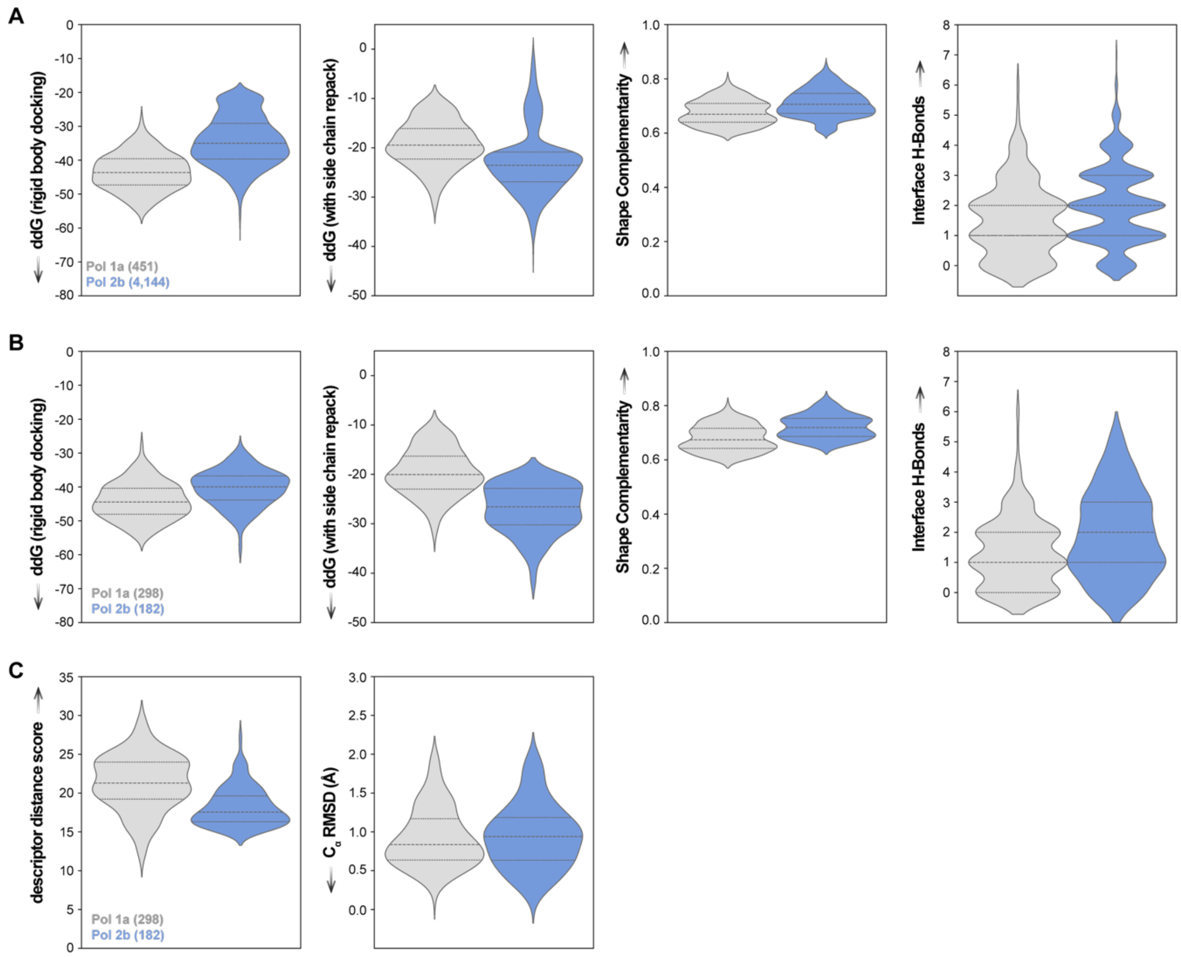


**Supplementary Figure S2 | Rosetta metrics and MaSIF-seach scores for selected *de novo* seeds.**

Violin plots showing Rosetta metrics for grid-selected *de novo* motifs (**A**) and MaSIF-search docked seeds selection (**B**), including binding energy (ddG), shape complementarity, the number of buried unsatisfied polar atoms, and interface hydrogen bonds. ddG was calculated under two conditions: with rigid-body docking (side chains fixed) and side-chain repacking. Scores are shown as violin plots for polymorph 1a (grey) and polymorph 2b (blue), with the number of designs displayed for each case. (**C**) MaSIF-search descriptor distance scores (references from Full set: 17.7; Patch matches with IPA score ≥ 0.8: 21.8; **Supplementary Fig. S25**) and $C_\alpha$ RMSD (≤2Å) to their respective Rosetta models for the docked seeds selection. Arrows mark the desired direction of improvement in metric values.

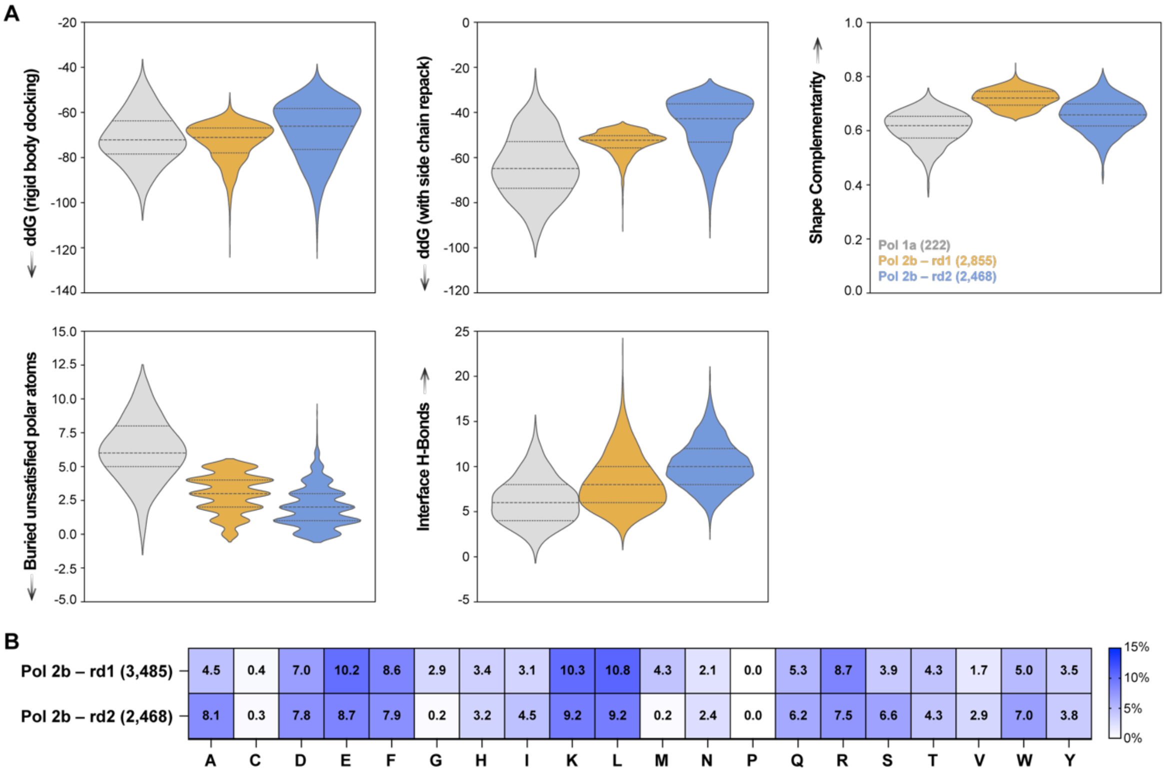


**Supplementary Figure S3 | Rosetta metrics for experimentally tested designs using yeast surface display.**

(**A**) Rosetta metrics for experimentally tested designs using yeast surface display, including Rosetta binding energy (ddG), shape complementarity, the number of buried unsatisfied polar atoms, and interface hydrogen bonds. ddG was calculated under two conditions: with rigid-body docking and side-chain repacking. Scores are presented as violin plots for polymorph 1a (grey, 222 designs) and polymorph 2b (orange, 2,855 rd1; and blue, 2,468 rd2 designs). (**B**) A heatmap compares amino acid composition in the polymorph 2b design interaction interfaces (≤5.5 Å from target) between rd1 and rd2, where rd2 constraints prohibited glycine and methionine at the interface while limiting phenylalanine, tryptophan, and tyrosine to a maximum of 10% each. Arrows mark the desired direction of improvement in metric values.

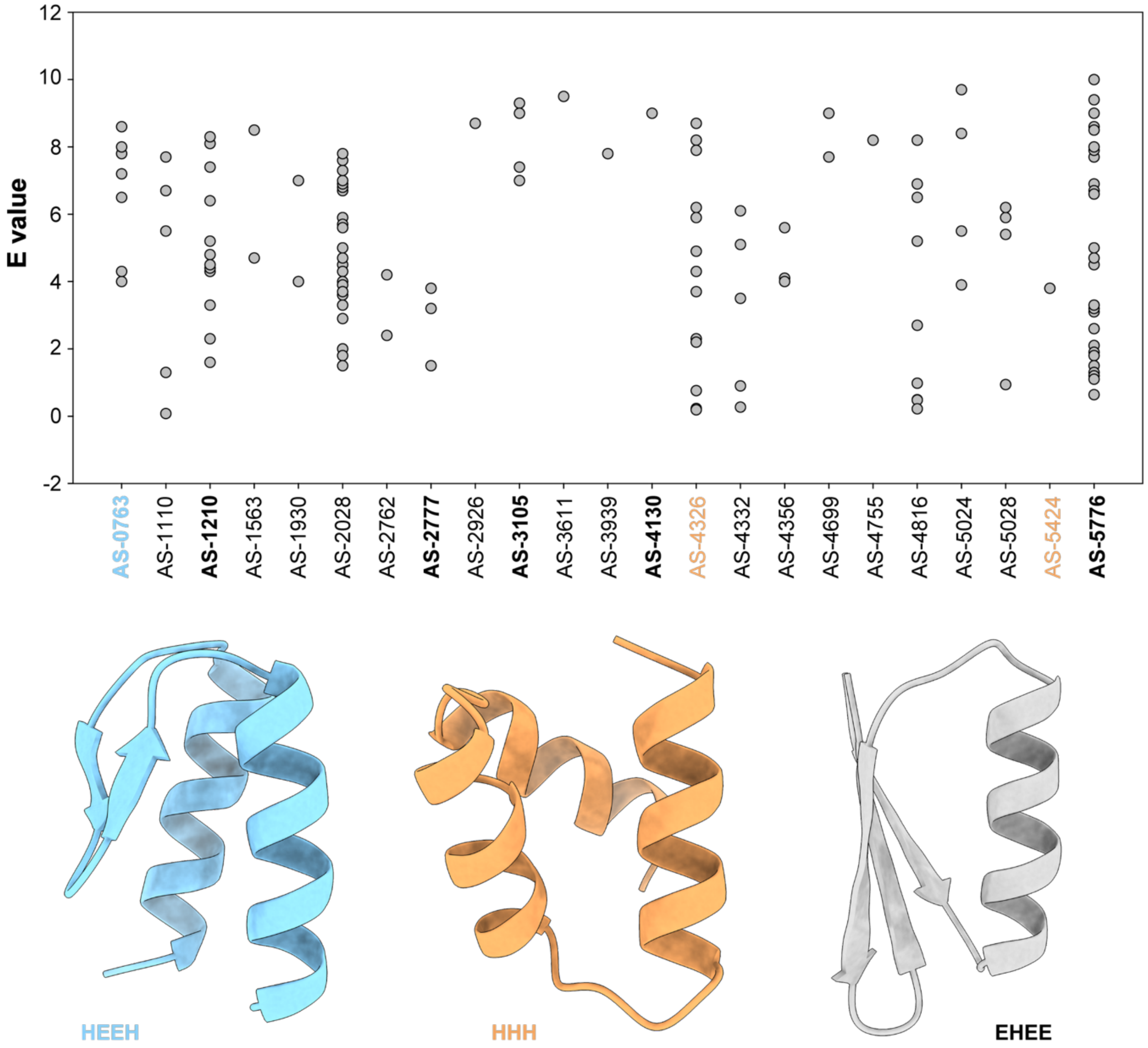


**Supplementary Figure S4 | Sequence Novelty and Fold Diversity of Selected Candidates**
The amino-acid sequences of 25 selected candidate binders (**Fig. 2A**) were queried against the SwissProt database using BLASTp[137,138]. All but two designs (AS-4763 and AS-5171) produced hits, shown as grey dots (one per hit), with E-values plotted on the y-axis. Nearly all hits showed positive E-values, indicating high sequence novelty and limited similarity to natural proteins. Candidates exhibited three structural folds (H: helix; E: beta-sheet): HEEH (blue), HHH (orange), and EHEE (grey). The design labels colors denote their respective fold, with experimentally validated designs highlighted in bold.

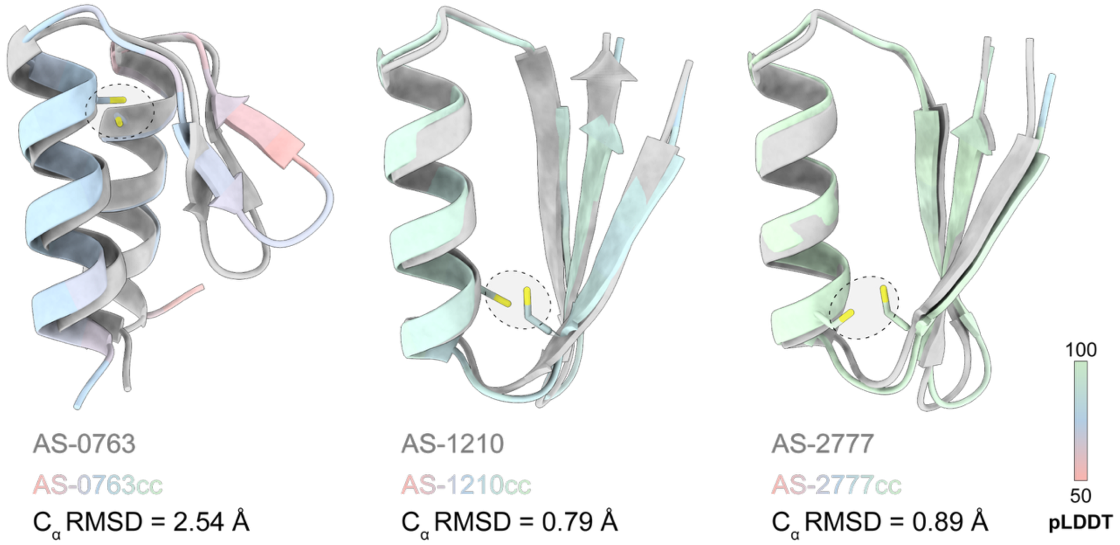


**Supplementary Figure S5 | Structural comparison of disulfide-stabilized designs**
Disulfide bonds were engineered using the Disulfide by Design tool[73], and the *in silico* foldability of the resulting sequences were evaluated using AlphaFold2 (AF2)[74]. Structural overlays compare AF2 models of the disulfide-stabilized designs (colored by pLDDT) with their respective Rosetta models (Grey) for AS-0763, AS-1210, and AS-2777. Engineered disulfide-bond cysteines are shown as sticks and highlighted with dashed circles. $C_\alpha$ RMSD was calculated using the UCSF ChimeraX MatchMaker tool (V 1.8) following model alignment.

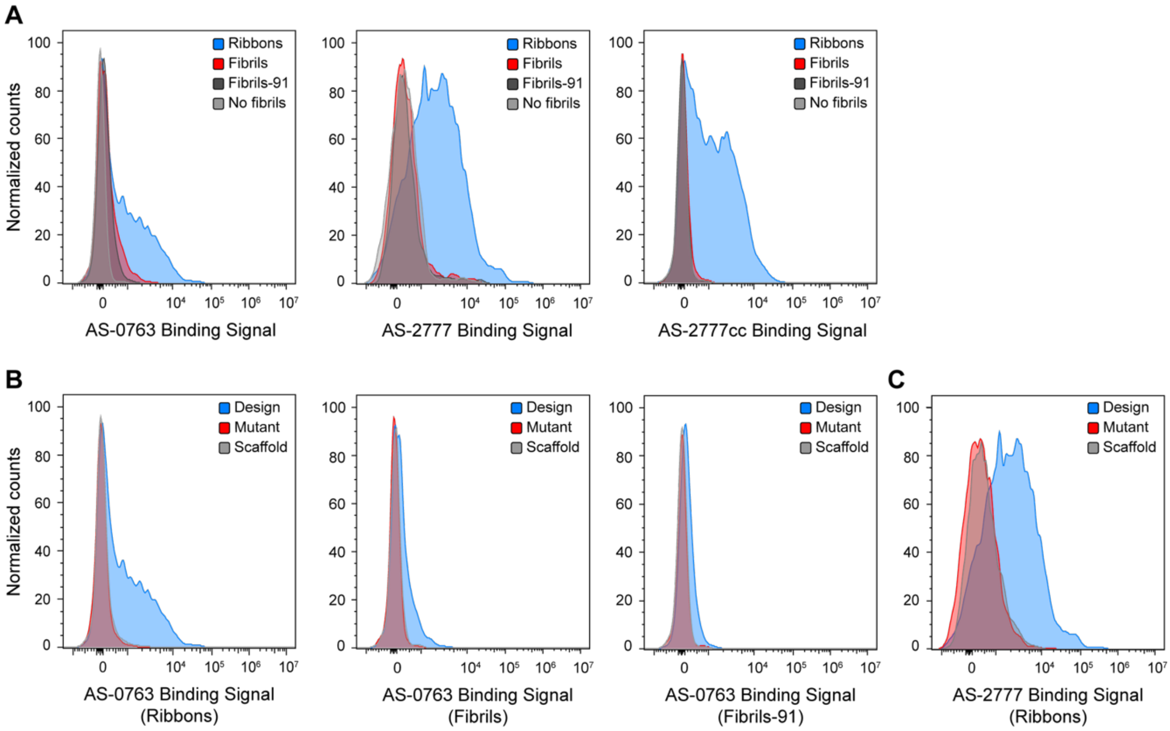


**Supplementary Figure S6 | Flow cytometry analysis of yeast-displayed designs against aSyn preparations.**

**(A)** Flow cytometry histograms showing binding of yeast-displayed AS-0763, AS-2777, and AS-2777cc to different *in vitro* aSyn fibril preparations (250 nM): Ribbons (blue), Fibrils (red), and Fibrils-91 (black). **(B, C)** Binding signals for designs AS-0763 **(B)** and AS-2777 **(C)** (blue), their respective interface knock-out mutants (red), and scaffolds (no interface; grey) against aSyn fibril preparations (250 nM). Yeast experiments were performed at 4°C as described in *Flow cytometry analysis and sorting of designs library* in the Methods section.

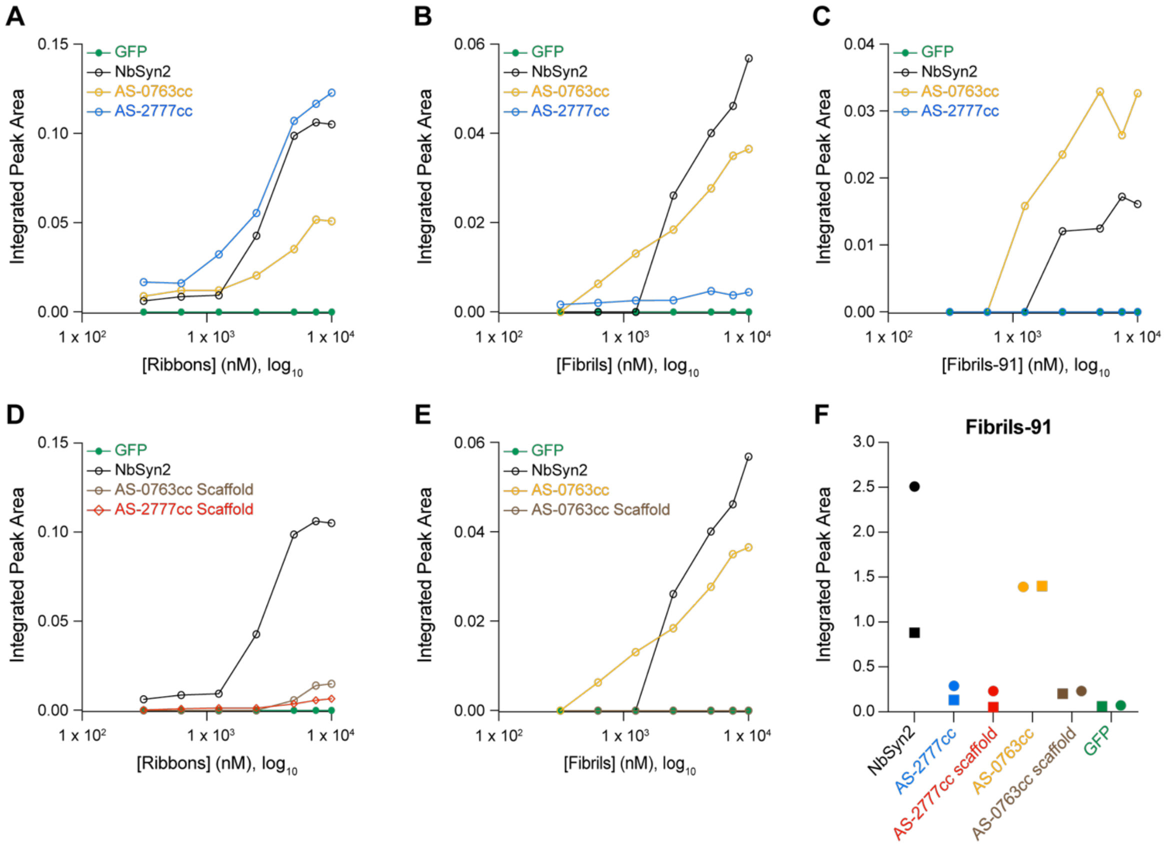


**Supplementary Figure S7 | Binding of GFP-fused designs to aSyn WT and mutant fibrils.**
**(A–C)** Titration binding curves of GFP-fused binders AS-0763cc (yellow), AS-2777cc (blue), NbSyn2 (positive control; black), and GFP (negative control; green), pre-mixed with aSyn fibrils and analyzed using FIDA1 equipped with a 488/525 nm fluorescence detector. **(A)** Binding to aSyn Ribbons, **(B)** binding to aSyn Fibrils, and **(C)** binding to aSyn Fibrils-91. These measurements are biological replicates of the titration curves reported in **Fig. 2D-F**. **(D)** Binding of knock-out mutants (scaffolds lacking designed interfaces), AS-0763cc scaffold (brown) and AS-2777cc scaffold (red), alongside NbSyn2 and GFP, to aSyn Ribbons. **(E)** Binding of AS-0763cc (yellow), its scaffold (brown), NbSyn2, and GFP to aSyn Fibrils. **(F)** Binding of GFP-fused designs, their respective scaffolds and controls to aSyn Fibrils-91 (5 μM). For all titration curves, the integrated peak area (y-axis), representing non-diffusive binding events, is plotted against fibril concentration (x-axis). Areas were estimated as described in [[75]] for titration curves (A-E) and panel (F). Individual measurements at each concentration are shown. For panel (F), each point represents a technical replicate (markers: Run I, circles; Run II, squares).

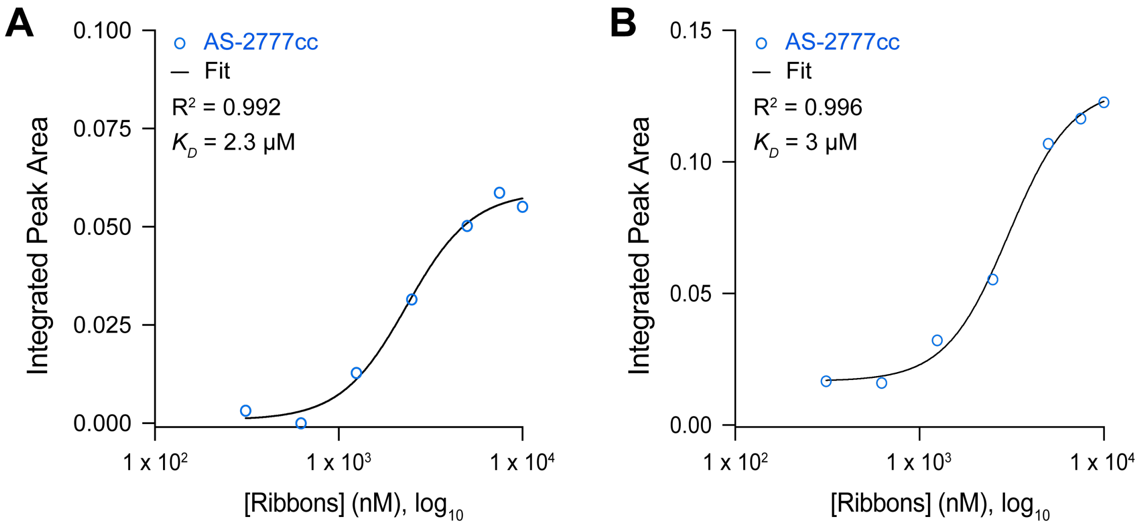


**Supplementary Figure S8 | Estimation of AS-2777cc binding affinity aSyn Ribbons preparation fibrils using FIDA.**

FIDA titration curves for aSyn Ribbons binding to AS-2777cc from two independent runs, previously reported in **Fig. 2D** (**A)** and **Supplementary Fig. S7A** (**B)**. Points represent areas of non-diffusive peaks, obtained as described in [[75]]. Data within the 0.312–10 µM concentration range were fitted using a nonlinear four-parameter curve analysis. Results from the two independent runs show strong agreement, indicating the low-micromolar affinity of AS-2777cc for aSyn Ribbons fibrils (apparent $K_D$ = 2.7 ± 0.3 µM).

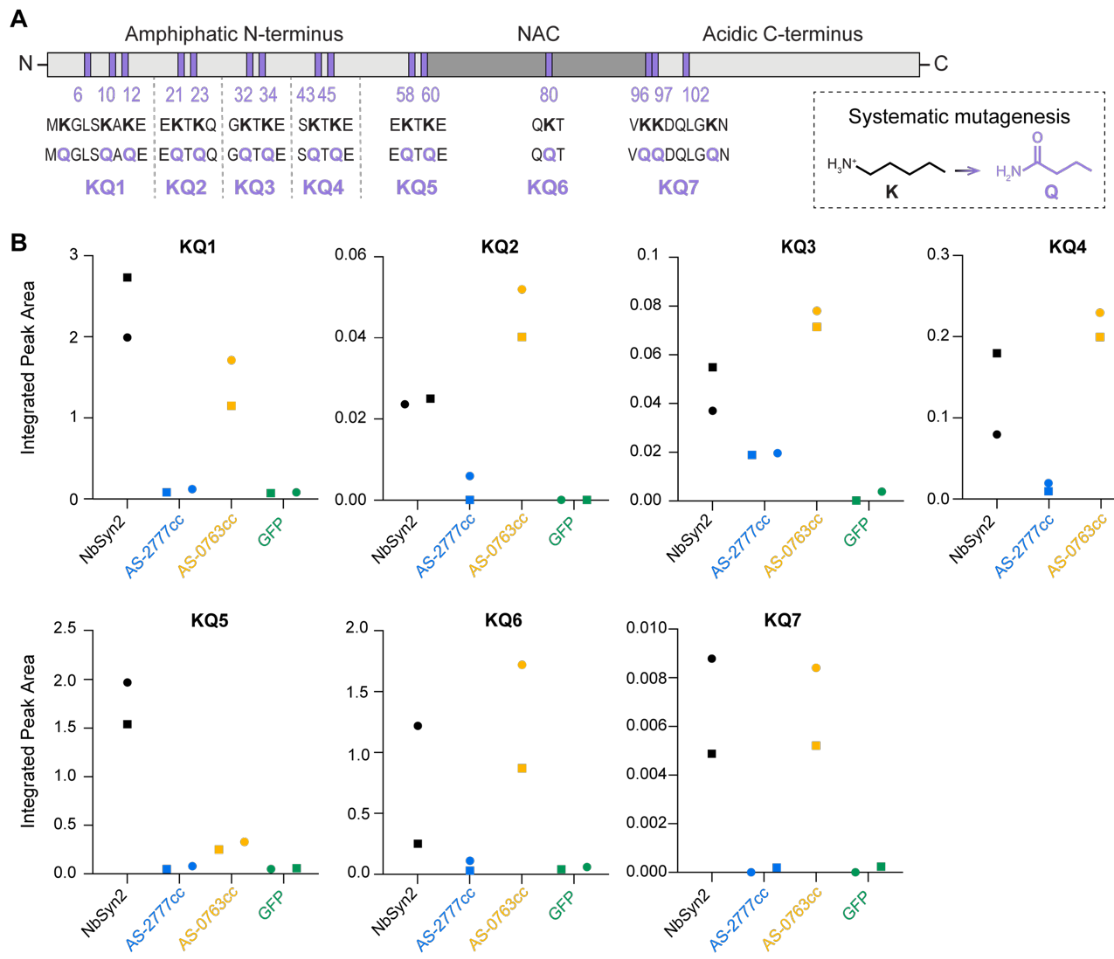


**Supplementary Figure S9 | Binding of GFP-fused designs to aSyn K-to-Q mutant fibrils measured by FIDA.**

**(A)** Schematic representation of the aSyn protein sequence with the tested K-to-Q mutants and their corresponding substitution sites indicated. **(B)** Binding of GFP-fused designs to a panel of aSyn K-to-Q mutant fibrils measured by FIDA. Integrated peak areas (y-axis), reflecting the fibril-binder complex, are compared to NbSyn2 (positive control) and GFP (negative control). Binder concentration was 100 nM, and K-to-Q fibrils were used at 2.5 µM, except for KQ4 at 5 µM. Each point represents an individual technical replicate (markers: Run I, circles; Run II, squares).

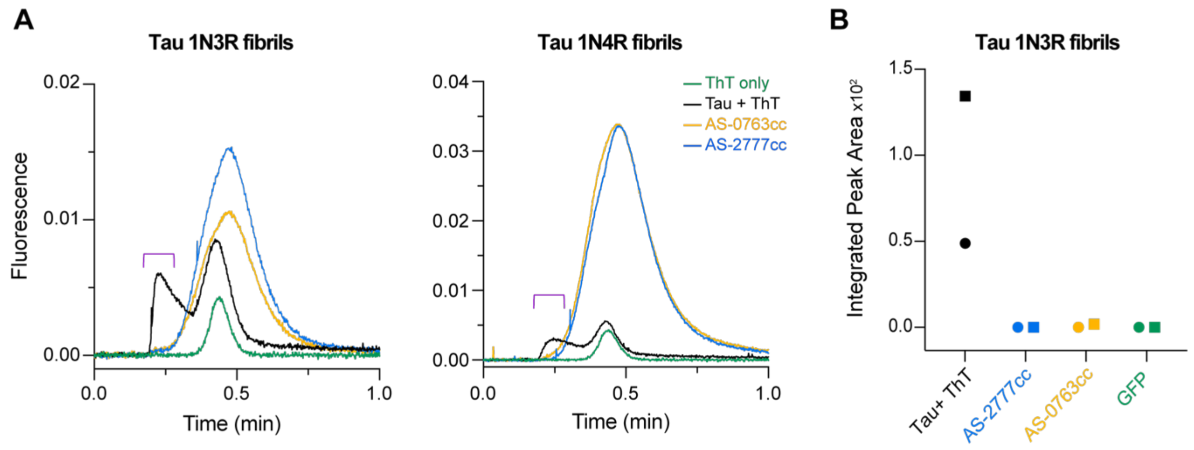


**Supplementary Figure S10 | Binding of GFP-fused designs to human WT Tau fibrils monitored using FIDA.**

**(A)** FIDA 488/525 nm Taylorgrams of pre-mixed GFP-fused designs (100 nM) or ThT (50 µM) with WT human Tau (4 µM): 1N3R fibrils (left) and 1N4R fibrils (right). The non-diffusive peak elution time for the fibril-binder complex is highlighted with a violet bracket, guided by Tau+ThT control. **(B)** Binding of GFP-fused designs to WT Tau fibrils 1N3R isoform was determined by integrating peak areas within the non-diffusive elution time as described in [75]. Integrated peak areas (y-axis) are compared to Tau+ThT (positive control) and GFP (negative control), with no measurable difference observed relative to GFP demonstrating binders' specificity towards aSyn fibrils. Each point represents an individual technical replicate (markers: Run I, circles; Run II, squares).

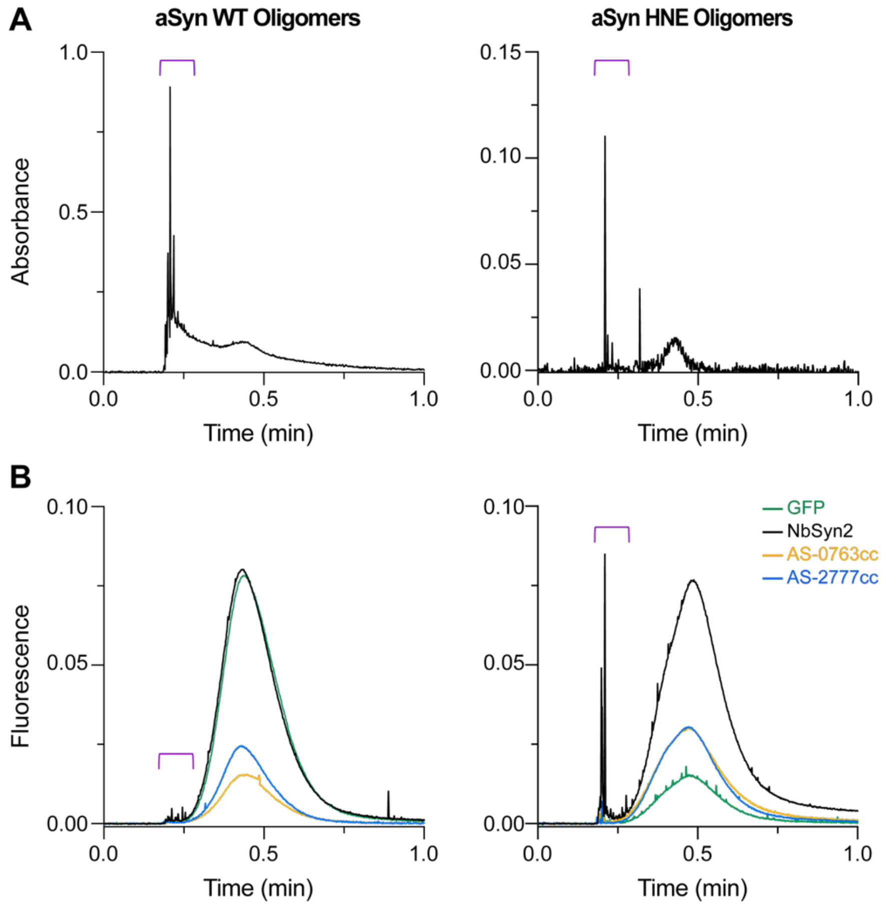


**Supplementary Figure S11 | Binding of GFP-fused designs to aSyn oligomers monitored using FIDA.** **(A)** FIDA $A_{280}$ nm Taylorgrams of aSyn oligomers: WT oligomers (left; 0.5 mg/ml) showing a transient curve in the elution profile indicative of particles with lower diffusion compared to aSyn monomers, and HNE-modified oligomers (right; 2 µM), displaying spikes in the elution profile consistent with large or higher-order assemblies. **(B)** FIDA 488/525 nm Taylorgrams of pre-mixed GFP-fused designs (100 nM) with aSyn oligomers: WT oligomers (left; 0.5 mg/ml) and HNE-modified oligomers (right; 1 µM). The non-diffusive peak corresponding to the oligomer or oligomer-binder complex is marked with a violet bracket.

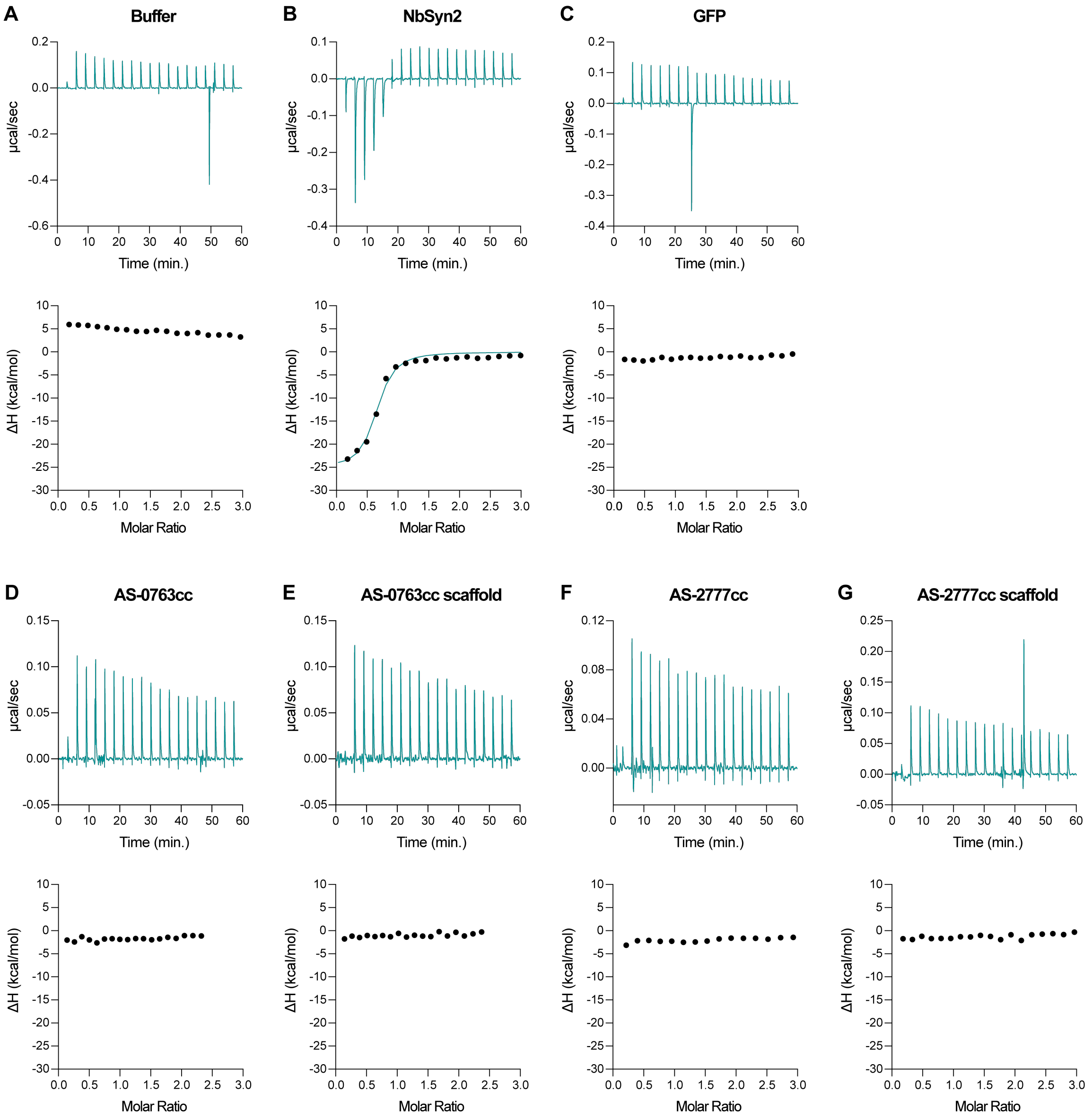


**Supplementary Figure S12 | ITC Analysis of WT aSyn monomer binding to designed proteins and controls.**

Isothermal titration calorimetry (ITC) experiments were performed with aSyn monomer titrated into **(A)** only buffer, **(B)** NbSyn2 (positive control), **(C)** GFP (negative control), and the designed proteins and their knock-out mutants: **(D)** AS-0763cc, **(E)** AS-0763cc scaffold, **(F)** AS-2777cc, and **(G)** AS-2777cc scaffold. The heat of dilution observed when titrating aSyn into only buffer (A) was subtracted from all experiments. The results confirm binding of NbSyn2 to aSyn monomer ($K_D$ = 338 nM, $N$ = 0.62, $\Delta H$ = -25.4 kcal/mol), while no binding was detected for the designed proteins, their knock-out mutants, or GFP under these experimental conditions ($K_D$ > 10 µM).

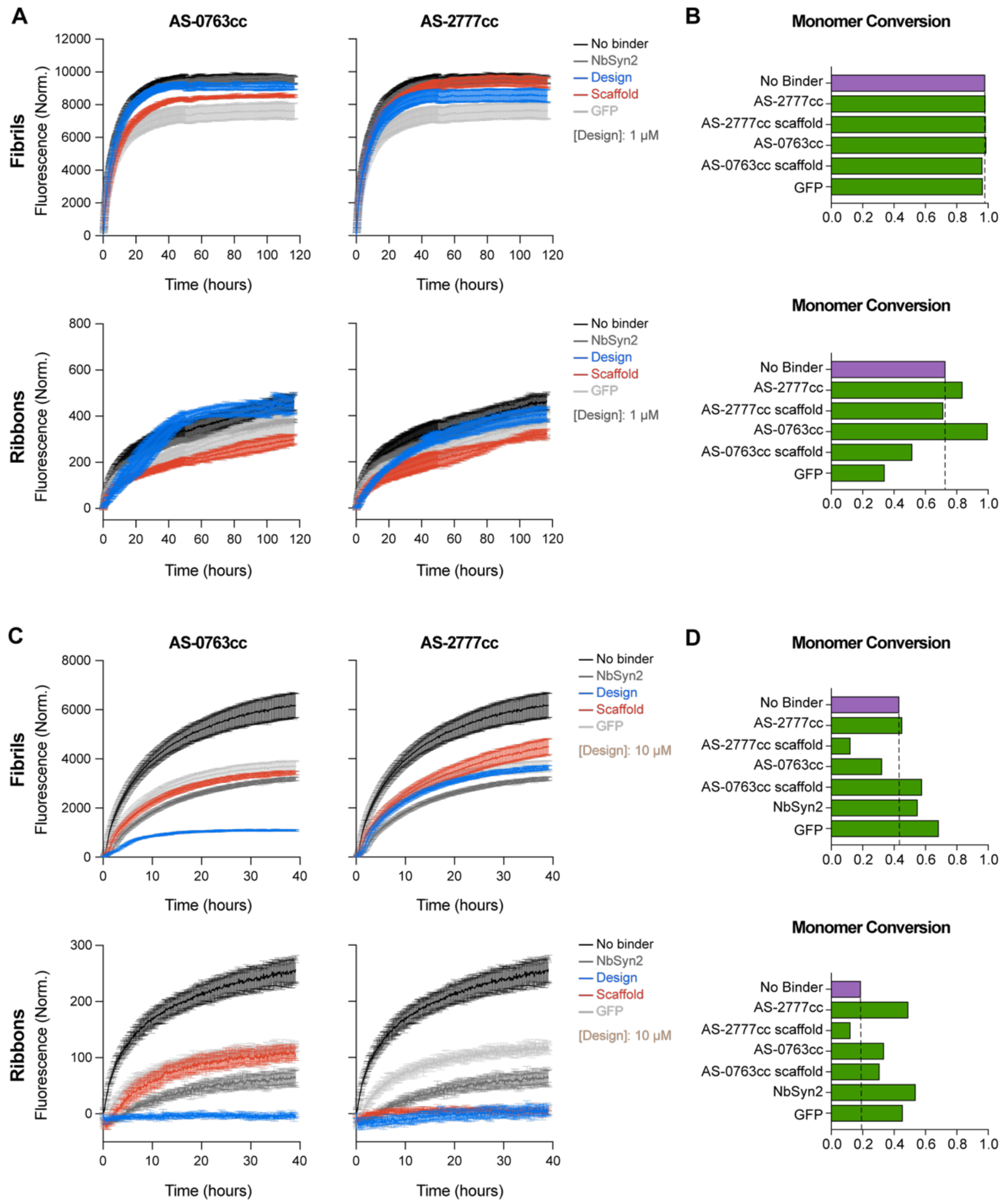


**Supplementary Figure S13 | Reproducibility of designed protein effects on aSyn seed elongation monitored by ThT binding and SDS-PAGE.**

**(A)** ThT aggregation kinetics curves showing the effect of 1 μM designs on the elongation of 0.5 μM aSyn seeds at a 2:1 binder:seeds stoichiometry (in monomer equivalents) in the presence of 20 μM aSyn monomer **(Fibrils, top ; Ribbons, bottom)** **(B)** Quantification of aSyn monomer conversion into fibrils at the endpoint of the elongation experiments in (A) using SDS-PAGE. The dashed line represents aSyn monomer conversion in the absence of binders. **(C)** ThT aggregation kinetics curves showing the effect of 10 μM designs on the elongation of 0.5 μM aSyn seeds at a 1:2 binder:aSyn monomer stoichiometry in the presence of 20 μM aSyn monomer **(Fibrils, top; Ribbons, bottom)** **(D)** Quantification of aSyn monomer conversion into fibrils at the endpoint of elongation experiments in (C) after fibrils sedimentation by centrifugation using SDS-PAGE. The dashed line represents aSyn monomer conversion in the absence of binders. These experiments are biological replicates to the ones reported in **Fig. 4**. SDS-PAGE gels used for monomer quantification are shown in **Supplementary Fig. S15**. ThT binding kinetics experiments were performed in duplicate (AS-2777cc, AS-2777cc scaffold, and GFP with Ribbons, AS-2777cc scaffold with Fibrils) or triplicate; mean values are plotted, and error bars represent standard deviation (SD); outlier measurements were excluded.

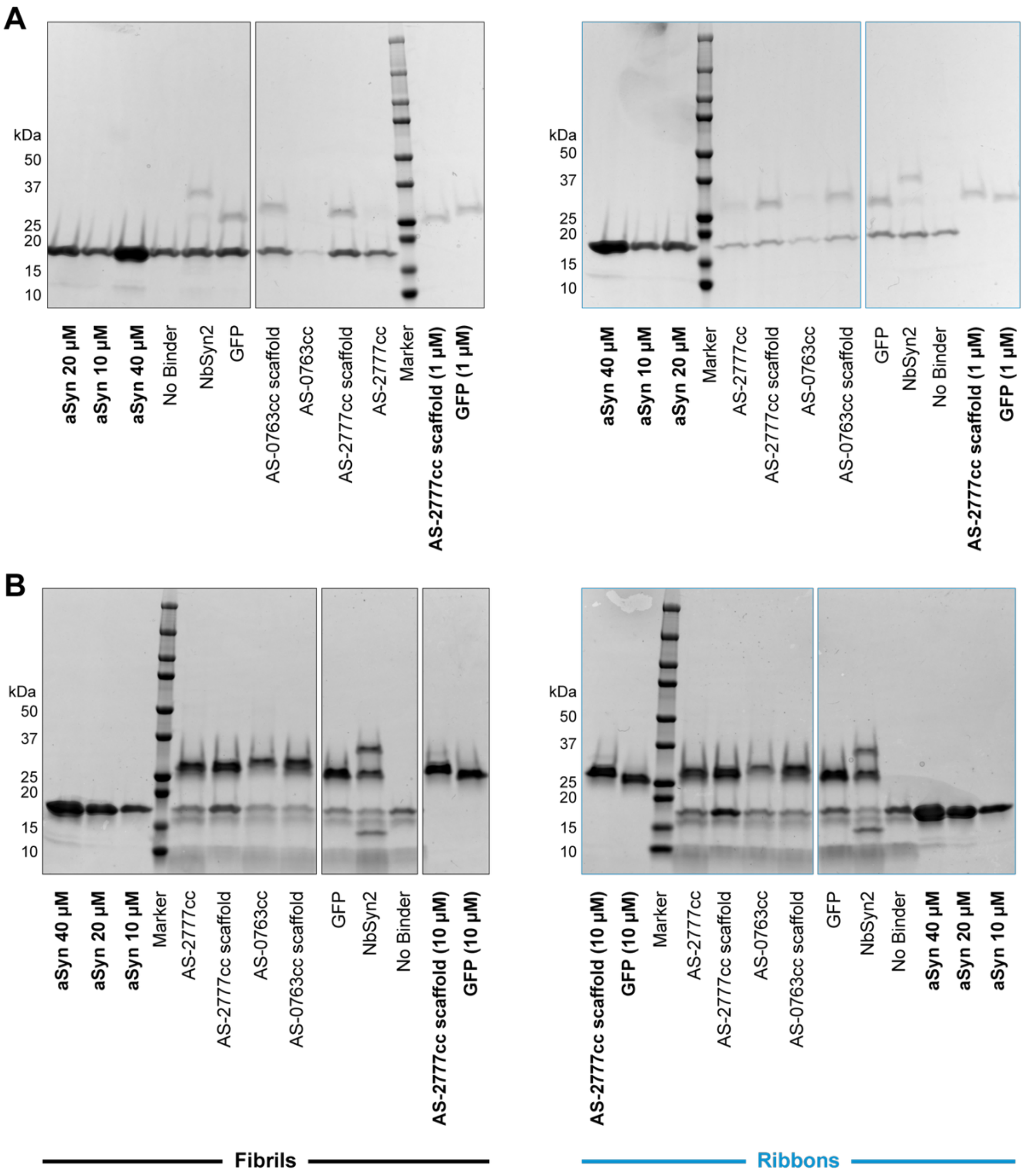


**Supplementary Figure S14 | SDS-PAGE analysis of aSyn monomer conversion into fibrils at the endpoint of aSyn seed elongation assays monitored by ThT binding reported in Fig. 4.** SDS-PAGE gels were used to quantify residual aSyn monomer at the endpoint of aSyn seeds (0.5 µM) elongation kinetics in the presence of aSyn monomer (20 µM) and the indicated designs monitored by ThT binding after fibrils sedimentation by centrifugation. **(A)** Endpoint SDS-PAGE gels for reactions with 1 µM designs and aSyn seeds prepared as **Fibrils (left; black)** or **Ribbons (right; blue)**, corresponding to ThT binding assays reported in **Fig. 4B**. **(B)** Endpoint SDS-PAGE gels for reactions with 10 µM designs and aSyn seeds prepared as **Fibrils (left; black) or Ribbons (right; blue)**, corresponding to ThT binding assays reported in **Fig. 4D**. Gels include native aSyn monomer standards at 40, 20, and 10 µM, used as controls for quantification of residual aSyn monomer concentrations. Additional controls include AS-2777cc scaffold and GFP samples at 1 µM (A) or 10 µM (B), with control samples highlighted in bold for clarity.

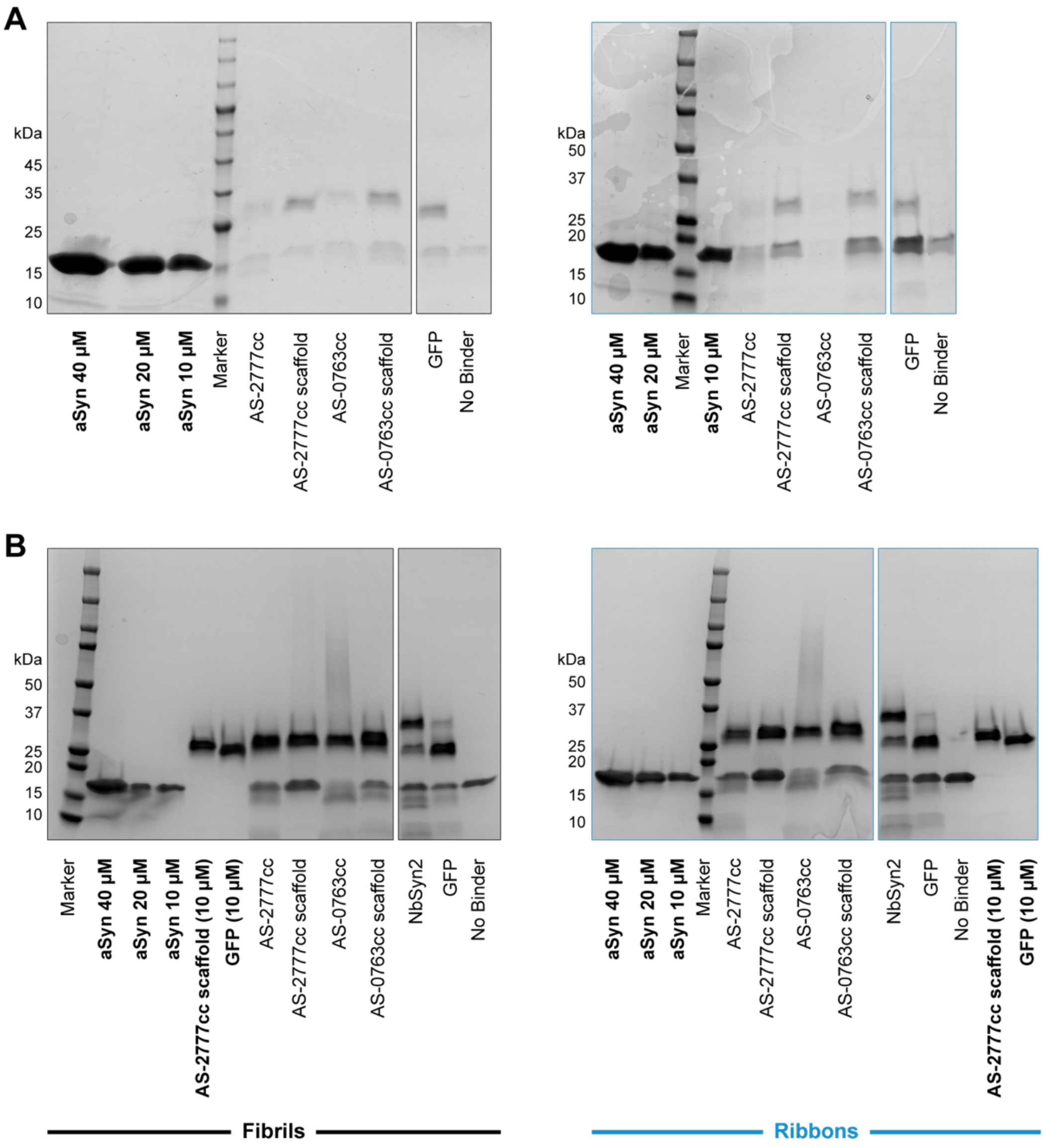


**Supplementary Figure S15 | SDS-PAGE analysis of aSyn monomer conversion into fibrils at the endpoint of aSyn seed elongation assays monitored by ThT binding reported in Supplementary Fig. S13.**

SDS-PAGE gels were used to quantify residual aSyn monomer at the endpoint of aSyn seeds (0.5 µM) elongation kinetics in the presence of aSyn monomer (20 µM) and the indicated designs monitored by ThT binding after fibrils sedimentation by centrifugation. **(A)** Endpoint SDS-PAGE gels for reactions with 1 µM designs and aSyn seeds prepared as Fibrils (left; black) or Ribbons (right; blue), corresponding to ThT binding assays reported in **Supplementary Fig. S13B**. **(B)** Endpoint SDS-PAGE gels for reactions with 10 µM designs and aSyn seeds prepared as Fibrils (left; black) or Ribbons (right; blue), corresponding to ThT binding assays reported in **Supplementary Fig. S13D**. Gels include native aSyn monomer standards at 40, 20, and 10 µM, used as controls for quantification of residual aSyn monomer concentrations. Additional controls include AS-2777cc scaffold and GFP samples at 1 µM (A) or 10 µM (B), with control samples highlighted in bold for clarity.

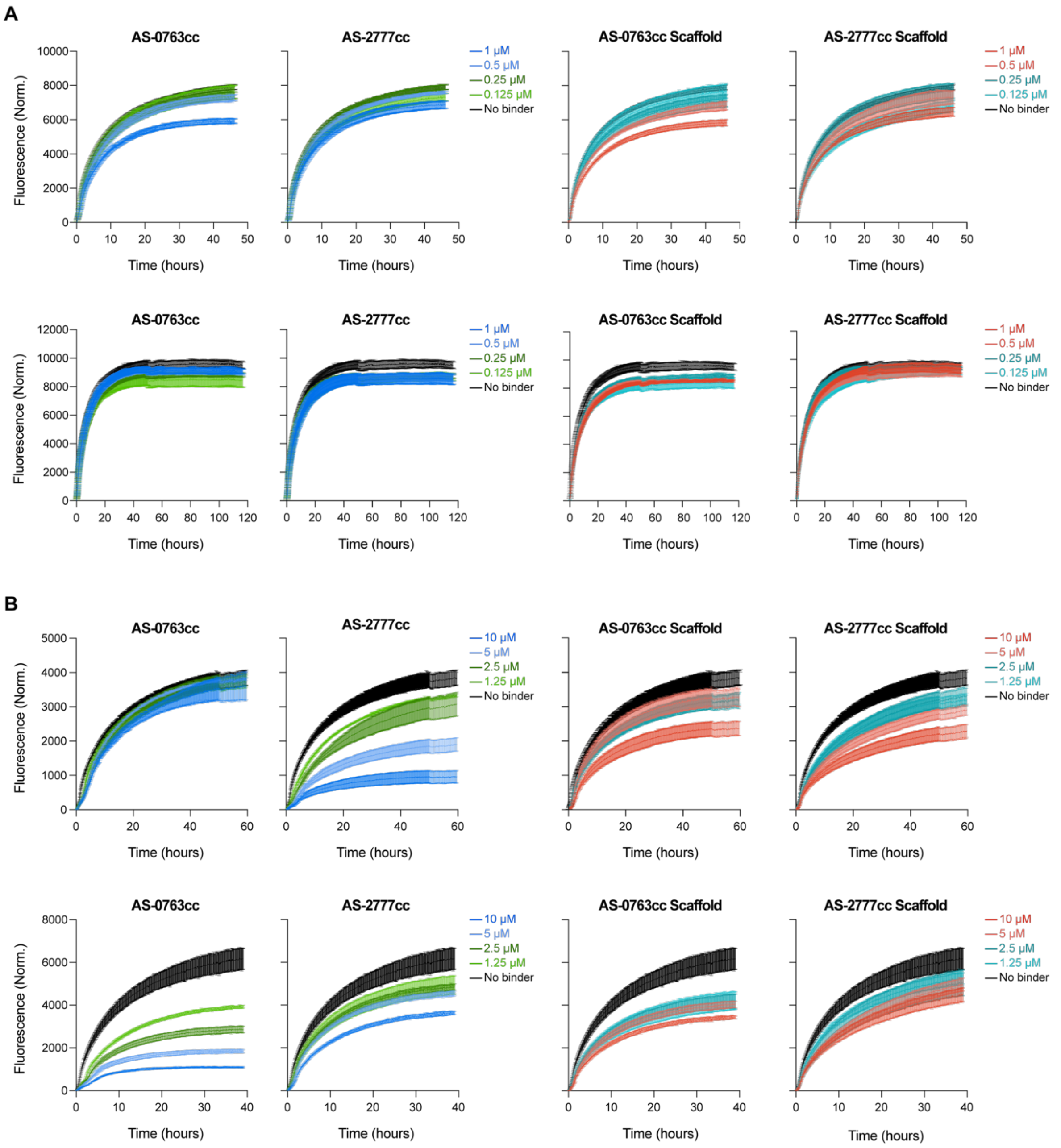


**Supplementary Figure S16 | Effect of varying designed protein concentrations on the elongation of aSyn "Fibrils" seeds assessed by ThT fluorescence.**

ThT fluorescence kinetics curves were used to monitor the elongation of 0.5 μM aSyn "**Fibrils**" seeds in the presence of 20 μM aSyn monomer and varying concentrations of binders from 0.125 μM to 1 μM (low-range; **A**) and 1.25 μM to 10 μM (higher-range; **B**). Results are from four independent experiments, each performed in triplicate, with mean values plotted and error bars representing standard deviation (SD); outlier measurements were excluded. In panel (B): data for AS-0763cc (10, 5 μM) and AS-0763cc scaffold (5 μM) in top section and data for AS-2777cc (5 μM) and AS-2777cc scaffold (10 μM) in bottom section represent 2 replicates.

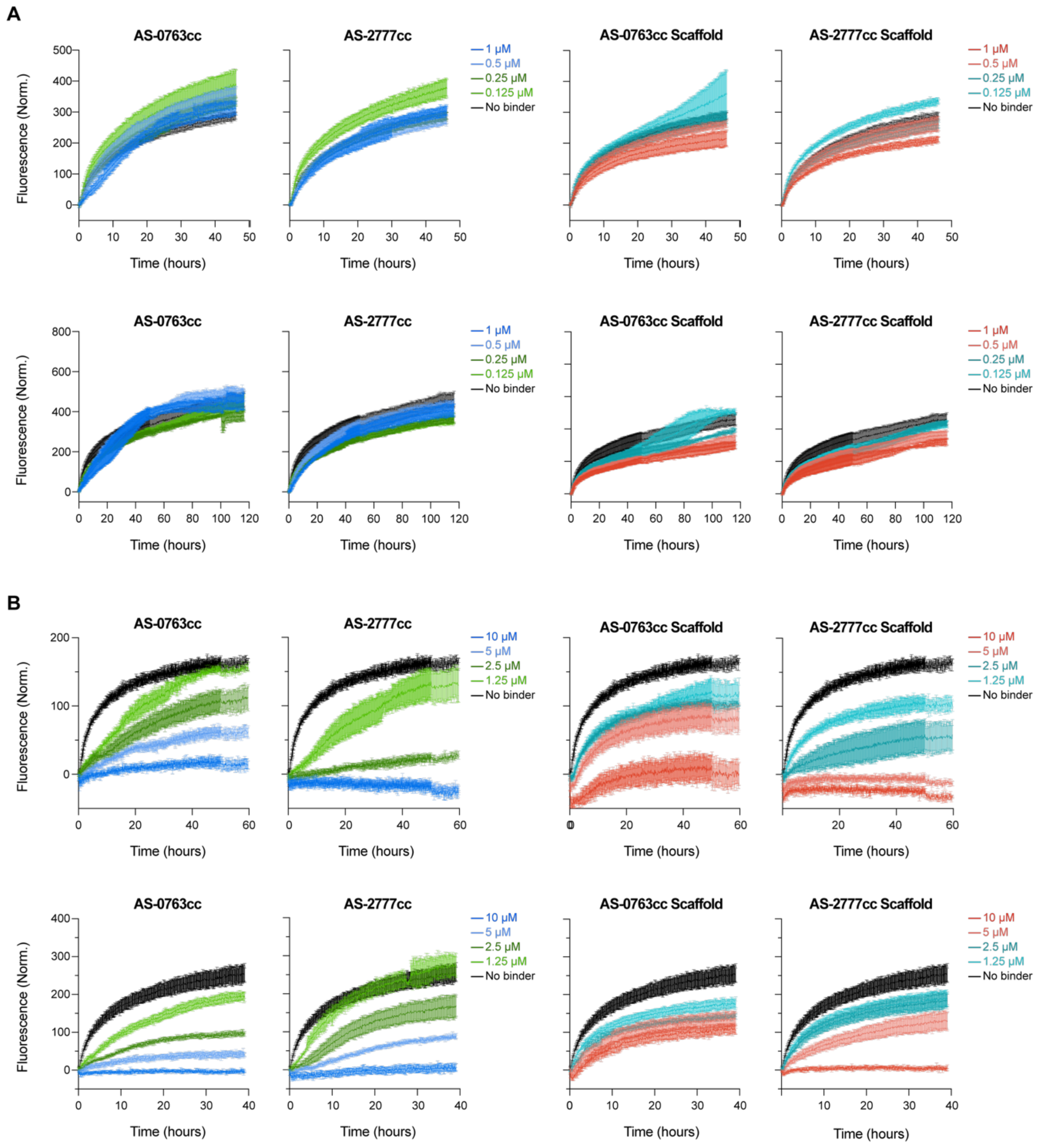


**Supplementary Figure S17 | Effect of varying designed protein concentrations on the elongation of aSyn "Ribbons" seeds assessed by ThT fluorescence.**

ThT fluorescence kinetics curves were used to monitor the elongation of 0.5 μM aSyn "**Ribbons**" seeds in the presence of 20 μM aSyn monomer and varying concentrations of binders from 0.125 μM to 1 μM (low-range; **A**) and 1.25 μM to 10 μM (higher-range; **B**). Results are from four independent experiments, each performed in triplicate, with mean values plotted and error bars representing standard deviation (SD); outlier measurements were excluded. In panel (A): data for AS-0763cc (0.5, 0.25, 0.125 μM), AS-2777cc (1, 0.5 μM), AS-0763cc scaffold (0.25, 0.125 μM), and AS-2777cc scaffold (1 μM) in bottom section; and in panel (B): data for AS-2777cc (1.25 μM) in the bottom section represent 2 replicates.

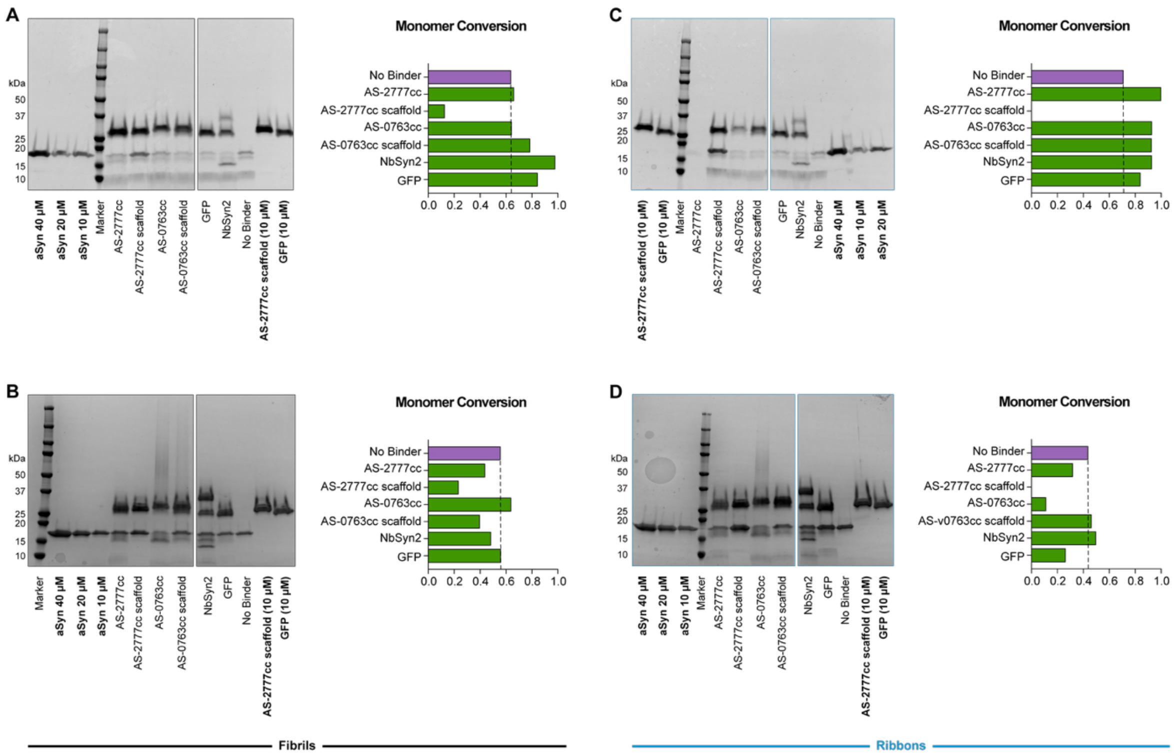


**Supplementary Figure S18 | SDS-PAGE analysis of aSyn monomer conversion into fibrils at the endpoint of aSyn seed elongation assays in the absence of ThT.**

SDS-PAGE gels used to quantify residual aSyn monomer at the endpoint of aSyn seeds (0.5 µM) elongation kinetics in the presence of aSyn monomer (20 µM) and the indicated designs (10 µM) and in the absence of ThT after fibrils sedimentation by centrifugation. Endpoint gels and quantifications show aSyn monomer conversion for the elongation of **(A, B)** aSyn seeds (**Fibrils: A, B**, black**; Ribbons: C, D**, blue) after 60 hours **(A, C)** and 40 hours **(B, D)**. Related experiments, performed in the presence of ThT are shown in **Fig. 4C for (A, C)**, and **Supplementary Fig. S13C for (B, D)**. Assays were conducted in low-bind Eppendorf tubes. Gels include native aSyn monomer standards at 40, 20, and 10 µM, used as controls for quantification of residual aSyn monomer concentrations. Additional controls include AS-2777cc scaffold and GFP samples at 10 µM, with control samples highlighted in bold for clarity. The dashed line represents aSyn monomer conversion in the absence of binders.

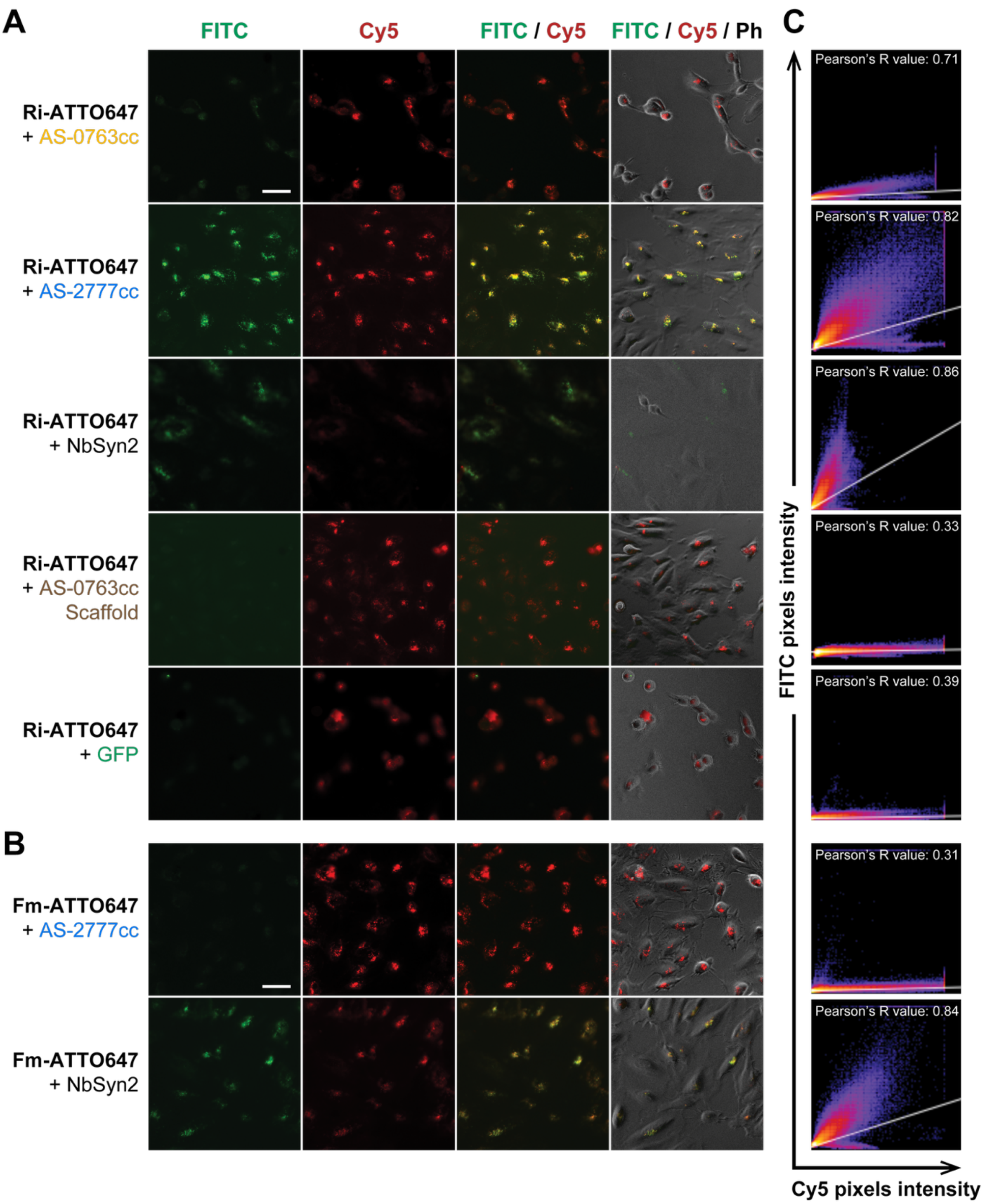


**Supplementary Figure S19 | Internalization of aSyn polymorphs in presence of binders by HeLa cells.**

**(A, B)** Representative live-cell fluorescence micrographs acquired after HeLa cells were exposed to Ri-ATTO647 (**A**) or Fm-ATTO647 (**B**) together with GFP-fused binder or controls (FITC: green; Cy5: red; Ph: Phase contrast). Cy5 signals correspond to aSyn polymorphs bound to or internalized by cells, whereas FITC signals reflect exclusively internalized GFP-fused binders. Additional control conditions are shown in **Supplementary Fig. S20C. (C)** Scatter plots showing pixel-based colocalization between the Cy5 (x-axis) and FITC (y-axis) channels for the micrographs shown in **(A)** and **(B)**. Object-based colocalization analysis is provided in **Supplementary Fig. S21B.** Scale bar = 50 µm.

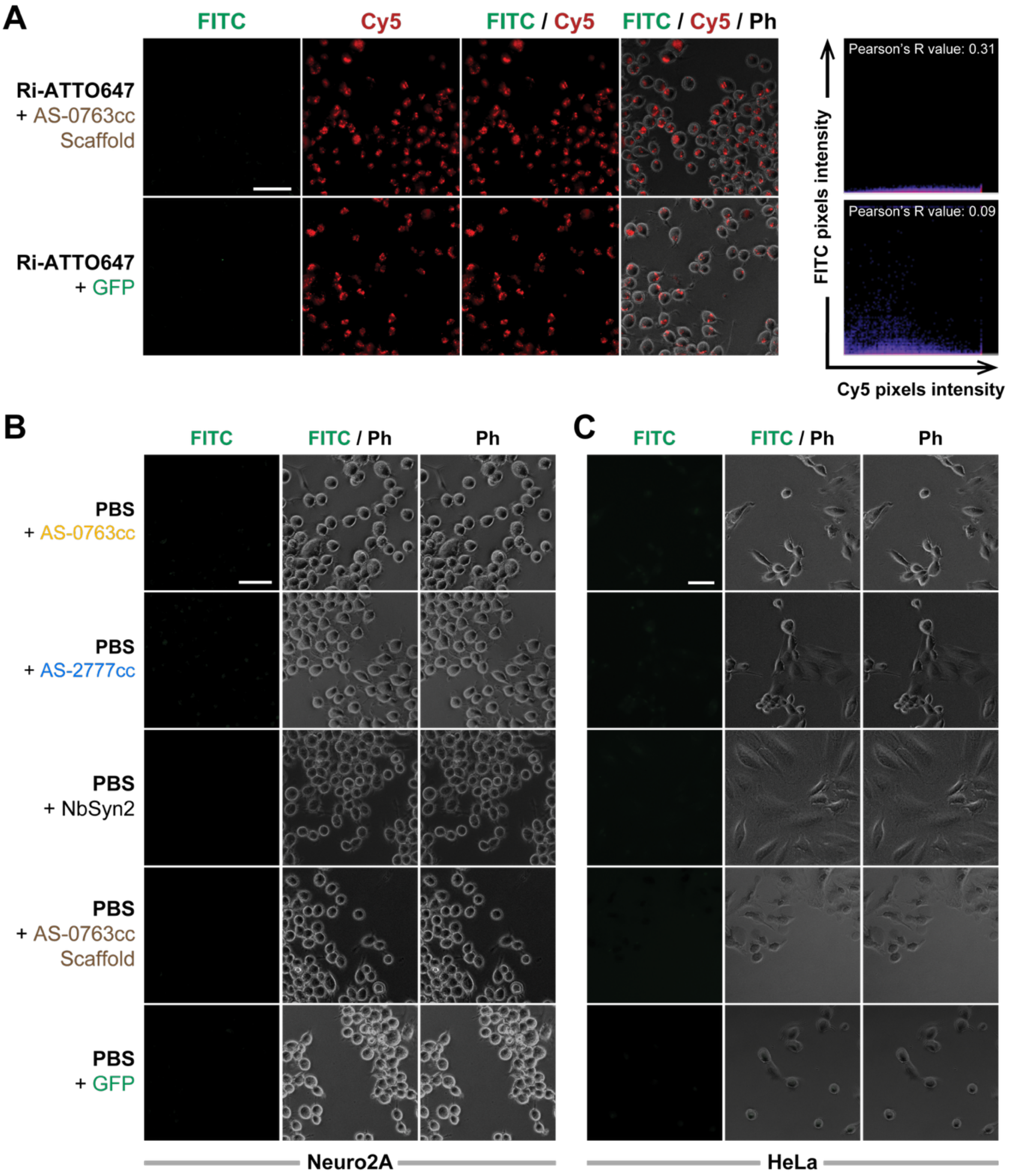


**Supplementary Figure S20 | Internalization of binders by Neuro2A and HeLa cells in presence or absence of ATTO647-labeled aSyn polymorphs, additional conditions.** Representative photomicrographs acquired by live fluorescence microscopy 2 hours after exposure of Neuro2A **(A, B)** and HeLa **(C)** cells to GFP-fused binders (red: Cy5; green: FITC; Phase contrast: Ph). **(A)** Neither GFP alone nor the AS-0763cc scaffold was internalized by Neuro2A cells in presence of Ribbons (Ri). Scatter plots to the right of the images illustrate the absence of pixel-based colocalization between Cy5 (x-axis) and FITC (y-axis) channels under these conditions. **(B, C)** No internalized FITC signal was detected in Neuro2A **(B)** or HeLa cells **(C)** exposed to binders in absence of aSyn polymorphs (Fibrils, Fm or Ribbons, Ri). Scale bar = 50 µm.

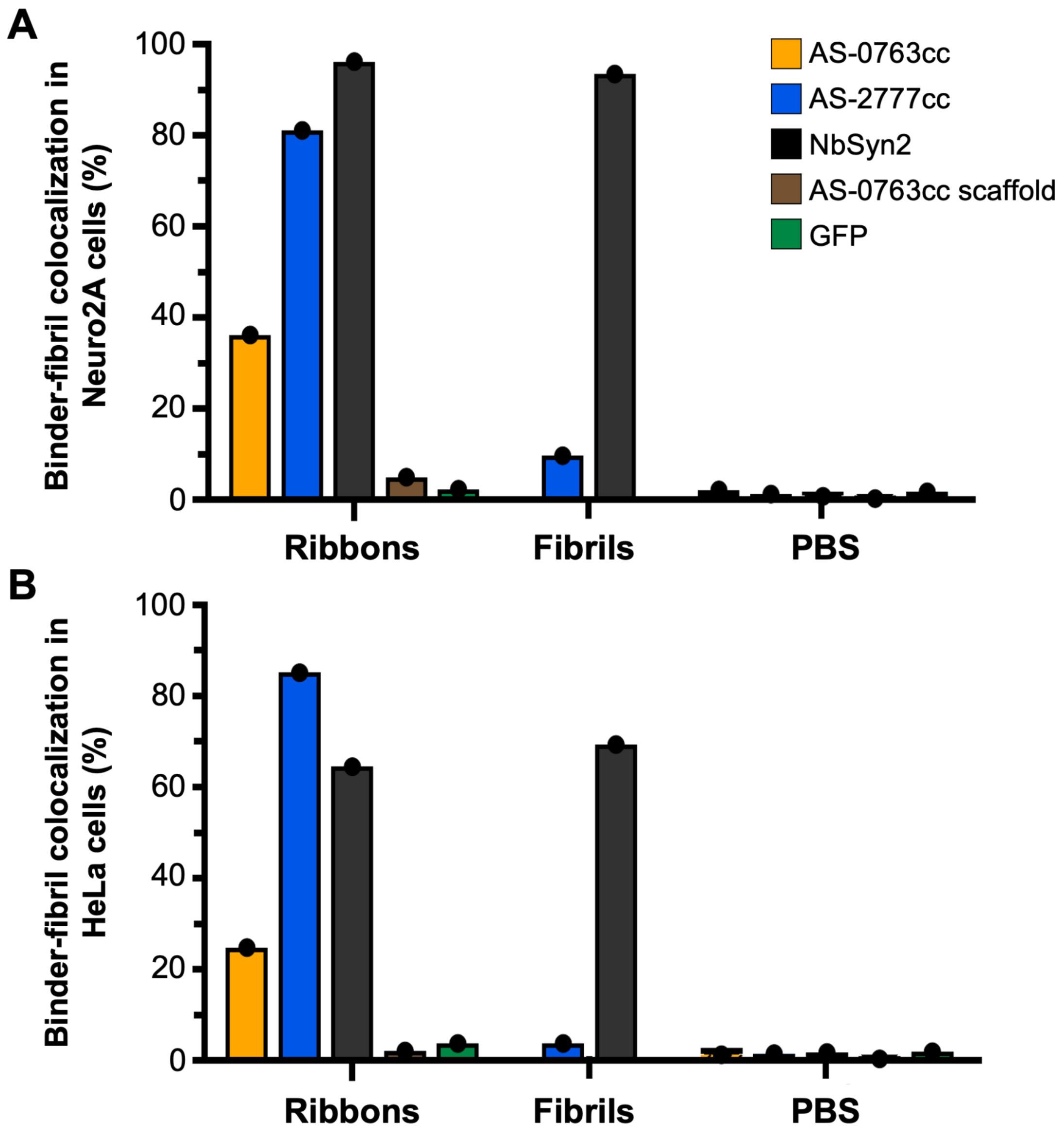


**Supplementary Figure S21 | Colocalization of binders with aSyn polymorphs after internalization by Neuro2A and HeLa cells.**
Object-based colocalization analysis was performed on photomicrographs acquired by live fluorescence microscopy two hours after exposure of cells to GFP-fused binders alone or to binders preincubated with ATTO647-labeled aSyn polymorphs, as shown in **Fig. 5** and **Supplementary Fig. S19 and S20**. Colocalization was quantified as the percentage of Cy5-positive objects colocalizing with FITC-positive objects in Neuro2A **(A)** and HeLa **(B)** cells.

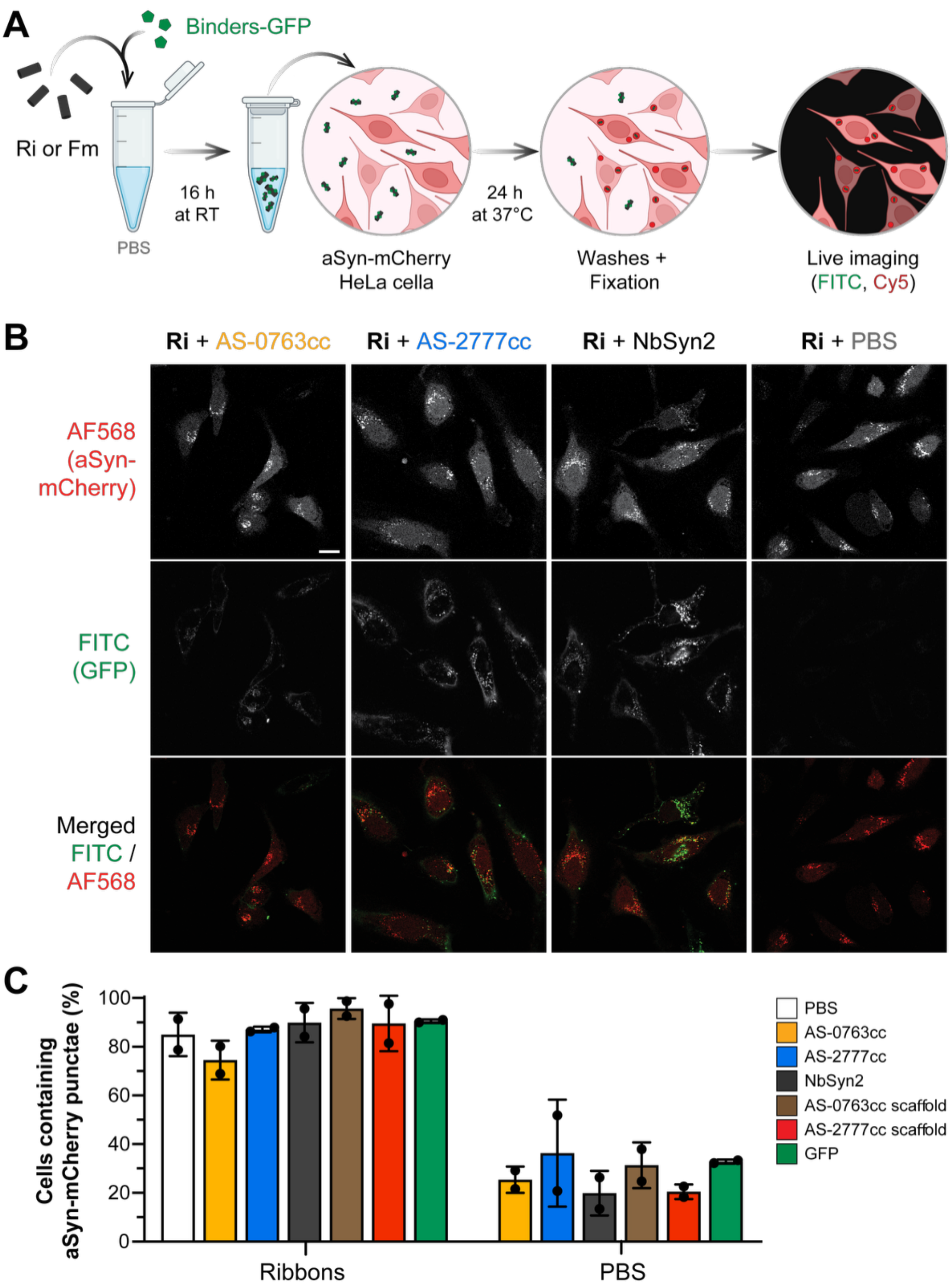


**Supplementary Figure S22 | Seeded aggregation of aSyn in HeLa cells in the presence and absence of binders.**

**(A)** Experimental outline for assessing the effect of GFP-fused binders on the seeding of unlabeled aSyn polymorphs. **(B)** Representative confocal photomicrographs of HeLa cells expressing aSyn–mCherry, fixed 24 hours after exposure to fragmented Ribbons (Ri; average length: 45 nm) in presence or absence of binders. Exposure to Ri induces the formation of bright mCherry-positive punctae detected in the AF568 channel. GFP-fused binders are detected in the FITC channel. Additional control conditions are shown in **Supplementary Fig. S23**. Scale bar = 20 µm. **(C)** Quantification of the proportion of HeLa cells containing mCherry-positive punctae 24 hours after exposure to Ri in presence or absence of binders. Experiments were performed in two independent replicates.

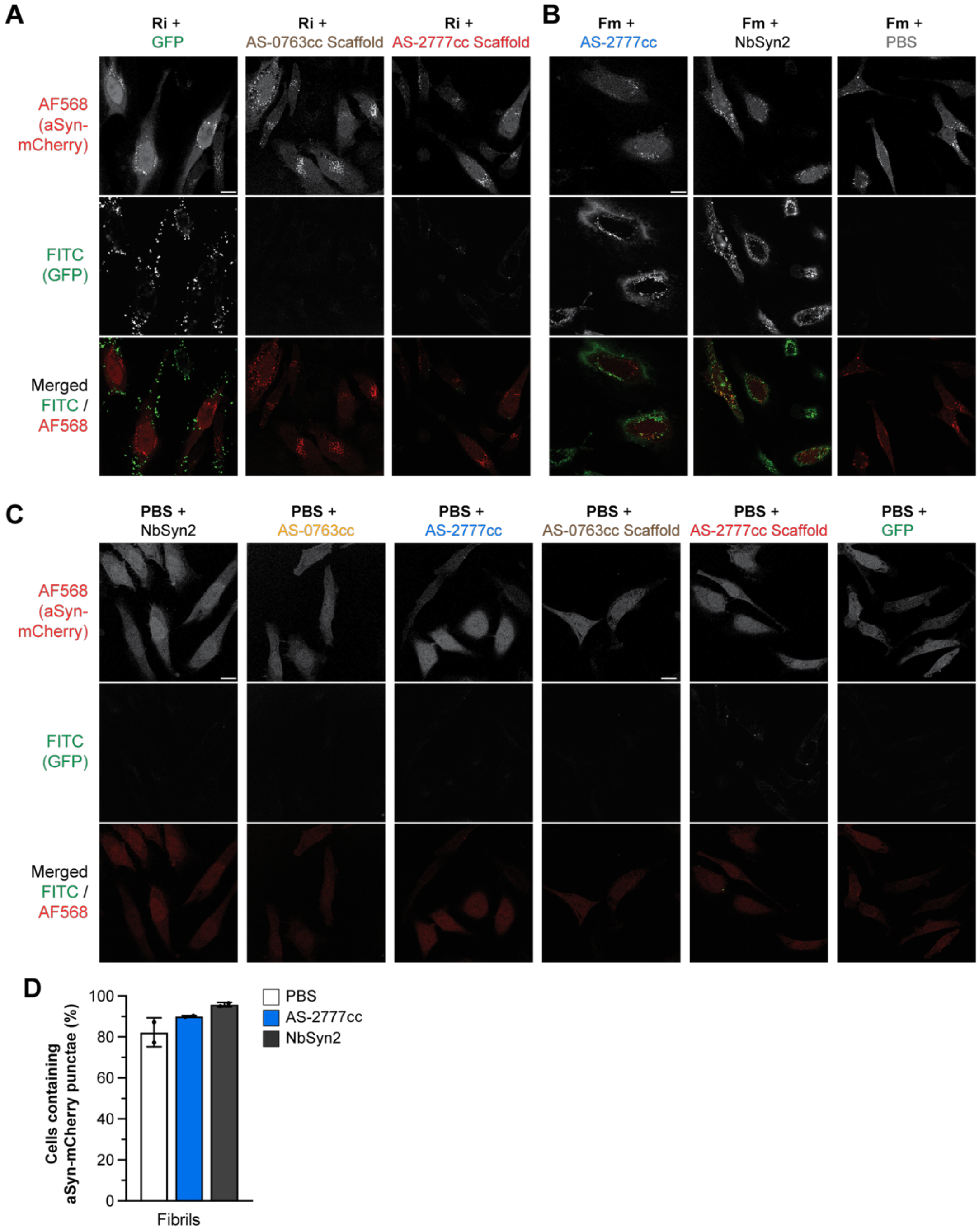


**Supplementary Figure S23 | Seeded aggregation of aSyn in HeLa cells in presence or absence of binders, additional conditions.**
**(A-C)**. Representative confocal photomicrographs of HeLa cells expressing aSyn-mCherry, fixed 24 hours after exposure to unlabeled aSyn polymorphs, either Ribbons (Ri; **A**), Fibrils (Fm; **B**) or PBS **(C)**, in presence or absence of GFP-fused binders. Exposure to either aSyn polymorph induces the formation of bright mCherry-positive punctae (AF568 channel). GFP-fused binders are detected in the FITC channel. Scale bar = 20 µm. **(D)** Quantification of the proportion of HeLa cells containing mCherry-positive punctae 24 hours after exposure to Fm in presence or absence of binders. Experiments were performed in two independent replicates.

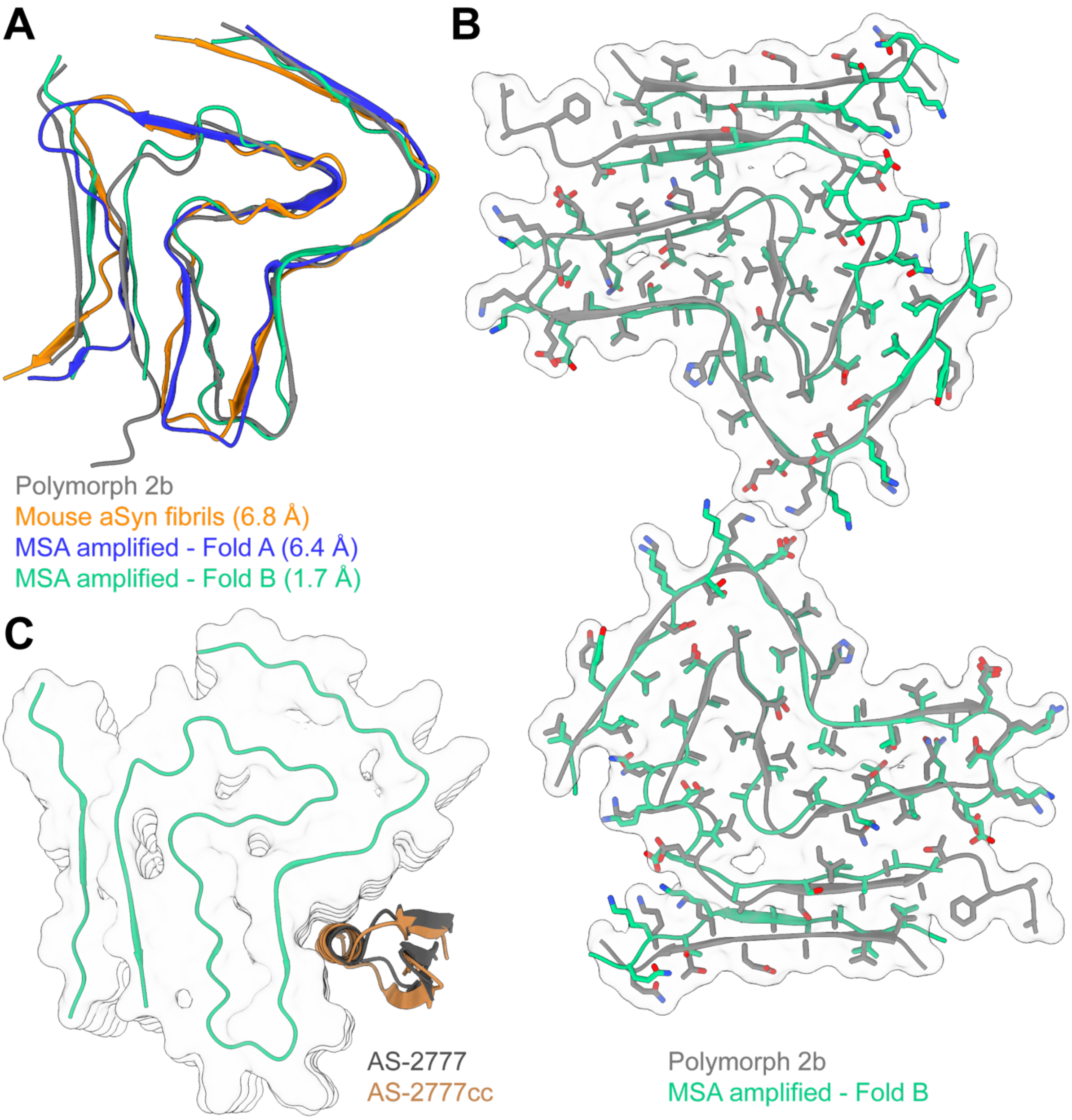


**Supplementary Figure S24 | Clinical relevance of *in-vitro* aSyn Polymorph 2b Fibrils.**
**(A)** Structural overlays comparing one protofilament layer of fibrils prepared *in vitro* from recombinant human WT aSyn either *de novo* aggregated Polymorph 2b (PDB: 6SST, grey) or seeded by MSA patient material (Fold A, PDB: 7NCG, blue; Fold B, PDB: 7NCI, green), and mouse aSyn (PDB: 8UIE, orange). The $C_\alpha$ RMSD was calculated using the UCSF ChimeraX MatchMaker tool (V 1.8) following model alignment. **(B)** Structural overlay of one fibril layer of aSyn Polymorph 2b (grey) and MSA-amplified Fold B fibrils (green), demonstrating strong agreement in backbone and sidechain positioning ($C_\alpha$ RMSD = 1.7 Å). **(C)** MaSF-search docked models of AS-2777 (grey) and AS-2777cc (brown) to MSA-amplified Fold B fibrils with descriptor distance scores (AS-2777: 27.3, AS-2777cc: 28.6) passing our defined thresholds (**Supplementary Fig. S25**).

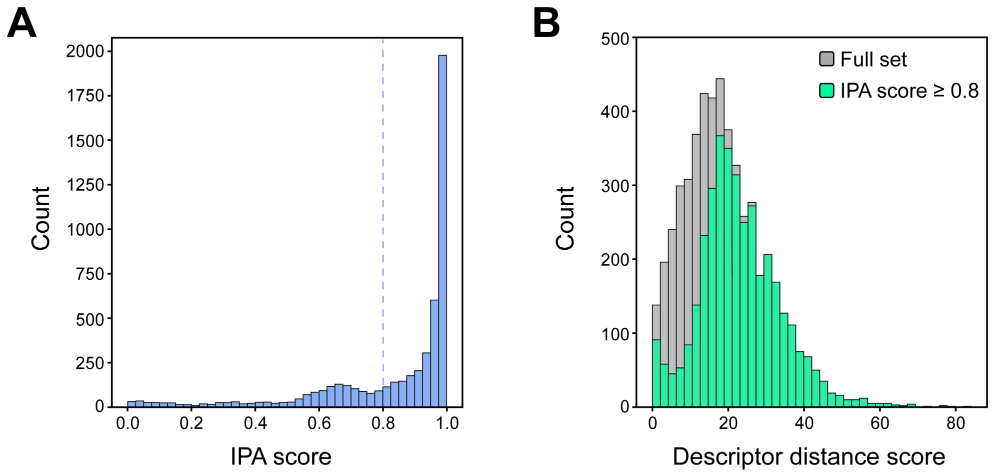


**Supplementary Figure S25 | Benchmarking MaSIF-Search descriptor distance score for patch complementarity.**

MaSIF-search descriptor distance scores, representing patch-to-patch complementarity based on surface fingerprints, were benchmarked using 5,188 patch pairs from 872 native complexes in the published MaSIF-search test set. Aligned patches were evaluated using the interface post-alignment (IPA) neural network, which assigns a score between 0 and 1 to indicate the likelihood that an alignment corresponds to a real interaction.

**(A)** MaSIF-search IPA scores for docked patches showed that 70% of the complex patches (3,665 / 5,188) exceeded the recommended threshold (0.8, dashed line) for a potential real interaction. **(B)** Descriptor distance scores calculated for all tested patches (Full set, median: 17.7, grey) and for patch matches with IPA score ≥ 0.8 (median: 21.8, green). Median descriptor distance scores were used as reference points for assessing the docking of *de novo* designs to fibril structures, where higher scores indicate better docked matches.

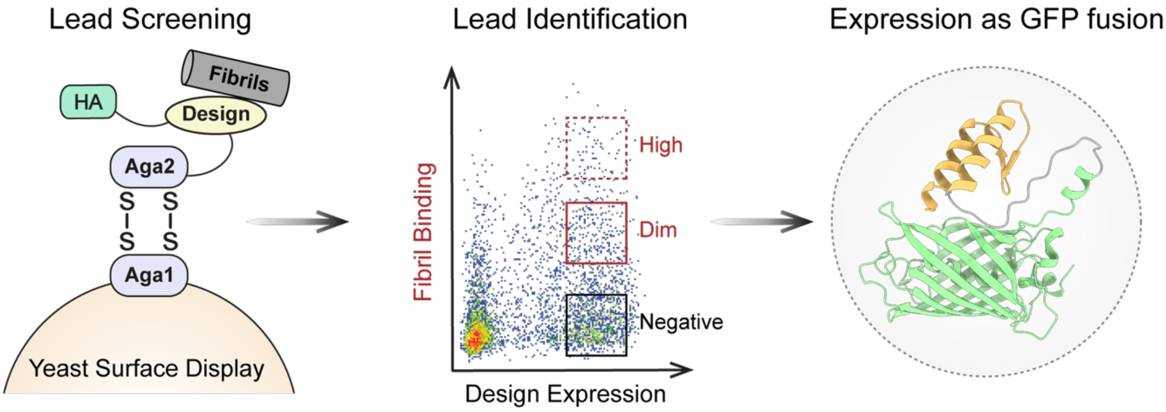


**Supplementary Figure S26 | Workflow for lead identification using yeast surface display and next generation sequencing.**
Approximately 6,000 *de novo* designs were screened against two aSyn polymorphs (Fm and Ri)[63] using yeast surface display. Design expression was detected using a FITC conjugated anti-HA antibody recognizing an HA tag at the C-terminus of each construct, with FITC$^{+}$ cells indicating design expression. Fibril Binding was assessed through sequential antibody labeling: anti-aSyn primary antibody followed by an Alexa Fluor™ 647-conjugated secondary antibody, where Alexa Fluor™ 647$^{+}$ cells indicated fibril binding. The double-positive (FITC$^{+}$/Alexa Fluor™ 647$^{+}$) population represented yeast expressing designs bound to fibrils. Based on Alexa Fluor™ 647 fluorescence intensity, yeast cells were sorted by fluorescence-activated cell sorting (FACS) into three populations: non-binders (negative), potential binders (dim), and strong binders (high). Two sequential rounds of sorting were performed as described under *Flow cytometry analysis and sorting of designs library* in the Methods section.

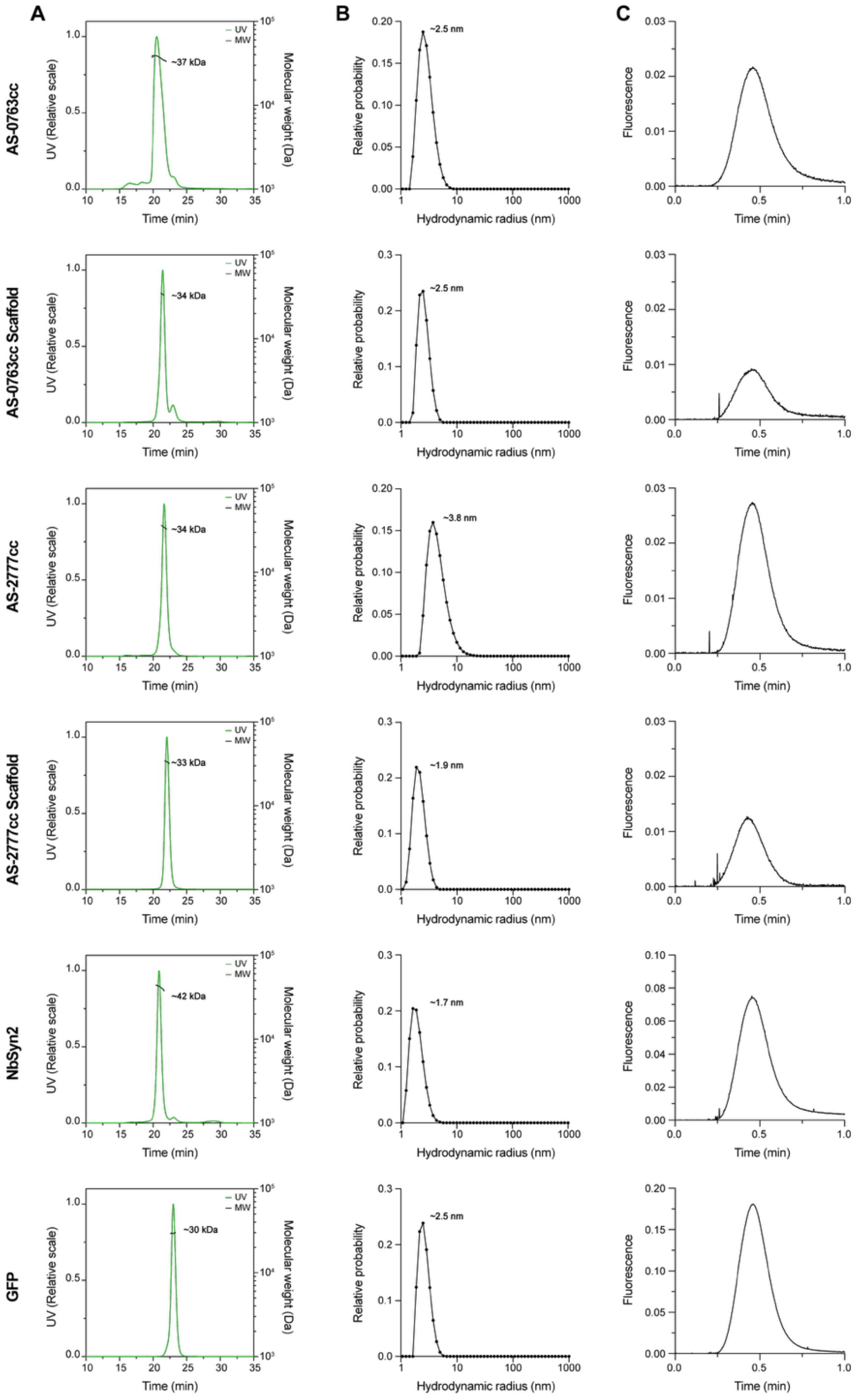


**Supplementary Figure S27 | Biophysical characterization of GFP-fused designs.**
**(A)** Molecular weight and oligomeric state of GFP-fused designs determined by SEC-MALS. **(B)** DLS analysis of binders at 10 µM and room temperature. Plots show normalized mass-weighted distributions of sample populations derived from the size distribution fit of the autocorrelation function (ACF). For each sample, ten ACFs were recorded from two independent capillaries. **(C)** FIDA 488/525 nm Taylorgrams of the GFP-fused binders at 100 nM. All designs predominantly (>90%) exhibited a diffusive peak, with negligible signs of aggregation (spikes on the left-side of the elution profile). Accurate determination of the hydrodynamic radius from the Taylorgram was hindered by sample sticking to the capillary, apparent as peak tailing at later times and varying amplitudes.

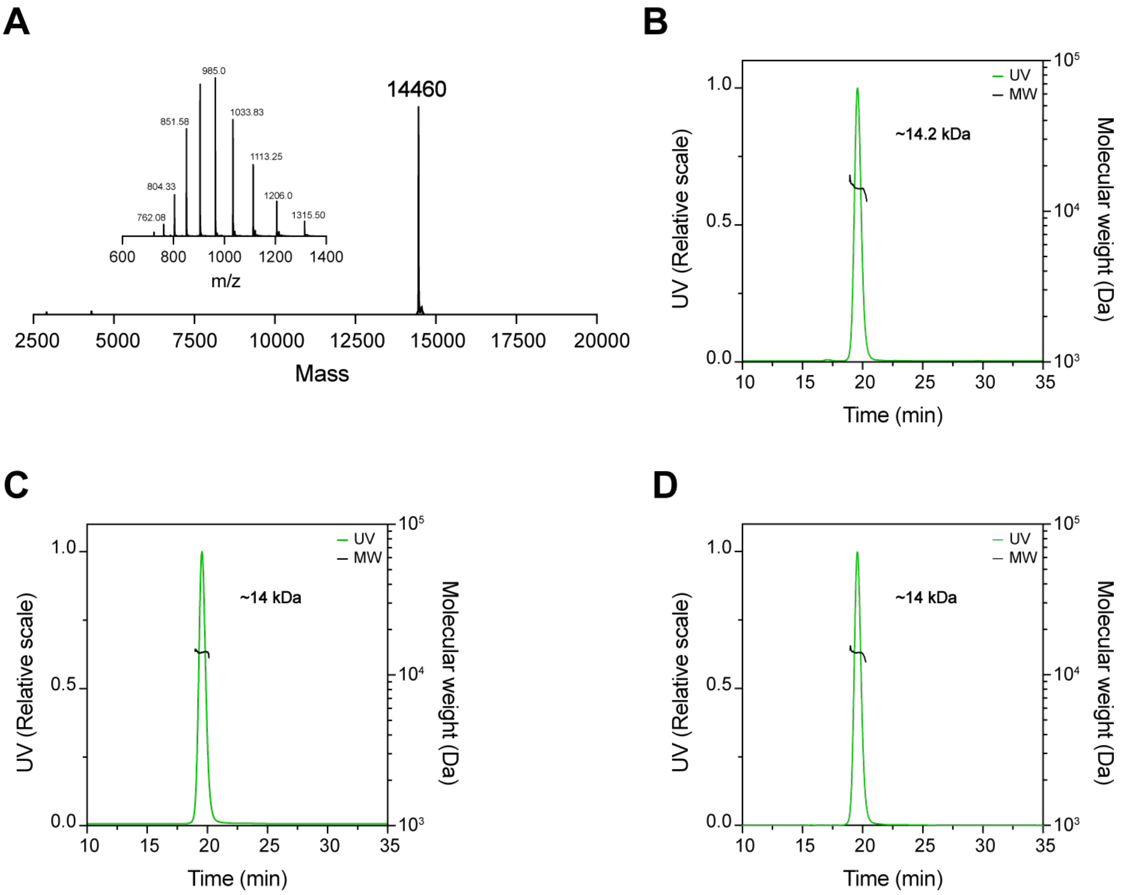


**Supplementary Figure S28 | Characterization of aSyn monomer by ESI-MS and SEC-MALS.**
**(A)** ESI-MS spectrum of HPLC-purified human WT aSyn, showing the observed molecular weight matches the theoretical value (14,460 Da). **(B-D)** Molecular weight and oligomeric state of aSyn monomers determined by SEC-MALS for: **(B)** HPLC-purified aSyn monomers, **(C)** SEC-purified aSyn monomers, and **(D)** aSyn monomer after concentration and buffer exchange using 3-kDa filters. SEC-MALS results confirm that aSyn monomers used in this study were monomeric across all preparation methods.

**Supplementary Table S1 | Summary of Rosetta design strategies for grafted designs**

This table summarizes the design strategies used to generate 5,545 designs targeting aSyn Polymorphs 1a and 2b. Except for Polymorph 1a FO, SO, $SO_{pm}$ strategies, design backbone minimization was prevented, and multiple degrees of freedom were enabled for side chain chi angles. For Polymorph 1a, filtered designs from FO, SO, $SO_{pm}$, and $SO_{alt}$ (829 designs) were stabilized via disulfide bond engineering and further optimized in Rosetta (Disulf), followed by a final optimization step to diversify the designed interfaces ($Disulf_{opt}$). For Polymorph 2b, two rounds of interface-focused refinement were performed: in round 1 (rd1), no constraints were imposed on interface residue composition, whereas in round 2 (rd2), methionine and glycine were prohibited at the interface, and phenylalanine, tryptophan, and tyrosine were each limited to a maximum of 10% to reduce hydrophobicity and potential aggregation. rd2 optimization was performed on a subset of rd1 designs (S6-S9).

FO: Full Optimization | SO: Scaffold Optimization | YSD: Yeast Surface display

| Target | Strategy name | Designed fraction | Design protocol | Chi angles repack | Filtered designs | YSD tested designs |
|---|---|---|---|---|---|---|
| Polymorph 1a | FO | Full binder | FastDesign Layer Design | Design | 248 | - |
| | SO | Scaffold interface | FastDesign Layer Design | Design | 179 | - |
| | $SO_{pm}$ | Scaffold interface | PackRotamers Layer Design | Design | 228 | - |
| | $SO_{alt}$ | Scaffold interface | PackRotamers Layer Design | Design | 174 | - |
| | Disulf | Scaffold interface | PackRotamers Layer Design | Design & Target | 132 | 108 |
| | $Disulf_{opt}$ | Full binder | FastDesign Layer Design | Design & Target | 132 | 114 |
| Polymorph 2b | rd1 – S1 | Scaffold interface | FastDesign Layer Design | Design | 2,152 | 560 |
| | rd1 – S2 | Scaffold interface | PackRotamers Layer Design | Design | 2,094 | 453 |
| | rd1 – S3 | Full binder | FastDesign Layer Design | Design & Target | 387 | 63 |
| | rd1 – S4 | Scaffold interface | FastDesign Layer Design | Design & Target | 1,548 | 218 |
| | rd1 – S5 | Full binder | PackRotamers Layer Design | Design | 614 | 105 |
| | rd1 – S6 | Scaffold interface | FastDesign Layer Design | Design & Target | 1,141 | 417 |
| | rd1 – S7 | Full interface | FastDesign Layer Design | Design & Target | 256 | 91 |
| | rd1 – S8 | Scaffold interface | FastDesign | Design & Target | 1,388 | 583 |
| | rd1 – S9 | Full interface | FastDesign | Design & Target | 700 | 365 |
| | rd2 | Full interface | FastDesign | Design & Target | 2,677 | 2,468 |

#### Supplementary Table S2 | Summary of all sorted designs.

A selection of ~6,000 designs were screened using YSD against two aSyn fibril preparations (Fm and Ri). Binding yeast cells were isolated using fluorescence-activated cell sorting (FACS) in two sequential rounds, using 1 µM fibrils for the first and 0.5 µM in the second. Cells were sorted into non-binding and two binding populations (Dim and High) based on fluorescence shift upon target binding. For each design, all targeting polymorph 2b, the design strategy is reported, with rd2 designs indicating their originating rd1 strategy. The sorted population, target fibril, and sorting conditions (sort 1 / sort 2 concentrations) are also provided. Some designs were identified via single-colony PCR on isolated cells post-sorting.

Binding and non-binding counts, along with the enrichment value (log[counts binding / counts non-binding]), indicate fold increase in binding. Designs with ≥ 10-fold enrichment (enrichment ≥ 1) were selected for further characterization. YSD was further used to assess the binding profiles of selected designs against different aSyn fibril preparations (Fm, Ri, and F91; 250 nM), alongside a negative control (Ab control). Binding specificity was evaluated using interface knockout mutants and scaffold-only controls. Experimentally tested designs are highlighted in bold.

| # | Design | Design strategy | Sort population | Counts Non-Bind. | Counts Binding | Enrichment *(log scale)* | YSD binding | YSD Specificity |
|---|---|---|---|---|---|---|---|---|
| **1** | **AS-0763** | **Pol 2b<br>rd1 – S1** | **Ribbons – Dim<br>1 µM / 0.5 µM** | **96** | **1495** | **1.19** | **Ribbons: ++<br>Fibrils: +<br>Fibrils-91: +<br>Ab control: -** | **- KO mutant<br>-Scaffold-only** |
| 2 | AS-1110 | Pol 2b<br>rd1 – S2 | Single colony PCR | - | - | - | Ribbons: +++<br>Fibrils: +++<br>Fibrils-91: +++<br>Ab control: +++ | - |
| **3** | **AS-1210** | **Pol 2b<br>rd1 – S2** | **Single colony PCR** | **-** | **-** | **-** | **Ribbons: +<br>Fibrils: +<br>Fibrils-91: ++<br>Ab control: -** | **-** |
| 4 | AS-1563 | Pol 2b<br>rd1 – S4 | Fibrils – High<br>1 µM / 0.5 µM | 38 | 1061 | 1.45 | Ribbons: ++<br>Fibrils: ++<br>Fibrils-91: +++<br>Ab control: + | - |
| 5 | AS-1930 | Pol 2b<br>rd2 (rd1 – S6) | Single colony PCR | - | - | - | Ribbons: +++<br>Fibrils: +++<br>Fibrils-91: +++<br>Ab control: ++ | -Scaffold-only |
| 6 | AS-2028 | Pol 2b<br>rd2 (rd1 – S6) | Fibrils – High<br>1 µM / 0.5 µM | 1191 | 123701 | 2.02 | Ribbons: +++<br>Fibrils: ++<br>Fibrils-91: ++<br>Ab control: - | - KO mutant |
| | | | Fibrils – Dim<br>1 µM / 0.5 µM | 1191 | 66881 | 1.75 | | |
| | | | Ribbons – High<br>1 µM / 0.5 µM | 32 | 89300 | 3.45 | | |
| | | | Ribbons – Dim<br>1 µM / 0.5 µM | 32 | 718 | 1.35 | | |
| 7 | AS-2762 | Pol 2b<br>rd2 (rd1 – S6) | Single colony PCR | - | - | - | Ribbons: +++<br>Fibrils: +++<br>Fibrils-91: +++<br>Ab control: ++ | - |
| **8** | **AS-2777** | **Pol 2b<br>rd2 (rd1 – S6)** | **Ribbons – Dim<br>1 µM / 0.5 µM** | **1729** | **38824** | **1.35** | **Ribbons: +++<br>Fibrils: -<br>Fibrils-91: -<br>Ab control: -** | **- KO mutant<br>-Scaffold-only** |
| 9 | AS-2926 | Pol 2b<br>rd2 (rd1 – S6) | Fibrils – High<br>1 µM / 0.5 µM | 58 | 975 | 1.23 | Ribbons: +<br>Fibrils: +<br>Fibrils-91: +<br>Ab control: - | - |
| | | | Fibrils – Dim<br>1 µM / 0.5 µM | 58 | 1266 | 1.34 | | |
| | | | Ribbons – High<br>1 µM / 0.5 µM | 1 | 918 | 2.96 | | |
| **10** | **AS-3105** | **Pol 2b<br>rd2 (rd1 – S7)** | **Single colony PCR** | **-** | **-** | **-** | **Ribbons: +<br>Fibrils: +<br>Fibrils-91: ++<br>Ab control: -** | **- KO mutant<br>-Scaffold-only** |
| 11 | AS-3611 | Pol 2b<br>rd2 (rd1 – S8) | Fibrils – Dim<br>1 µM / 0.5 µM | 3 | 2644 | 2.95 | Ribbons: ++<br>Fibrils: +<br>Fibrils-91: +<br>Ab control: - | - |

| | | | | | | | | |
|---|---|---|---|---|---|---|---|---|
| 12 | AS-3939 | Pol 2b<br>rd2 (rd1 – S8) | Fibrils – Dim<br>1 µM / 0.5 µM | 12 | 510 | 1.63 | Ribbons: ++<br>Fibrils: ++<br>Fibrils-91: ++<br>Ab control: - | - |
| **13** | **AS-4130** | **Pol 2b<br>rd1 – S8** | **Fibrils – High<br>1 µM / 0.5 µM** | **75** | **25671** | **2.53** | **Ribbons: +++<br>Fibrils: +++<br>Fibrils-91: +++<br>Ab control: -** | **-Scaffold-only** |
| | | | **Fibrils – Dim<br>1 µM / 0.5 µM** | **75** | **4450** | **1.77** | | |
| | | | **Ribbons – High<br>1 µM / 0.5 µM** | **2** | **4650** | **3.37** | | |
| 14 | AS-4326 | Pol 2b<br>rd2 (rd1 – S8) | Single colony PCR | - | - | - | Ribbons: -<br>Fibrils: -<br>Fibrils-91: -<br>Ab control: - | - |
| 15 | AS-4332 | Pol 2b<br>rd2 (rd1 – S8) | Fibrils – Dim<br>1 µM / 0.5 µM | 66 | 1279 | 1.29 | Ribbons: +++<br>Fibrils: ++<br>Fibrils-91: +++<br>Ab control: - | -Scaffold-only |
| | | | Ribbons – High<br>1 µM / 0.5 µM | 1 | 938 | 2.97 | | |
| 16 | AS-4356 | Pol 2b<br>rd2 (rd1 – S8) | Single colony PCR | - | - | - | Ribbons: +<br>Fibrils: +<br>Fibrils-91: +<br>Ab control: + | - |
| 17 | AS-4699 | Pol 2b<br>rd2 (rd1 – S8) | Fibrils – Dim<br>1 µM / 0.5 µM | 180 | 2516 | 1.15 | Ribbons: ++<br>Fibrils: ++<br>Fibrils-91: +<br>Ab control: - | - |
| | | | Ribbons – High<br>1 µM / 0.5 µM | 1 | 1246 | 3.09 | | |
| 18 | AS-4755 | Pol 2b<br>rd2 (rd1 – S8) | Ribbons – Dim<br>1 µM / 0.5 µM | 670 | 10189 | 1.18 | Ribbons: -<br>Fibrils: -<br>Fibrils-91: -<br>Ab control: - | - |
| 19 | AS-4763 | Pol 2b<br>rd2 (rd1 – S8) | Ribbons – Dim<br>1 µM / 0.5 µM | 86 | 1736 | 1.31 | Ribbons: +<br>Fibrils: -<br>Fibrils-91: -<br>Ab control: - | - |
| 20 | AS-4816 | Pol 2b<br>rd1 – S8 | Single colony PCR | - | - | - | Ribbons: ++<br>Fibrils: ++<br>Fibrils-91: ++<br>Ab control: + | - |
| 21 | AS-5024 | Pol 2b<br>rd1 – S8 | Fibrils – High<br>1 µM / 0.5 µM | 5 | 625 | 2.09 | Ribbons: +++<br>Fibrils: +++<br>Fibrils-91: +++<br>Ab control: ++ | - |
| | | | Fibrils – Dim<br>1 µM / 0.5 µM | 5 | 2947 | 2.77 | | |
| | | | Ribbons – High<br>1 µM / 0.5 µM | 1 | 3084 | 3.49 | | |
| 22 | AS-5028 | Pol 2b<br>rd2 (rd1 – S8) | Fibrils – Dim<br>1 µM / 0.5 µM | 51 | 924 | 1.26 | Ribbons: +<br>Fibrils: +<br>Fibrils-91: +<br>Ab control: - | - |
| 23 | AS-5171 | Pol 2b<br>rd2 (rd1 – S9) | Fibrils – Dim<br>1 µM / 0.5 µM | 3 | 1117 | 2.57 | Ribbons: +<br>Fibrils: -<br>Fibrils-91: -<br>Ab control: - | - |
| | | | Ribbons – High<br>1 µM / 0.5 µM | 1 | 600 | 2.78 | | |
| 24 | AS-5424 | Pol 2b<br>rd2 (rd1 – S9) | Ribbons – High<br>1 µM / 0.5 µM | 1 | 1042 | 3.02 | Ribbons: +++<br>Fibrils: ++<br>Fibrils-91: ++<br>Ab control: + | - |
| **25** | **AS-5776** | **Pol 2b<br>rd2 (rd1 – S9)** | **Ribbons – High<br>1 µM / 0.5 µM** | **1** | **1426** | **3.15** | **Ribbons: ++<br>Fibrils: +<br>Fibrils-91: +<br>Ab control: -** | **- KO mutant** |

**Supplementary Table S3 | Rosetta scores for experimentally validated designs**

Rosetta scores for designs expressed as GFP-fusion proteins, assessed for binding to different aSyn fibrils using FIDA. Designs targeted polymorph 2b and resulted from multiple design strategies, with equal representation of rd1 and rd2 optimization rounds. For rd2 designs, the preceding rd1 strategy is indicated in the brackets. All tested designs met computational thresholds. The identified aSyn pan-binder (AS-0763) and polymorph-specific aSyn Ribbons binder (AS-2777) are highlighted in bold.

| Strategy name | Design name | ddG (rigid body docking) | ddG (with side chain repack) | Shape complementarity | Interface H-bonds | Buried unsat. polar atoms |
|---|---|---|---|---|---|---|
| **rd1 – S1** | **AS-0763** | **-71.5** | **-53.4** | **0.68** | **10** | **4** |
| rd1 – S2 | AS-1210 | -68.7 | -53.3 | 0.74 | 6 | 3 |
| rd1 – S8 | AS-4130 | -88.6 | -69.9 | 0.69 | 12 | 4 |
| **rd2 (rd1 – S6)** | **AS-2777** | **-53.9** | **-36.8** | **0.75** | **9** | **4** |
| rd2 (rd1 – S7) | AS-3105 | -55.9 | -33.1 | 0.72 | 8 | 3 |
| rd2 (rd1 – S9) | AS-5776 | -56.9 | -33.3 | 0.69 | 11 | 1 |
